# Kinetic Modeling of Target-Amplification-Free CRISPR-Cas-Based Autocatalysis Reactions

**DOI:** 10.64898/2026.03.03.709462

**Authors:** Matthew Wester, Jongwon Lim, An Bao Van, Katherine Koprowski, Enrique Valera, Rashid Bashir

**Affiliations:** Department of Bioengineering, University of Illinois Urbana-Champaign, Urbana, IL 61801, USA; Nick Holonyak Jr. Micro and Nanotechnology Laboratory, University of Illinois Urbana-Champaign, Urbana, IL 61801, USA; Materials Research Laboratory, University of Illinois Urbana-Champaign, Urbana, IL 61801, USA; VinUni-Illinois Smart Health Center, VinUni Campus, Hanoi 100000, Vietnam; Carl R. Woese Institute for Genomic Biology, University of Illinois at Urbana-Champaign, Urbana, Illinois 61801, USA; Department of Mechanical Science and Engineering, University of Illinois at Urbana-Champaign, Urbana, IL 61801, USA; Department of Electrical and Computer Engineering, University of Illinois at Urbana-Champaign, Urbana, IL 61801, USA; Department of Materials Science and Engineering, University of Illinois at Urbana-Champaign, Urbana, IL 61801, USA; Department of Biomedical and Translation Science, Carle Illinois College of Medicine, University of Illinois at Urbana-Champaign, Urbana, IL 61801, USA; Chan Zuckerberg Biohub Chicago, Chicago, IL 60642, USA

## Abstract

CRISPR-Cas-based diagnostics utilize the Cas enzyme’s trans-cleavage activity to generate signal and have become popular platforms for sensitive nucleic acid detection. Recently, autocatalytic systems have been demonstrated to improve the time to response and sensitivity in some cases. However, mechanistic description of these assays is limited and optimization relies on simple trial-and-error. In this work, we present the first comprehensive kinetic model that integrates all major biochemical processes involved in these assays, including cleavage reactions, nucleic acid equilibrium kinetics, inhibition of trans-cleavage by single-stranded DNA, and degradation of single-stranded reaction components. We discuss the biochemical foundations and implementation of the ordinary differential equation model, which is built for adaptation to different reaction schemes. We use the full model to investigate the role of nucleic acid stability in assay performance for a typical nucleic acid design and show that our model demonstrates inhibition effects consistent with experimental data. We describe the reaction behavior, derive a simplified analytical model and compare its performance to the full analytical model. Finally, we demonstrate tools developed for rapid *in silico* optimization to guide the rational design of future target-amplification-free CRISPR-Cas-based autocatalysis assays.

## Introduction

Since the discovery of the trans-cleavage activity of some class 2 clustered regularly interspersed palindromic repeat-associated (CRISPR-Cas) enzymes, they have held promise for rapid diagnostic applications.^[1]^ CRISPR-Cas enzymes work by recognizing a 20-30 nucleotide region of DNA or RNA based on complementarity to a provided guide or CRISPR RNA sequence (gRNA or crRNA, respectively); upon successful target recognition, cleavage of the nucleic acid occurs via a cis-cleavage mechanism.^[2]^ In certain Cas types, including Cas12 and Cas13, this triggers a conformational change that enables the enzyme to indiscriminately cleave single-stranded DNA and RNA (ssDNA and ssRNA). Single-stranded nucleic acid molecules designed with a fluorophore and quencher pair can then be cleaved, leading to a signal increase as the fluorophore-quencher pairs separate.^[3]^ This principle was first demonstrated with the Specific High-Sensitivity Enzymatic Reporter UnLOCKing (SHERLOCK) assay, which used Cas13 to detect genome fragments of viruses and bacteria with single-base discrimination capabilities^[4]^ To achieve high sensitivity, the SHERLOCK assay used a recombinase polymerase amplification (RPA) pre-amplification step to enrich the target sequence before conversion to target RNA using T7 polymerase. This allowed for the detection of concentrations as low as approximately 2 aM, but with two hours required for RPA and 1-3 hours for CRISPR-Cas-based detection. This type of assay was expanded to Cas12 with DNA Endonuclease-Targeted CRISPR Trans Reporter (DETECTR), which similarly used an RPA target pre-amplification step.^[5]^ This assay similarly had an attomolar limit of detection but greater than an hour assay time.

Many subsequent assays utilizing pre-amplification followed by a CRISPR-Cas-based detection step have demonstrated similar performance across a variety of targets.^[6]^ While these achieve high sensitivity, the prolonged assay time limits their application as rapid diagnostic tests. Further, the use of multi-step processes increases assay complexity. While various approaches such as phase separation and integrated microfluidic devices have been employed to mitigate the latter issues, these approaches still rely on pre-amplification of the target nucleic acid and the associated time delay.^[7,8]^ Alternatively, single CRISPR-Cas-based assays that lack the pre-amplification step can provide more rapid results but with the practical limit of detection around 1 pM.^[9]^ To improve the assay time while maintaining high sensitivity, an emerging approach involves the implementation of positive feedback systems; these focus on signal amplification rather than direct amplification of the target analyte (Supplementary Table 1).^[10–24]^ Among these, fully CRISPR-Cas-based signal amplification systems are attractive for their fast time to result and low complexity, while maintaining the high specificity and target multiplexing capabilities characteristic of all CRISPR-Cas-based systems.

These autocatalytic systems work by introducing an engineered nucleic acid molecule containing a second distinct target as well as one or more trans-cleavage sites. Upon trans-cleavage of this molecule by the activated enzyme, a cascade is initiated wherein the target strand is exposed to and activates a second Cas enzyme, which drives further trans-cleavage. We refer to this molecule as the autocatalytic nucleic acid (ANA) because it is responsible for initiating the autocatalytic phase of the reaction. The first demonstration of such a system, a CRISPR-Cas-only amplification network (CONAN), utilized a switchable caged gRNA (scgRNA) containing a trans-cleavage site flanked by fluorophore and quencher molecules.^[11]^ Upon cleavage of the scgRNA, the fluorophore-quencher pair dissociated and the released gRNA activated the second enzyme (along with its DNA target, which was provided in solution), leading to autocatalytic activation. This assay achieved a 5 aM limit of detection in four hours, on the same time scale as CRISPR-Cas assays with pre-amplification. A subsequent demonstration by Zhang et al., brought the reaction time down to 20 minutes with a reported 1 aM limit of detection.^[12]^ This system detected target RNA with Cas13 and used Cas12 to drive the autocatalytic phase. The primary ANA was composed of a target DNA blocked by complementary RNA containing a cleavage site flanked by locked nucleic acid (LNA) bases to enhance its stability. As in most demonstrated CRISPR-Cas-based assays, a distinct reporter molecule was used.

Rather than the two-component ANAs used in these demonstrations, subsequent publications have favored one-component ANAs in which the target region is blocked by the nucleic acid secondary structure. This includes the system by Deng at al., which used a circularized double-stranded DNA to achieve a 1 aM limit of detection in 15 minutes.^[14]^ The most popular design is a hairpin structure, with some variant of this structure used in at least six CRISPR-Cas-only autocatalytic systems to date.^[15,17–21]^ A DNA hairpin is the simplest design to achieve an ANA because it is composed of a blocked DNA sequence and a cleavable single-stranded region. Beyond their simplicity, these structures may be chosen because they have been studied extensively and used in existing biosensors.^[25–29]^

Among the assays demonstrated, there are significant differences in the design and performance. Further, there is little discussion of the theoretical basis for these assays to guide rational design and optimization. In this paper, we aim to provide the first computational model integrating all major mechanisms present in CRISPR-Cas-based autocatalytic reactions. We describe these mechanisms and their implementation in a system of ordinary differential equations (ODEs) that is easily adaptable to different reaction mechanisms described in literature. We specifically highlight the modeling of ANA dynamics and the competitive cleavage of ssDNA reaction components. We demonstrate that this model can replicate the characteristics of assays in existing literature. We derive a simplified analytical model and test its agreement against the full numerical model. Finally, we develop and demonstrate simple, *in silico* modeling tools for optimization of CRISPR-Cas-based autocatalytic assays.

## Results and Discussion

### CRISPR-Cas-Based Autocatalysis Reactions

To model a typical CRISPR-Cas-based autocatalytic system, we first define the core reactions driving signal production. Although specific implementations vary between previous assays, the individual reactions and overall scheme remain similar. For brevity, the descriptions in this manuscript are generalized for Cas12, but the system can be applied to Cas13 with minimal changes. The reaction can be conceptualized in two stages (Fig. 1a). In the target detection stage, the target strand of DNA is detected by the first Cas ribonucleoprotein (RNP) complex, initiating the trans-cleavage process. In the signal amplification stage, the ANA is cleaved and the ANA target strand is recognized by the second RNP complex. Each of these stages increases the concentration of active enzyme that can further trans-cleave the reporter and ANA. A more in-depth reaction scheme is shown in Figure 1b, with reactions color-coded according to their types – Michaelis-Menten, reversible, or irreversible. The full reaction system is detailed in Supplementary Note 1.

**Figure 1.**
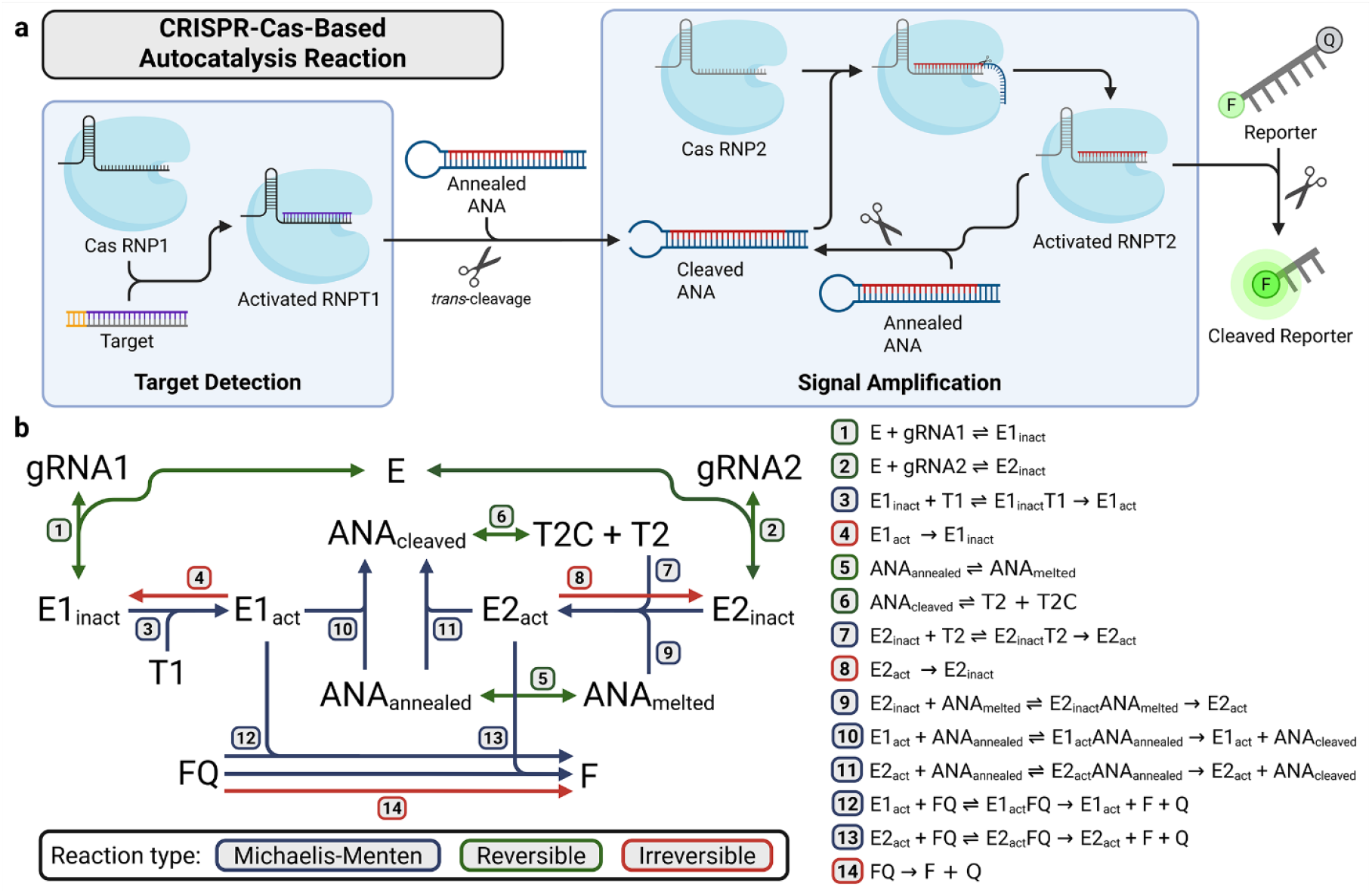
CRISPR-Cas-based autocatalysis reaction schematics. a) Simplified autocatalysis scheme. The target detection stage begins when the target nucleic acid is recognized by the ribonucleoprotein (RNP) complex RNPT1. This results in trans-cleavage of the autocatalytic nucleic acid (ANA). The trans-cleaved ANA activates RNPT2, which cleaves additional ANA. ANA cleavage initiates the signal amplification stage. Activated RNP complexes then cleave the reporter molecule. b) Detailed reaction schematic of the CRISPR autocatalysis reaction, showing the complex interactions between various species. Reactions are colored according to their classification as Michaelis-Menten, reversible, or irreversible. The full reaction scheme is detailed in Supplementary Note 1.

The reaction scheme includes several categories, with the selection of reaction rate constants detailed in Supplementary Note 1 and summarized here. Briefly, RNP complex formation (reactions 1 and 2) progresses through a tight binding step and a cleavage step, with binding occurring on the time scale or seconds, unbinding in days, and cleavage in minutes.^[30]^ Because the RNP is often pre-complexed, creating mature gRNA that does not undergo later cleavage during binding, this reaction is modeled as a simple irreversible reaction. Cis-cleavage (reactions 3, 7, and 9) converts the RNP into its active state and can be modeled as a Michaelis-Menten process.^[31]^ Cis-cleavage is ignored in analysis of single CRISPR-Cas based reactions because it occurs at the beginning of the reaction on the time scale of seconds.^[32]^ However, it is significant in autocatalytic reactions because cis-cleavage of the ANA-derived target continues throughout the reaction, thereby affecting the rate of autocatalysis. Trans-cleavage (reactions 10-13) is the mechanism for ANA cleavage and signal production, so it is central to the reaction. The trans-cleavage behavior of Cas enzymes has been characterized extensively for single CRISPR-Cas-based assays and can be described according to its catalytic efficiency (*k*_*cat*_/*K*_*M*_).^[9,32,33]^ This is also modeled as a Michaelis-Menten process, and the parameters can be measured through a fairly simple experiment (Fig. S1, Supplementary Note 2). We also consider enzyme inactivation (reactions 4 and 8) as an irreversible process, although this reaction is very slow and has little effect on the time scale of autocatalytic reactions.^[31]^ We model non-specific degradation of the reporter (reaction 14) as a slow but constant, irreversible process.^[34]^ Finally, we model the behavior of the uncleaved and cleaved ANA (reactions 5 and 6) as a two-state equilibrium detailed in Supplementary Note 3. The rate constants for these reactions are derived from the standard Gibbs free energy change between the two states and the time scale for equilibrium to be established after perturbation, which is on the order of microseconds to milliseconds.^[28,35]^ This type of equilibrium should reasonably capture the behavior of a hairpin.^[36]^ This is discussed in more depth in a subsequent section.

### Model Setup and Operation

With the reactions defined, the simulation of the CRISPR-Cas-based autocatalytic system is based on solving a system of ODEs to obtain the concentration of each of the species involved over a certain time range. For simple systems, the ordinary equations can be written by hand and input directly into an ODE solver. However, this approach is difficult to scale to systems such as the autocatalytic one, with the described system composed of 24 reactions and 33 species if the tracking of ssDNA fragments is disabled. Further, the manual approach does not readily support testing different reaction schemes. With this in mind, we developed the software for our model to readily support changes to the reaction scheme and modification of constants. The model setup is detailed in Supplementary Note 4. Briefly, each autocatalytic system is defined in its own function based on a plain text list of reactions. After defining default rate constants and other options, helper functions create an ODE system object that can be solved with built-in MATLAB ODE solvers. Additional helper functions are provided to set up the reaction and set the initial concentrations. Finally, several functions are provided to prepare and run experiments, plot their outcomes, and select data from the experiments for custom analysis. The software is intended to provide a user-friendly option for modeling these systems while maintaining the ability of the user to design custom experiments.

The standard experiment to demonstrate the model starts by defining the concentration of species involved in RNP complex formation and ANA annealing in three tubes; the incubation steps for these tubes are then individually simulated to obtain the final concentration (Fig. 2a). The function implementing the incubation steps serves as a template for creating similar protocols. The user then specifies the concentration of any species added to the reaction (such as the reporter and target) and the dilution of each of the pre-incubation tubes and the initial concentrations are prepared by another function (Fig. 2b). The user then selects the ODE function corresponding to the desired reaction system and specifies any constants to be modified from their default values. The ODE function and the initial conditions are passed to a built-in MATLAB solver, which returns the time-series concentration data for each species. This data can be passed to a plotting function that produces a summary of the reaction results and several additional figures tracking the concentration of individual species (Fig. 2c, Fig. S2).

**Figure 2.**
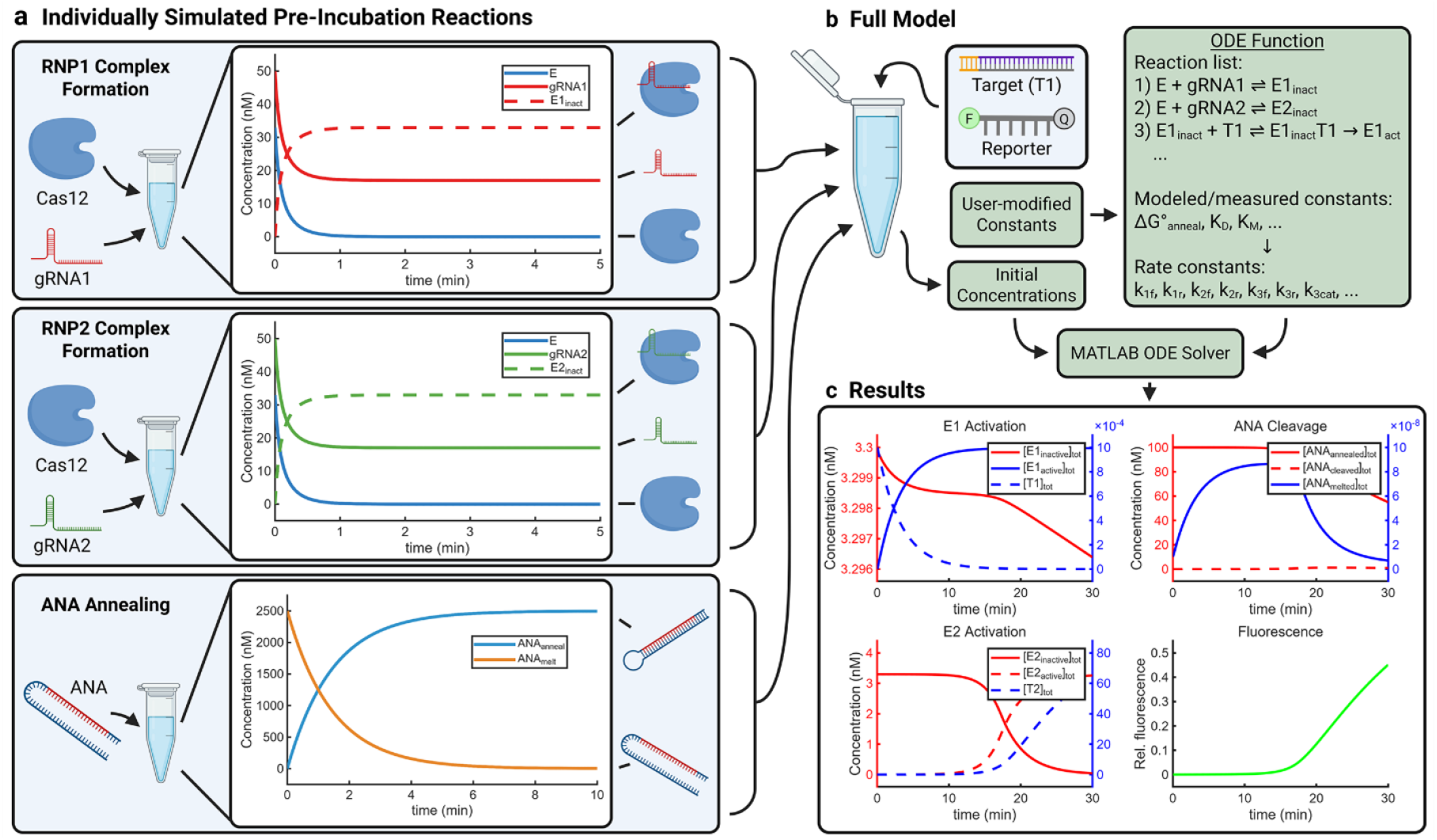
Typical CRISPR-Cas-based autocatalysis kinetic model workflow. a) The developed software enables *in silico* modeling of specific reaction steps, including distinct pre-incubation and annealing. Data is shown for RNP1 and RNP2 complex formation and ANA annealing. b) Following pre-incubation, the tubes are combined and additional reagents are included to complete the initial species concentrations for the full simulation. An ordinary differential equation (ODE) function is programmatically generated from a list of reactions and modeled or measured thermodynamic and kinetic parameters. The ODE function and initial conditions are used as inputs for a built-in MATLAB ODE solver, which generates time-series data. c) The results are plotted to show distinct stages of the reaction, including enzyme activation, ANA cleavage, and fluorescence signal increase.

The results in Figure 2c reflect the key processes involved in a typical CRISPR-Cas-based autocatalysis reaction. This starts with E1 activation, which occurs on the time scale of seconds to minutes, in line with the cis-cleavage rates. This does not significantly increase the signal, as the concentration of active enzyme is approximately equal to the assay target concentration (1 pM). However, this enzyme slowly begins to cleave the ANA, which dissociates and activates E2. This process can be seen to accelerate around 10 minutes into the reaction as the autocatalytic mechanism begins to dominate. This results in an exponential increase in fluorescence. We can also evaluate the performance of the model under control conditions to confirm it aligns with the expected behavior (Fig. S3). It is observed that when the second enzyme is removed, the reaction performs as a single CRISPR-Cas assay, where the positive sample exhibits a fluorescence curve typical of Michaelis-Menten kinetics over a long time scale. These control experiments also verify the presence of delayed background amplification for a negative sample, consistent with the performance of previously published CRISPR-Cas-based autocatalytic assays.

### Modeling of Autocatalytic Nucleic Acid

For demonstration of this kinetic model and its capabilities, we utilize a hairpin ANA design. In addition to being the most common structure, hairpins are simple to model because they can be reasonably described by a two-state equilibrium (Fig. 3a).^[36]^ Further, while the stoichiometry of multi-strand ANAs must be optimized, hairpins have a fixed stoichiometry. In a system based on the hairpin ANA, a two-state equilibrium is established between the annealed and melted forms. Trans-cleavage by the activated Cas enzyme results in the cleaved form, which is in equilibrium with the dissociated target and complement strands. In this model, the target is the primary species that drives signal amplification while the melted ANA is the primary species that drives noise. Therefore, these equilibria are key to assay optimization.

**Figure 3.**
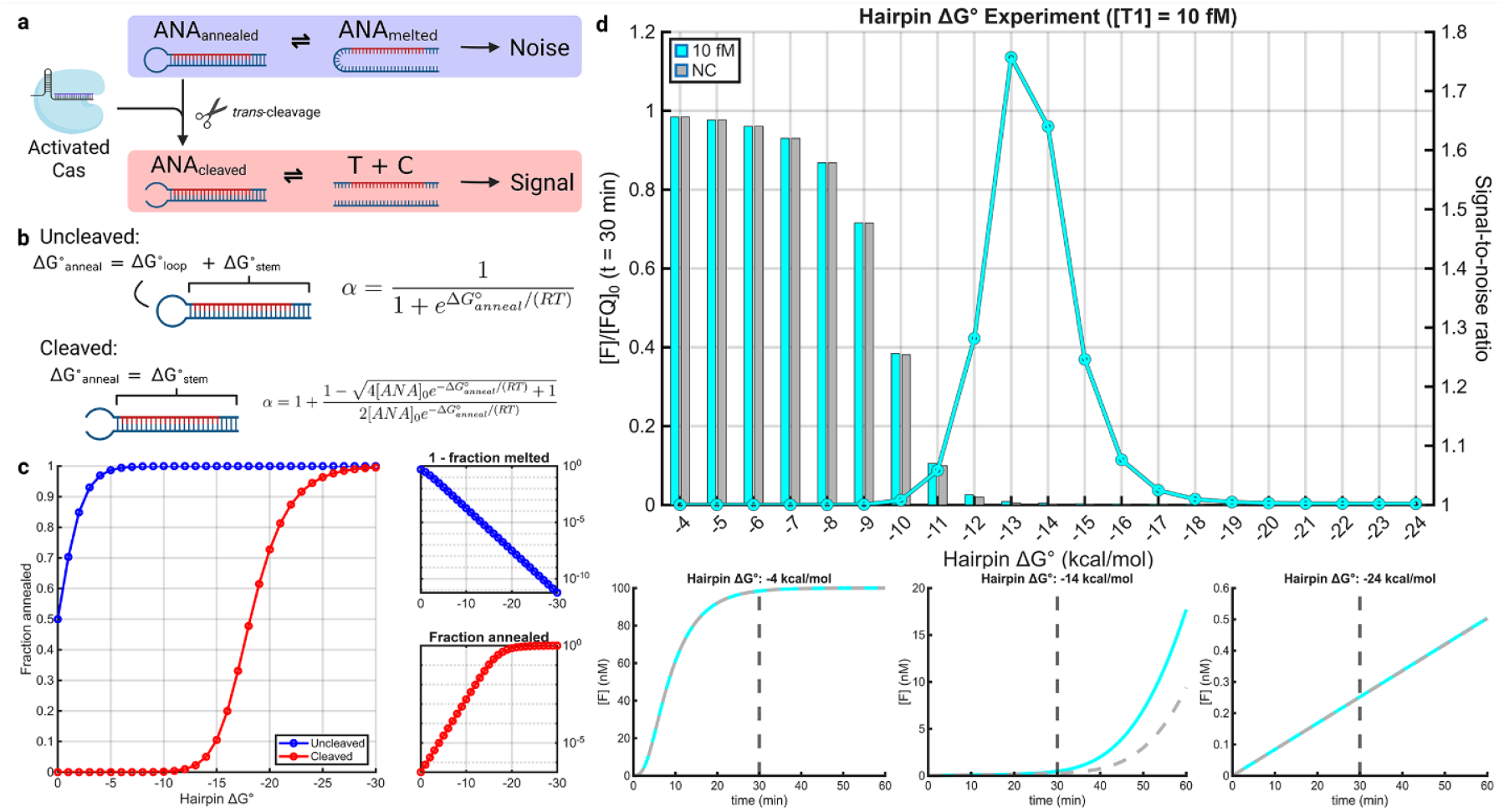
Variation of annealing free energy in a hairpin model. a) Schematic of two-state model used for uncleaved and cleaved ANA. Uncleaved ANA is in equilibrium between melted and annealed states while cleaved ANA is in equalibirium with its target and complement strands. b) Model of secondary structure free energy before and after cleavage, with corresponding equations for *α*. 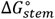 varies depending on the stem design while 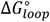 is approximately 4 kcal/mol for a DNA hairpin. c) Fraction of annealed ANA before and after cleavage as a function of hairpin 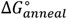) Autocatalytic amplification results as a function of hairpin 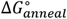, showing how the reaction characteristics such as signal-to-noise ratio depend on the ANA free energy.

In the hairpin model, the free energy of annealing for the uncleaved molecule can be considered as the sum of the free energy for the stem and the loop, with intact loops generally destabilizing the molecule (Fig. 3b).^[28]^ The exact free energy penalty depends on the loop length and buffer conditions, but is approximately 4 kcal/mol in ssDNA for the loop length range utilized in previous ANA designs. In practice, the free energy of the uncleaved hairpin can be adjusted easily by changing the stem length. After cleavage, we consider the free energy of annealing as the free energy for the stem with the loop-associated penalty removed. This results in a decrease in the free energy after cleavage, with the resulting molecule more stable than before. This approach aligns with the results of NUPACK simulations for both hairpin and bulge molecules (Fig. S4). However, the duplex molecule has an additional concentration-dependent entropic penalty related to the loss of translational freedom during molecular association that was not present in the initial hairpin.^[37]^ Thus, while the fraction of ANA that is annealed is intrinsic to the reaction, the fraction of cleaved ANA that remains associated depends on the strand concentration (Supplementary Note 5). Consequently, the prevalence of the signal- and noise-generating species cannot be directly compared based on free energy values. Instead, the parameter *α* is introduced to describe the fraction of ANA in the blocked state as a function of free energy and concentration.

Calculating *α* for each molecule as a function of the stem free energy (with the loop penalty fixed at 4 kcal/mol) demonstrates the relationship between ANA stability and the fraction annealed in each state (Fig. 3c). When the hairpin Δ*GG*^∘^ is zero, approximately 50% of the uncleaved hairpin is melted at equilibrium, while the cleaved hairpin is nearly 100% melted. As the hairpin Δ*G*^∘^ becomes more negative, the annealed fraction of the uncleaved ANA exponentially approaches one while the annealed fraction of the cleaved ANA approaches one at a reduced exponential rate. This results in a free energy window where the uncleaved hairpin is almost completely annealed and the cleaved hairpin is nearly completely melted, which should be optimal in terms of maximizing the signal-to-noise ratio (SNR). However, it is unclear where exactly in this 10-15 kcal/mol window the SNR will be maximized.

The full kinetic model can be applied to explore this phenomenon in more detail (Fig. 3d, Fig. S5). At a high hairpin Δ*G*^∘^, both the signal and noise are high and so the SNR is poor (inset graph one). As the hairpin is stabilized, the signal begins to outpace the noise (inset graph two), resulting in a maximum SNR of ~1.76 for a 10 fM concentration. At 60 minutes, the SNR increases to ~2.93 (Fig. S6). Further reduction in the hairpin Δ*G*^∘^ suppresses the autocatalytic function entirely, and the system functions as a single CRISPR-Cas reaction (inset graph three). At a concentration below the LOD for a single CRISPR-Cas reaction, this reduces the SNR to one. However, when the target concentration is above the LOD for these systems (e.g. 100 pM), we observe that the SNR approaches a stable positive value (Fig. S7). Thus, while decreasing the Δ*GG*^∘^ of the ANA results in a more stable molecule, it can also reduce signal to the extent that autocatalysis is entirely inhibited. The results of this experiment support the optimization of the Δ*GG*^∘^ for the annealed ANA as a critical component of assay design. In addition to changing the structure length, approaches like adding LNA can modify this value.^[12]^

While hairpin designs are common, a variety of ANA structures have been demonstrated in previous literature.^[11–24]^ These include two-strand molecules with bulge structures and more complex designs like circular DNA nanostructures (Supplementary Table 1). For ANA designs with more complex structures, the cleavage process involves additional complexity (Fig. S8). The design and analysis of these molecules is typically limited to calculation of the minimum free energy (MFE) secondary structure by nucleic acid analysis software such as NUPACK.^[38]^ In practice, the free energy of simple uncleaved structures can be estimated from melting curve experiments, which show good agreement for hairpins and bulges (Figs. S9 and S10). However, the process of ANA cleavage and conversion to T2 is more complex than is reflected in this type of analysis. First, additional secondary structures exist in equilibrium with the MFE structure. Some of these may have different cleavage sites or partially exposed target strands. It is not obvious which secondary structures may activate the Cas enzyme, although the free energy change during crRNA binding may predict Cas activation.^[39]^ After cleavage, the ANA is typically presumed to dissociate and expose the target strand, however, this ignores the interaction of the cleavage products with other nucleic acids. Notably, bulge ANAs such as those demonstrated by Zhang et al. have an additional variable to consider because the target and complement can be introduced at different ratios.^[12]^ Despite this complexity, it is likely that the results of this analysis can be applied to other ANA designs. Notably, bulge features have a similarly destabilizing effect to hairpins, so the model here could be readily adapted to these.^[40]^ The majority of designs in previous literature are variants of either a hairpin or bulge design, so it is likely that this analysis broadly applies.

### Modeling of Background ssDNA Cleavage

The cleavage of ssDNA is the basis for the core functions of CRISPR-Cas-based autocatalytic reactions. However, any single-stranded nucleic acids present in solution are also expected to undergo trans-cleavage.^[41,42]^ Other than a reported moderate preference for certain bases, the targeting of ssDNA for trans-cleavage appears to be random.^[42,43]^ Thus any excess ssDNA sources such as excess ANA or the complementary strand of either target may present significant inhibition. Excess target beyond the amount required for enzyme activation also inhibits reporter trans-cleavage (Fig. S11). To be accurate, the autocatalytic model must capture this behavior.

This is challenging because the exact mechanism of this cleavage is unclear. Direct evidence demonstrates that cis-cleavage occurs by enzyme binding and translation along the target until reaching the recognized sequence.^[44]^ However, similarly strong evidence does not exist for trans-cleavage, to the authors’ knowledge. Li et al. reported that the trans-cleavage products were present with a length of 2 or 4 nucleotides.^[41]^ These fragments appeared within minutes of reaction initiation, indicating that they were not the result of progressive degradation steps; this may indicate that the Cas enzyme tends to cleave fragments from the end of the strand. Sun et al. found that the addition of LNA at specific locations near the 3’ end of ssDNA could direct site-specific cleavage, consistent with a model in which the Cas enzyme binds near the 3’ end and moves in the 5’ direction.^[15]^ This mechanism is also indicated in dsDNA with a 3’ toehold.^[45]^ However, a clear mechanism that constrains trans-cleavage to the end of the strand has not been demonstrated. Regardless of the mechanism, it is likely that ssDNA strands undergo multiple cleavage events during the degradation process, increasing the inhibitory effect beyond what would be expected from their initial concentration.

Figure 4a shows the cleavage products generated from a two-component ANA (pictured) in a gel electrophoresis experiment (Supplementary Note 6). The distribution of cleavage products seen after one hour of cleavage is beyond what would be expected due to slight variation of the initial trans-cleavage location, consistent with our previous work.^[18]^ The small ssDNA fragments generated are likely not stained well by the SYBR dye used in this assay. Only a trace of the cleavage products can be observed at two hours, with the products appearing completely digested after three hours indicating complete degradation of these intermediates. To assess the kinetic effects of excess ssDNA directly, we designed an inhibition experiment. This involves conducting multiple parallel Michaelis-Menten experiments with varying concentration of inhibitor (Fig. S12, Supplementary Note 2). In this case, the inhibitor was a 31-nt ssDNA molecule unrelated to any other assay components; this length is in the typical length range for ANA components released during the reaction. Increasing the concentration of the inhibitor results in a reduction in the reporter cleavage rate (Fig. 4b). This relationship can be visualized in a Lineweaver-Burk plot, which shows results consistent with competitive inhibition (Fig. 4c). This aligns with the assumption that background ssDNA competes for the catalytic site of the Cas enzyme. The data presented here indicate an inhibition constant of approximately ~196 nM, indicating a moderately strong inhibitor. This value should be interpreted cautiously, as it is based on a standard equation that assumes the inhibitor is not degraded.

**Figure 4.**
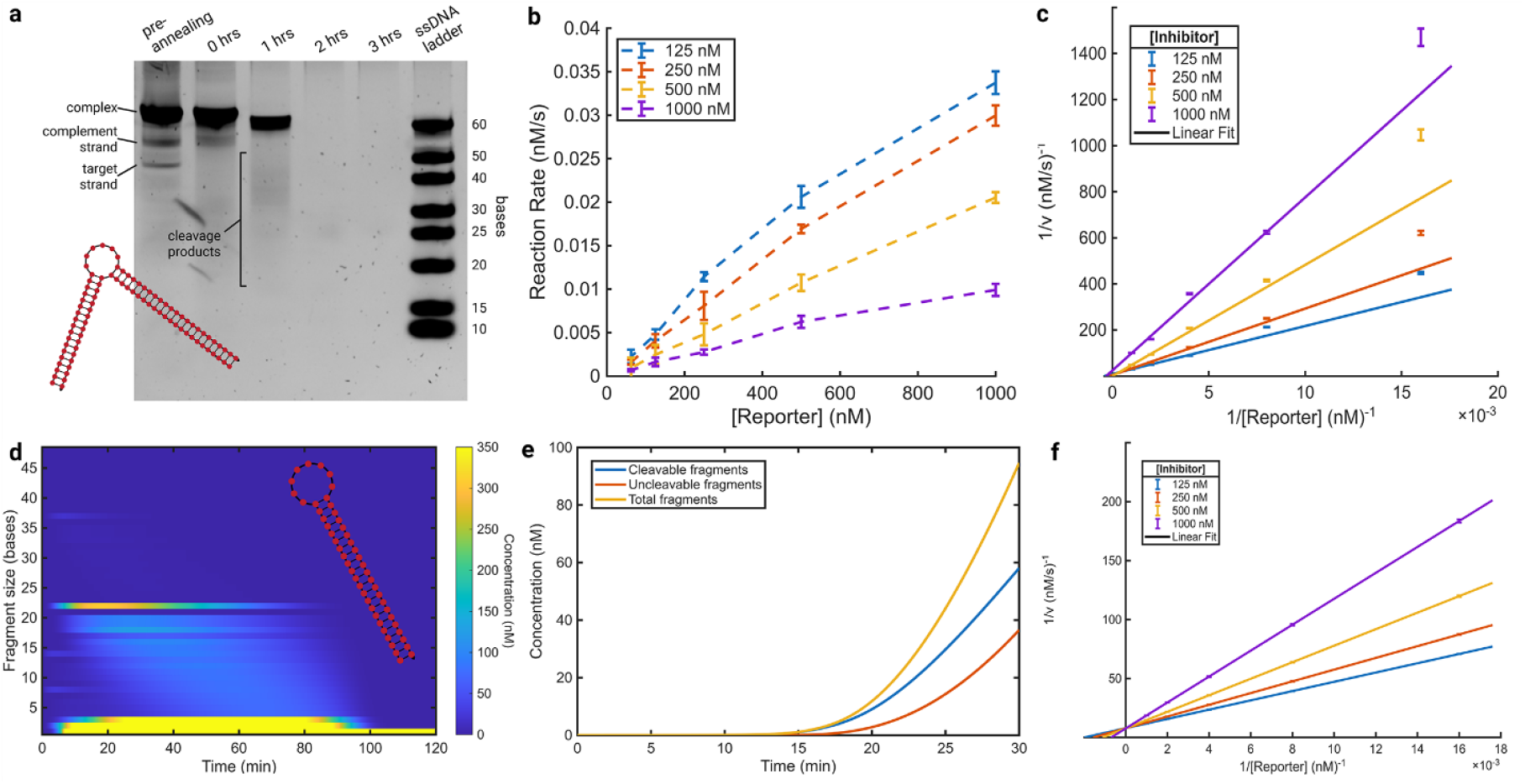
Modeling reaction inhibition by ssDNA fragmentation. a) Polyacrylamide gel electrophoresis experiment showing degradation of cleavage products on the time scale of approximately one hour. b) Inhibition experiment results plotting reaction rate as a function of single-stranded DNA (ssDNA) inhibitor concentration. Error bars indicate the mean plus or minus the standard error (n=4). c) A Lineweaver-Burk plot of the experiment in (b) shows increased inhibition resulting from increased ssDNA concentration. Both the data and standard error were transformed and plotted along with a linear fit to the data. d) Simulated DNA fragment distribution resulting from an *in silico* experiment modeled after the gel conditions in (a) for the hairpin model. e) Total concentration of ssDNA fragments over time, separated based on whether they are cleavable and uncleavable due to complete degradation. f) Lineweaver-Burk plot generated from simulated data corresponding to the experiment plotted in (b) and (c), demonstrating general agreement between the measured and simulated inhibition.

To model this phenomenon efficiently in our MATLAB model, a vector-based process for considering DNA fragmentation by the Cas enzyme was developed. This process is detailed in Supplementary Note 7. It tracks the concentration of fragments of different lengths in a vector, which allows for vector-based Michaelis-Menten kinetics:

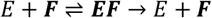

The model incorporates an affinity vector to reflect the binding kinetics of the enzyme for fragments of different lengths. Nalefski et al. noted that at least three-fold greater cleavage activity was observed against 10- or 20-nt substrates compared to 5-nt substrates.^[42]^ Based on this, the model includes option for length-dependent modeling as well as the default length-independent binding. The model also requires a fragmentation matrix that expresses the probability for fragments of each length to be trans-cleaved into fragments of every other length. This can be configured for cleavage at the end of the strand or random cleavage.

Using this process, the model predicts the production of DNA fragments over the course of experiment. Figure 4d shows an *in silico* recreation of a CRISPR-Cas-based autocatalysis experiment using the hairpin model. The concentrations for this experiment (1 μM final ANA concentration and ~50 nM active Cas enzyme) are typical of those used for visualizing cleavage products. The model is configured for length-independent binding and cleavage between 2-5 nucleotides from the strand end. Identifiable bands corresponding to cis-cleavage products can be seen to appear almost immediately. The approximate lengths of these products are described in Supplementary Note 1. As the experiment proceeds, small fragments accumulate at a high concentration while the prominent cleavage products slowly decrease in length. We hypothesize that this produces the smeared bands visible in gels analyzing cleavage products. ssDNA fragments are nearly completely broken down into small fragments after two hours, consistent with our gel observations. These results do not attempt to account for the binding affinity of the gel stain, so small fragments will appear more prominent than in an actual gel. In an experiment with an ANA concentration more typical for a reaction (100 nM), the concentration of cleavable fragments can be seen to increase to nearly the total ANA concentration in the first 30 minutes, effectively doubling the concentration of inhibitors (Fig. 4e). The generation of fragments should scale with the overall enzyme activity, so the generation of these species increases exponentially. However, for short reactions typical of CRISPR-Cas-based autocatalysis assays, these fragments may not present a significant challenge.

We can use the full model with the inhibition mechanism for *in silico* experiments. For example, simulation of a single Michaelis-Menten experiment achieves fair agreement with our measured results (Fig. S13). This can be applied to the inhibition experiment as well. Modeling the exact experimental conditions from Figure 4c results in a similar Lineweaver-Burk plot, indicating agreement of the model with the experimentally measured inhibition (Figure 4f). While these plots show a different magnitude for the inverse of velocity (~1400 vs. ~200), this quantity is very sensitive to fluctuation in enzyme activity that may occur between experiments. For example, lowering *k*_*cat*_ of the enzyme trans-cleavage to 0.01 s^-1^ results in a magnitude greater than observed experimentally (Fig. S14).

In addition to background ssDNA cleavage, the model also includes cleavage of ssDNA components of the reaction, such as T2. This likely does not represent a significant source of inhibition in most assays due to the low concentration of these species, but their degradation may affect the overall performance of the assay. However, it is difficult to capture the full complexity of this process, as the exact site of cleavage may determine if the species remain functional. The gel-like visualization along with the individual species plots can help to identify problematic generation of fragments in a certain reaction scheme.

### Analysis and Optimization of CRISPR-Cas-Based Autocatalysis Assays

The full numerical model can be useful both for analysis and *in silico* optimization of system behavior. To analyze these assays, it is necessary to first qualitatively describe the fluorescence signal observed in the reactions. The characteristic curve can be split into three regions: the (i) exponential phase, (ii) linear phase, and (iii) saturation phase (Fig. 5a). Because our model can show the concentration of each species throughout the reaction, it is particularly helpful in understanding the behavior throughout each phase. As discussed previously, the exponential phase signal is driven by autocatalytic activation of E2. This phase lasts until either all enzymes are active or the activating components (such as ANA) are fully consumed. The ANA is in excess of the enzyme in about half of previously demonstrated reactions (Supplementary Table 1). At this point, the amount of active enzyme is effectively constant, so the rest of the reaction behaves similarly to a Michaelis-Menten reaction, with a constant velocity. A linear cleavage rate is maintained until the depletion of reporter leads to saturation.

**Figure 5.**
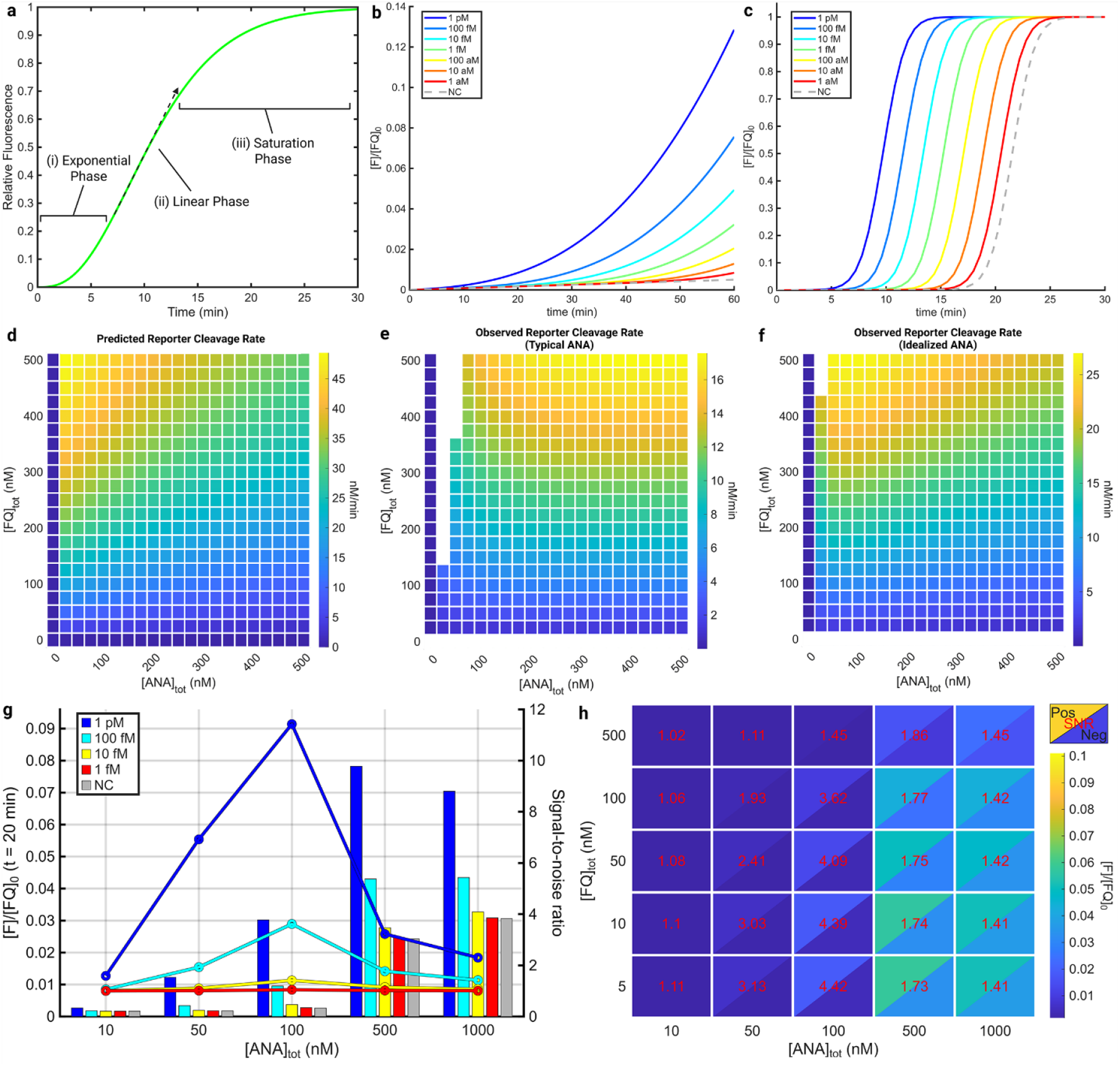
Description and optimization of CRISPR-Cas-based autocatalysis assays. a) Characteristic plot demonstrating the different phases of the autocatalytic reaction. b) Demonstration of prolonged exponential behavior in the developed model ([*E2*] ≈ 10 nM, [*ANA*] ≈ 50 nM, [*FQ*] = 100 nM, 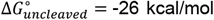, ΔΔ*G*^∘^ = 4 kcal/mol, and *k*_*cat, trans*_= 1 s^-1^). c) Demonstration of delayed sigmoidal assay response in the developed model ([E2] ≈ 10 nM, [*ANA*] ≈ 50 nM, [*FQ*] = 100 nM, 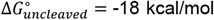, and ΔΔ*G*^∘^ = 4 kcal/mol, and *k*_*cat, trans*_ = 1 s^-1^). d) Predicted reporter cleavage rate in the linear phase of the reaction based on the simplified analytical model ([*E2*] ≈ 10 nM). e) Observed reporter cleavage rate in full numerical model with a typical ANA design (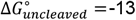 and 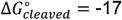). f) Observed reporter cleavage rate in the full numerical model with an idealized ANA (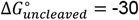 and 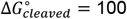). g) Optimization of reaction ANA concentration using the single variable optimization tool ([E2] ≈ 10 nM, 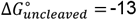 and 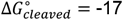). The bar graphs show the fraction of reporter cleaved (left axis) and the overlaid line graphs show the signal-to-noise ratio (right axis). h) Optimization of ANA and reporter concentration using the dual variable optimization tool under the same conditions. The plotted output reflects both the SNR and signal magnitude because the SNR is only meaningful above a detection threshold.

Previous assays generally produce output resembling one of two characteristic curves (Fig. S15). By systematically varying the pre- and post-cleavage free energy, we can observe both of these reaction modes (Fig. S16). The first observed mode occurs when the assay is maintained in the exponential phase during its entire operation, such as in the ASCas or AutoCAR assays.^[12,14]^ This behavior generally occurs in our model when the post-cleavage Δ*GG*^∘^ is very low, assuming that the uncleaved ANA is stable enough to sufficiently suppress background amplification (Figure 5b). The modeled species concentration data indicates that this occurs in assays with highly stable ANAs because the activation of E2 is extremely slow. Decreasing the stability of the cleaved ANA increases the speed at which signal amplification occurs and results in either delayed sigmoidal behavior (Figure 5c) or the background overtaking the signal, depending on the stability of the uncleaved ANA. This second mode has been observed in assays such as the split activator assay.^[15]^ While the exponential mode allows for faster results, the signal difference between sample concentrations is minimal, making quantification once regular sources of experimental variation are introduced. Further, conditions resulting in both rapid and sensitive assays are difficult to achieve. For example, the reaction in Figure 5b can distinguish 1 aM samples from the negative control; lowering the catalysis rate for trans-cleavage by a factor of ten or reducing ΔΔ*GG*^∘^ (the difference in free energy from the uncleaved to cleaved states) from 4 kcal/mol to 0 kcal/mol increases the apparent limit of detection to 1-10 fM (Fig. S17). Increasing ΔΔ*GG*^∘^ can increase the sensitivity of the assay beyond the practically observed values, but a fundamental limit between time and sensitivity exists even at arbitrarily large ΔΔ*GG*^∘^ (Fig. S18). It is unclear what value for ΔΔ*GG*^∘^ is achievable in practice, but this is likely the primary factor in determining the limit of detection.

It has previously been demonstrated that Michaelis-Menten kinetics can be reasonably applied to single CRISPR-Cas-based reactions by applying certain analytical simplifications; this can be extended to more complex predictions, such as the theoretical limit of detection for those assays.^[9,32]^ To provide a more general understanding of these assays and guide optimization, it may be reasonable to provide a similar model for autocatalytic CRISPR-Cas-based reactions. Deng et al. developed a model for their ANA and observed that their autocatalytic reaction remained exponential under conditions where reagents were not depleted, although a quantitative comparison was not made between their derived equation and the exponential fit.^[14]^ They similarly observed linear performance, but did not evaluate the agreement of the linear rate quantitatively. Therefore, it remains unclear if a simplified analytical model can reliably capture the behavior of autocatalytic reactions. Because parameters can be easily modified in our reaction, it is uniquely suited to determine whether analytical predictions based on a simplified model hold true in the full reaction.

To investigate this, we developed simplified analytical models for our reaction during the exponential and linear phases (Supplementary Notes 9 and 10). Including the full ANA equilibrium during the exponential phase results in a system that cannot be reduced to a closed-form solution. Thus, the key assumption that the ANA dissociates immediately upon cleavage is made. The resulting equations predict that fluorescence develops during the exponential phase according to:

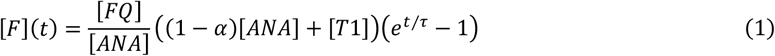

where τ is the time constant:

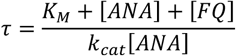

This results in two testable predictions. The exponential phase should extend for the duration:

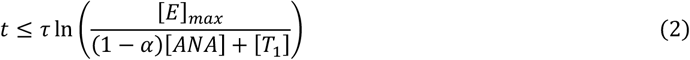

and result in a certain fraction of reporter cleavage:

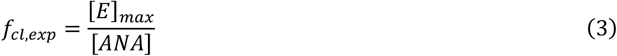

during the exponential phase. During the linear phase, we derive a prediction for the constant rate:

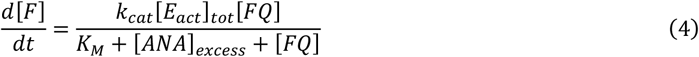

which includes the inhibitory effect of the excess ANA. These predictions are tested in detail in Supplementary Note 10. We observe that the predicted trends as a function of the enzyme, ANA, and reporter concentration are partially captured, although the duration of the exponential phase is routinely underestimated and the linear cleavage rate is overestimated. Figure 5d shows the predicted relationship for the linear phase cleavage rate and Figure 5e shows the observed relationship, with a clear difference in the effect of ANA concentration. If the ANA free energies are idealized to more closely match the assumed behavior, this discrepancy is corrected (Fig. 5f). This indicates that the key limitation is a result of the simplifying assumption made regarding the ANA equilibrium. Therefore, unless an analytical model is developed that encompasses this mechanism, simplified models can only be used for general guidance in this type of system.

The analytical model can still point to potential routes for optimization, but the full numerical model is necessary for optimization experiments. To achieve this, we developed optimization experiment functions (detailed in Supplementary Note 2). For example, these can be used to determine the optimal concentration of ANA for a set of assay conditions. In this case, we applied it to the best-performing hairpin design (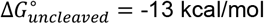 with [E2] ≈ 10 nM and [*FQ*] = 100 nM). Figure 5g demonstrates that a peak SNR is achieved in a 20-minute detection time when the concentration of ANA is set to 100 nM, the same as the reporter concentration. This is roughly in the middle of the ANA range utilized in previous assays (Supplementary Table 1). Detailed plots reveal that increasing the ANA concentration increases the signal amplification, but the background amplification driven by the melted ANA eventually decreases the SNR at high ANA concentrations (Fig. S19). The ratio of ANA to reporter is similarly important and can be explored with the dual variable optimization tool (Fig. 5h). The complete results from this experiment reveal that excessive ANA can drive background amplification and inhibit reporter cleavage, but excessive reporter can inhibit ANA cleavage, slowing the autocatalytic process (Fig. S20).

Other routes of assay optimization have been proposed beyond modification of the free energy and reagent concentrations. This includes modifying the trans-cleavage rate constants by engineering the Cas protein, modifying the crRNA, or adding components that enhance trans-cleavage.^[46–48]^ This can easily be replicated in the demonstrated model by varying the rate constants, including in optimization experiments. Other approaches include variations in the experimental protocol, such as including a pre-activation step.^[19]^ This can also be implemented in our software with little difficulty. Finally, the reaction scheme itself can be modified, such as by adding multiple parallel crRNAs.^[12,49]^ This requires modifying the reaction scheme, which requires additional modifications but is still broadly compatible with the demonstrated model.

## Conclusions

In this work, we developed and demonstrated the first comprehensive kinetic model of autocatalytic CRISPR-Cas-based assays. First, we outlined the reactions composing the model and its implementation. We discussed the principles guiding the design and performance of ANA, a key component in autocatalytic systems, and used our model to expand this discussion from general principles to complete assay performance. We measured the inhibition of trans-cleavage by non-specific ssDNA and implemented a DNA fragmentation mechanism in our model to reflect the likely mechanism reported in literature. We demonstrated that the resulting model can qualitatively replicate the measured inhibition. Next, we used the model to investigate the behavior commonly observed in existing literature. We derived a simplified analytical model for the reaction system and tested its performance against the full numerical model. To the authors’ knowledge, the discrepancy between these models cannot be addressed in a closed-form analytical solution, so a full numerical model is necessary to capture assay performance. Finally, we developed tools capable of running *in silico* optimization experiments over reagent concentrations, ANA free energy values, and reaction constants.

Direct comparison of the model to experimental data comes with substantial challenges. First, existing experimental data is provided in terms of fluorescence rather than reporter cleavage, which makes quantitative comparison difficult. For this reason and in line with previous analysis of single CRISPR-Cas-based assays, we recommend that data normalization and calibration protocols be applied to published data where possible.^[32,33]^ This is achieved easily with the inclusion of a passive reference dye (Supplementary Note 11). Second, the model tracks many species at once. Even when data is converted to concentrations, only a few species can be practically measured in a single reaction system. Finally, while standard reaction parameters (*k*_*cat*_ and *K*_*M*_) can easily reported for comparison across single CRISPR-Cas-based assays, there are not currently simple protocols to directly measure ANA performance beyond melting analysis.

The primary purpose of this work is to provide guidance towards rational design of CRISPR-Cas-based systems. The developed model is provided as open source so that it can be adapted and applied to various reaction schemes. Key improvements to the model should focus on more accurately capturing ANA equilibrium for a wider range of designs, ideally while considering a larger ensemble of secondary structures. With these improvements, future work could progress towards automated testing of ANA designs generated by nucleic acid design software.

## Supporting Information

The authors have cited additional references within the Supporting Information.^[50–58]^ The MATLAB library developed for and used in this manuscript will be made available upon request.

## Acknowledgements

R.B. and E.V. acknowledge partial support from the Jump ARCHES (Applied Research in Community Health through Engineering and Simulation) endowment through the Health Care Engineering Systems Center at UIUC and OSF. This work was partially supported by VinUni-Illinois Smart Health Center and the NIH (R01AI148385 & R01EB032725 A).

We thank the staff at the Holonyak Micro and Nanotechnology Laboratory at UIUC for facilitating the research and the funding from the University of Illinois. Figures 1-5 were created in BioRender.

## Data Availability Statement

The data that supports this study will be made available upon request.

