## Supporting Information for "Kinetic Modeling of Target-Amplification-Free CRISPR-Cas-Based Autocatalysis Reactions"

### Table of Contents

(Page Number)


**Supplementary Figures** (1)

1. Standard measurement of Michaelis-Menten parameters (2)
2. Standard output tabs for the CRISPR-Cas-based autocatalytic model (3)
3. Control experiments (4)
4. Demonstration of change in free energy due to cleavage of a simple hairpin and bulge structure (5)
5. Detailed autocatalytic assay results as a function of hairpin $\Delta G^{\circ}$ for 10 fM target concentration (6)
6. Autocatalytic assay results as a function of hairpin $\Delta G^{\circ}$ for 10 fM target concentration at 60 minutes (7)
7. Autocatalytic assay results as a function of hairpin $\Delta G^{\circ}$ for a range of assay target concentrations (8)
8. Potential cleavage process for a complex ANA (9)
9. Estimation of free energy as a function of enthalpy and entropy for a unimolecular hairpin structure (10)
10. Estimation of free energy as a function of enthalpy and entropy for a bimolecular bulge structure (11)
11. Excess target experiment (12)
12. Full results of inhibition experiment (13)
13. Measurement of Michaelis-Menten parameters from simulated data (14)
14. Simulated Lineweaver-Burk plot (15)
15. Reproduced data demonstrating the different reaction regimes (16)
16. Systematic variation of $\Delta G^{\circ}_{uncleaved}$ and $\Delta G^{\circ}_{cleaved}$ (17)
17. Prolonged exponential behavior under variable $k_{cat}$ and $\Delta\Delta G^{\circ}$ (18)
18. Investigation of the $\Delta\Delta G^{\circ}$ required to maintain a 1 aM limit of detection while reducing detection time (19)
19. Individual reactions in the ANA concentration single variable optimization experiment (20)
20. Complete results from the ANA and reporter concentration dual variable optimization experiment (21)

**Supplementary Notes** (22)

1. Full Reaction Schemes (23)
2. Kinetic Experimental Protocols (28)
3. ANA Rate Constants (31)
4. Model Structure (34)
5. ANA Thermodynamics (44)
6. Gel Experimental Protocol (48)
7. DNA Fragmentation Kinetics (50)
8. Exponential Phase Analytical Simplification (53)
9. Linear Phase Analytical Simplification (56)
10. Evaluation of Analytical Predictions (58)
11. Data Calibration and Normalization Protocol (64)

**Supplementary Tables** (66)

1. CRISPR-Cas-based Autocatalytic Literature (67)
2. Nucleic Acid Sequences and Free Energies (69)

**References** (70)

#### Supplementary Figures

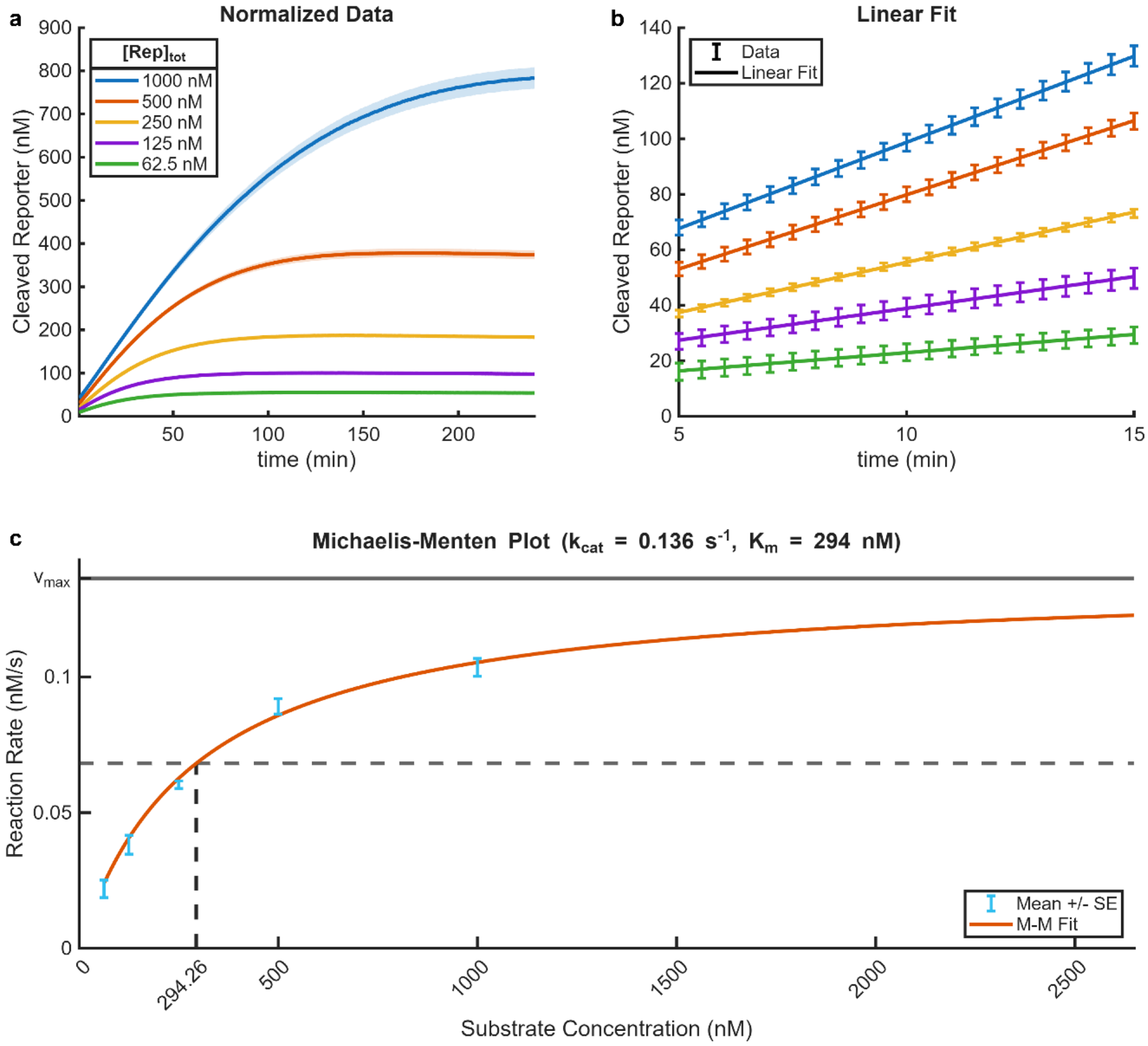


Figure S1. Standard measurement of Michaelis-Menten parameters (activated by unblocked target sequence). a) Cleaved reporter concentration over the entire 4-hour experiment, with varying total substrate (reporter) concentration. The shaded region represents the mean plus or minus the standard error (n=4). b) Selected data over the linear region of the reaction, with a linear fit applied. The error bars represent the mean plus or minus the standard error. c) Michaelis-Menten plot (reaction rate versus substrate concentration), with the plotted Michaelis-Menten fit. The error bars represent the standard error associated with the linear fits from (b).

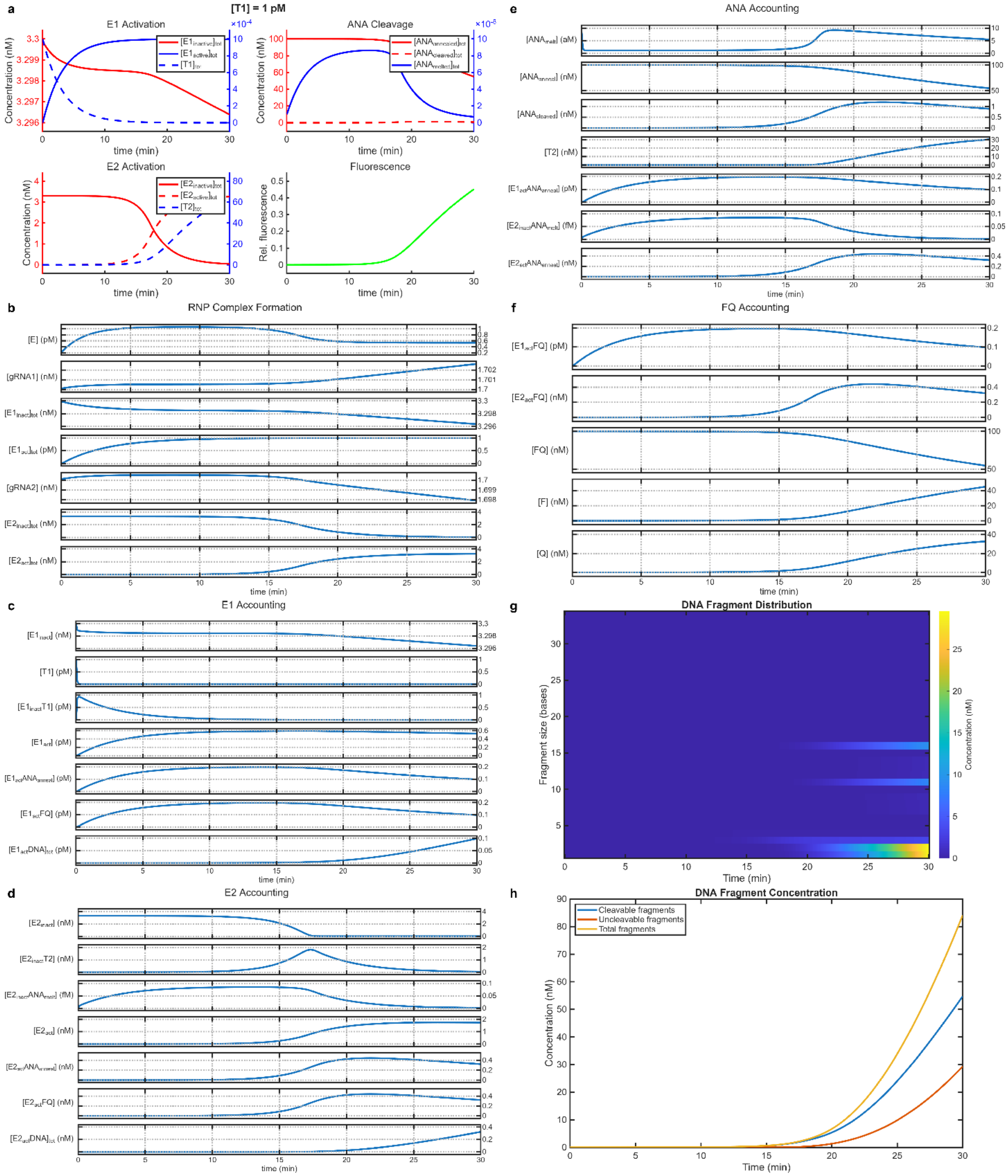

Figure S2. Standard output tabs for the CRISPR-Cas-based autocatalytic model. a) Summary panel of the reaction highlighting E1 activation, ANA cleavage, E2 activation, and fluorescence output. b-f) Plotted concentration of individual species involved in RNP complex formation (b), E1ativation and cleavage (c), E2 activation and cleavage (d), ANA equilibrium (e), and reporter cleavage (f). g) Heatmap showing the concentration of generated ssDNA fragments of various sizes. h) Fragmented ssDNA concentration, showing total fragments as a sum of cleavable or uncleavable fragments.

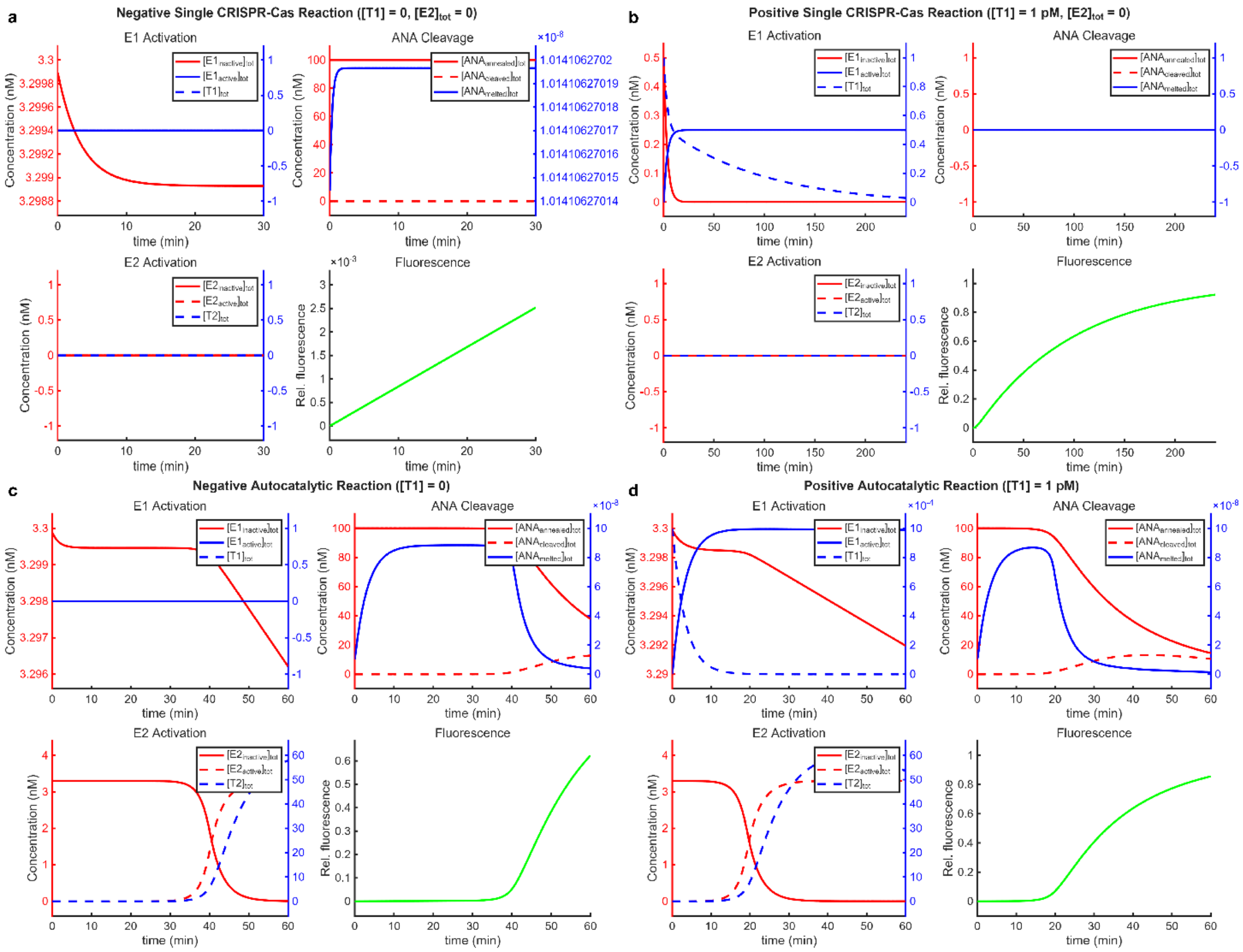


Figure S3. Control experiments. a) Setting the target concentration and the starting concentration of E2 to zero results in behavior typical of a negative single-CRISPR-Cas reaction, with a slow linear increase in fluorescence as a result of non-specific reporter degradation. b) Increasing the target concentration to 1 pM with the concentration of E2 at zero demonstrates a positive single-CRISPR-Cas reaction, with the fluorescence output approximated by Michaelis-Menten kinetics. c) Including all autocatalytic components at zero target concentration results in delayed amplification while E1 does not show activation. This is consistent with the reported negative signal amplification in literature. d) A complete autocatalytic reaction with a target concentration of 1 pM results in rapid signal amplification after an initial delay.

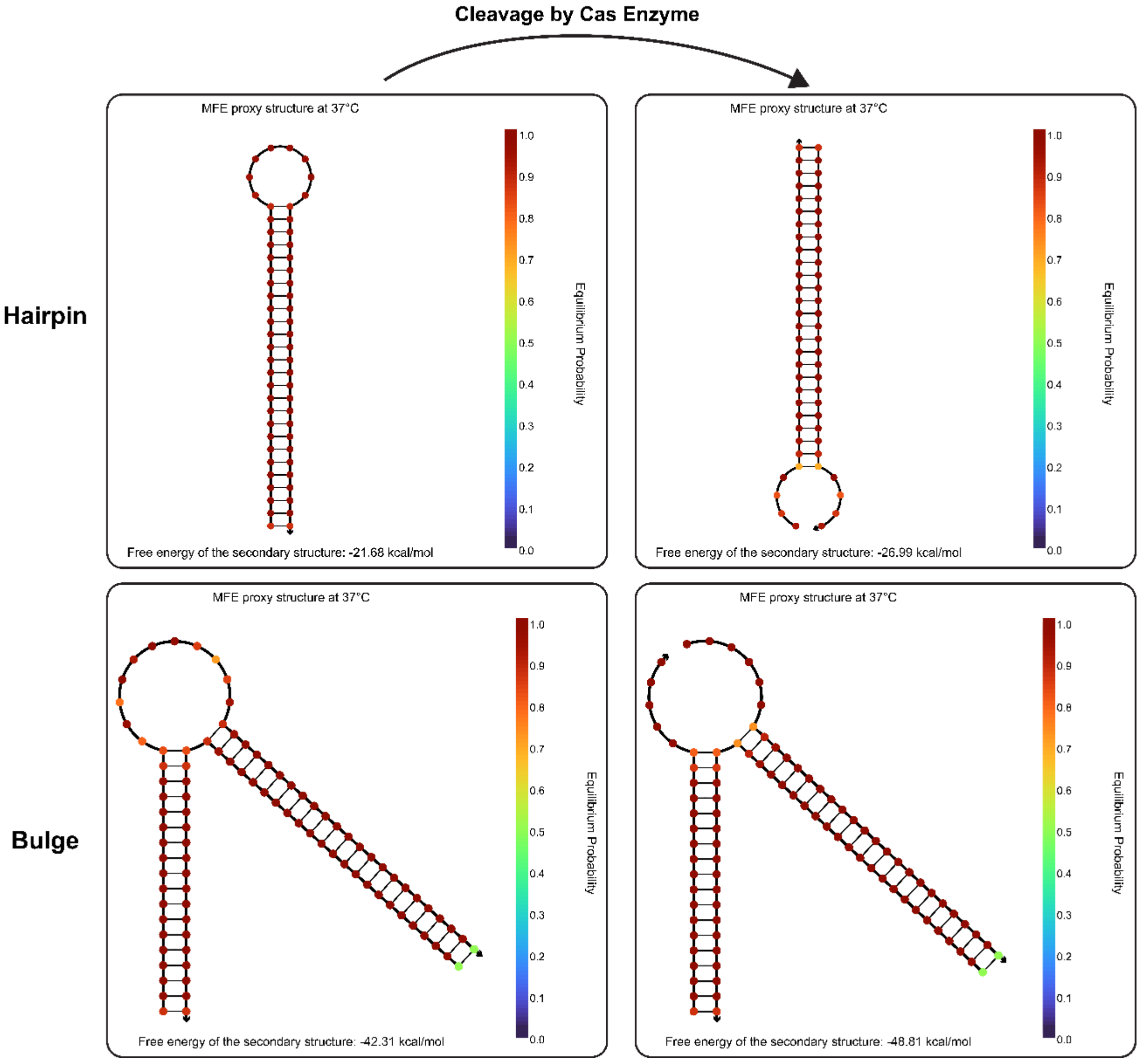


Figure S4. Demonstration of change in free energy due to cleavage of a simple hairpin (Hairpin10) and bulge (Bulge11) structure. The free energy of the MFE secondary structure can be seen to decrease as a result of cleavage. Structures were modeled using the NUPACK web server with strand concentrations of 100 nM, temperature of 37°C, [$Na^+$] of 50 mM, and [$Mg^{++}$] of 10 mM.[1,2] Exact sequences are provided in Supplementary Table 2.

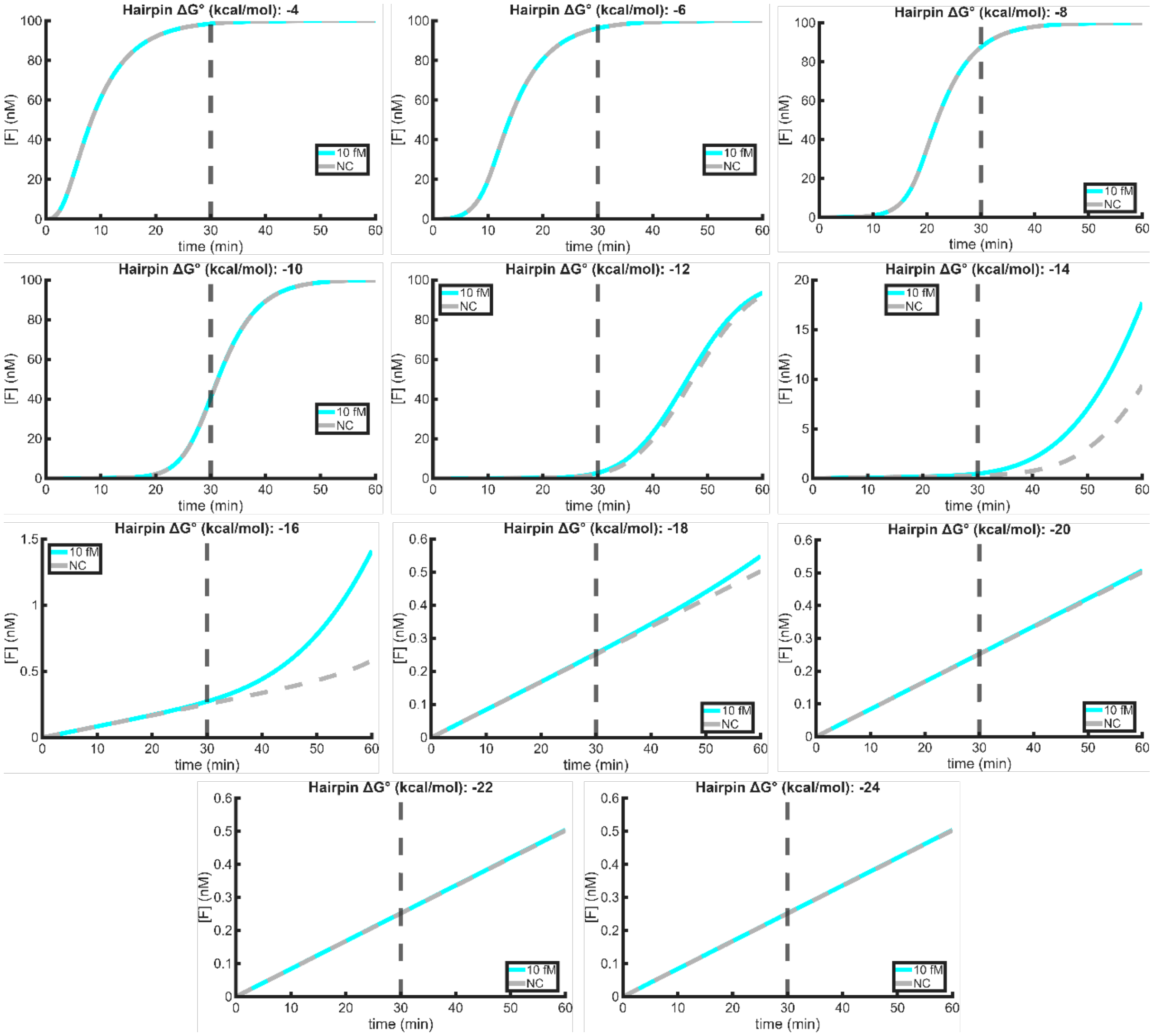


Figure S5. Detailed autocatalytic assay results as a function of hairpin $\Delta G^{\circ}$ for 10 fM target concentration. The difference between the positive and negative samples can be seen to increase as the hairpin free energy decreases. Once the free energy is below the optimal range, the observed signal approaches linear behavior, similar to a single-CRISPR-Cas assay. For these experiments, reporter concentration is 100 nM and all other rates and concentrations are set at their default values.

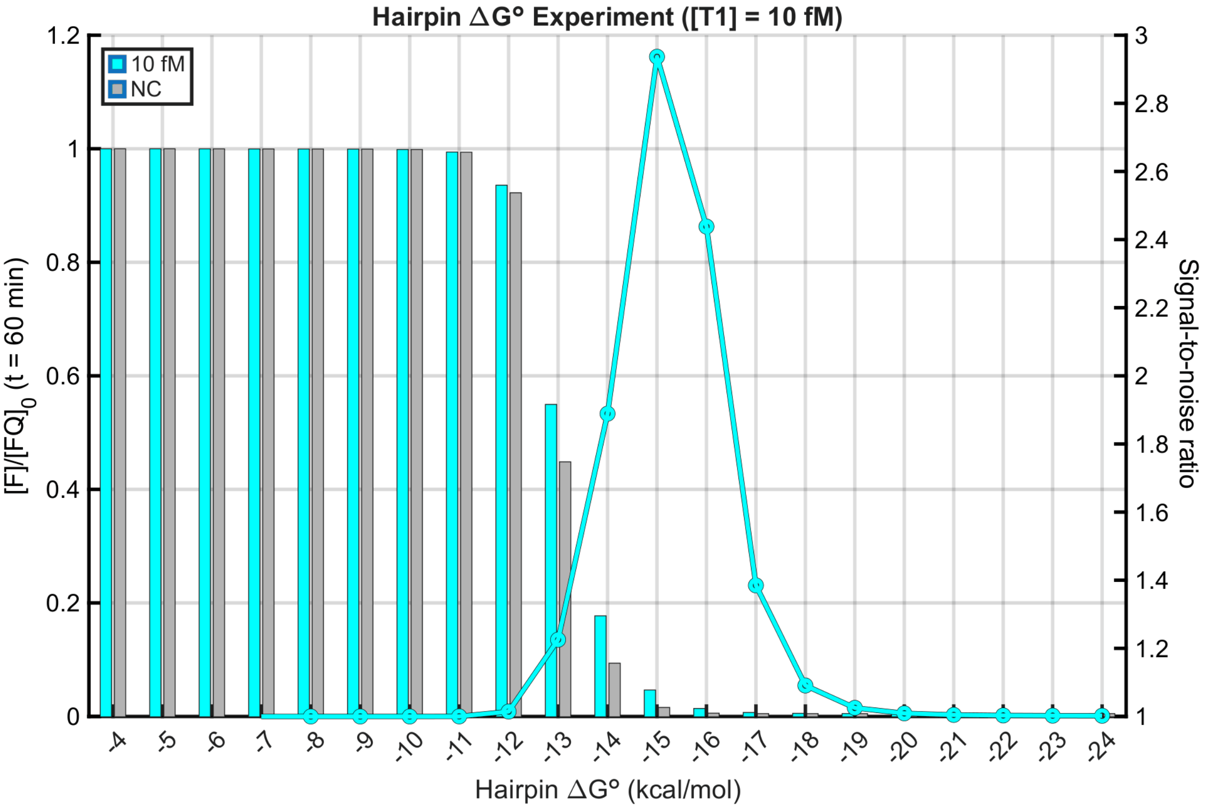


Figure S6. Autocatalytic assay results as a function of hairpin $\Delta G^{\circ}$ for 10 fM target concentration at 60 minutes. The signal-to-noise ratio reaches a maximum of nearly 3, compared to ~1.75 at 30 min.

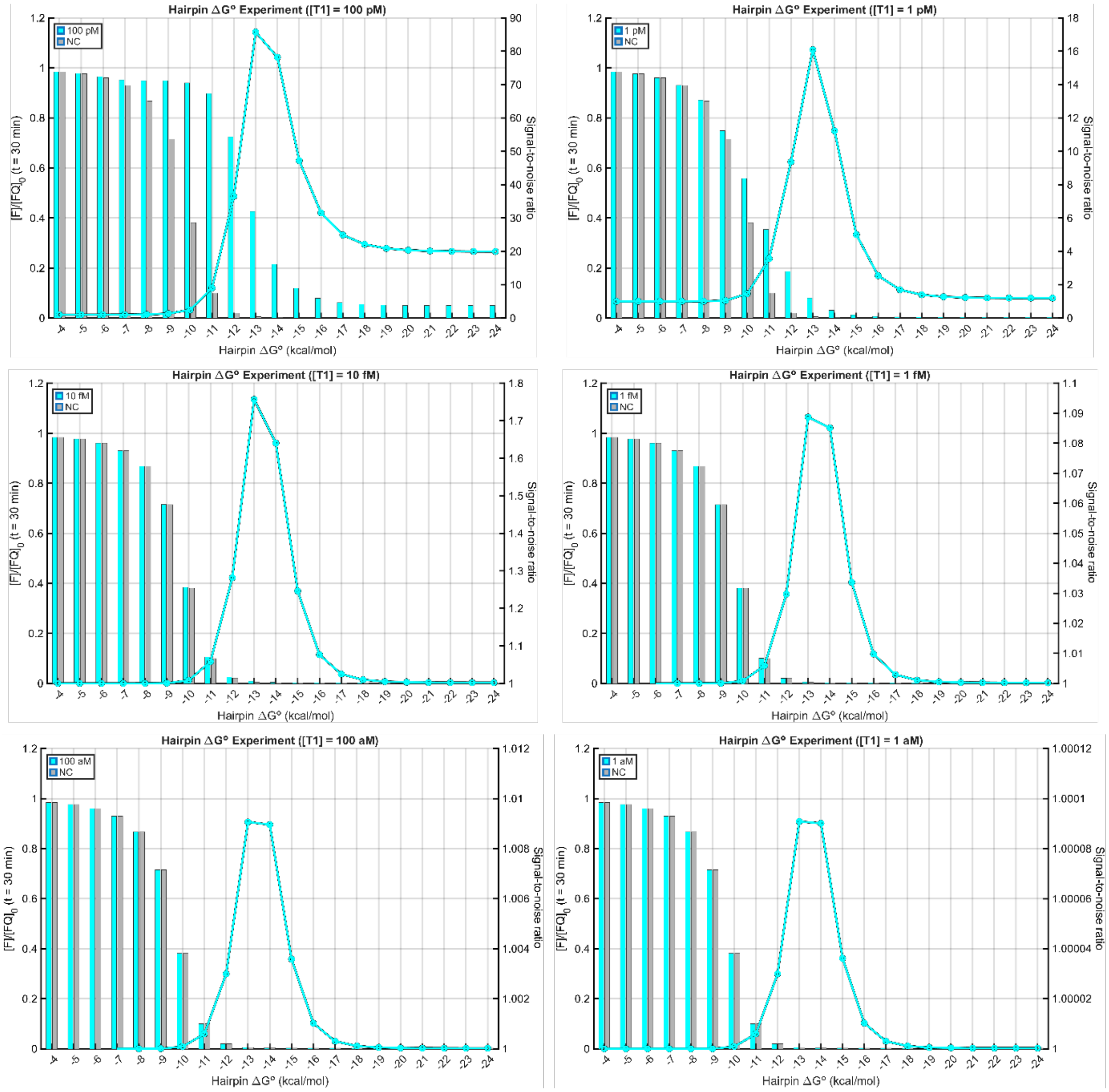


Figure S7. Autocatalytic assay results as a function of hairpin $\Delta G^\circ$ for a range of assay target concentrations. The peak SNR is observed to increase as the target concentration increases. For these experiments, the reporter concentration is 100 nM and all other rates and concentrations are set at their default values.

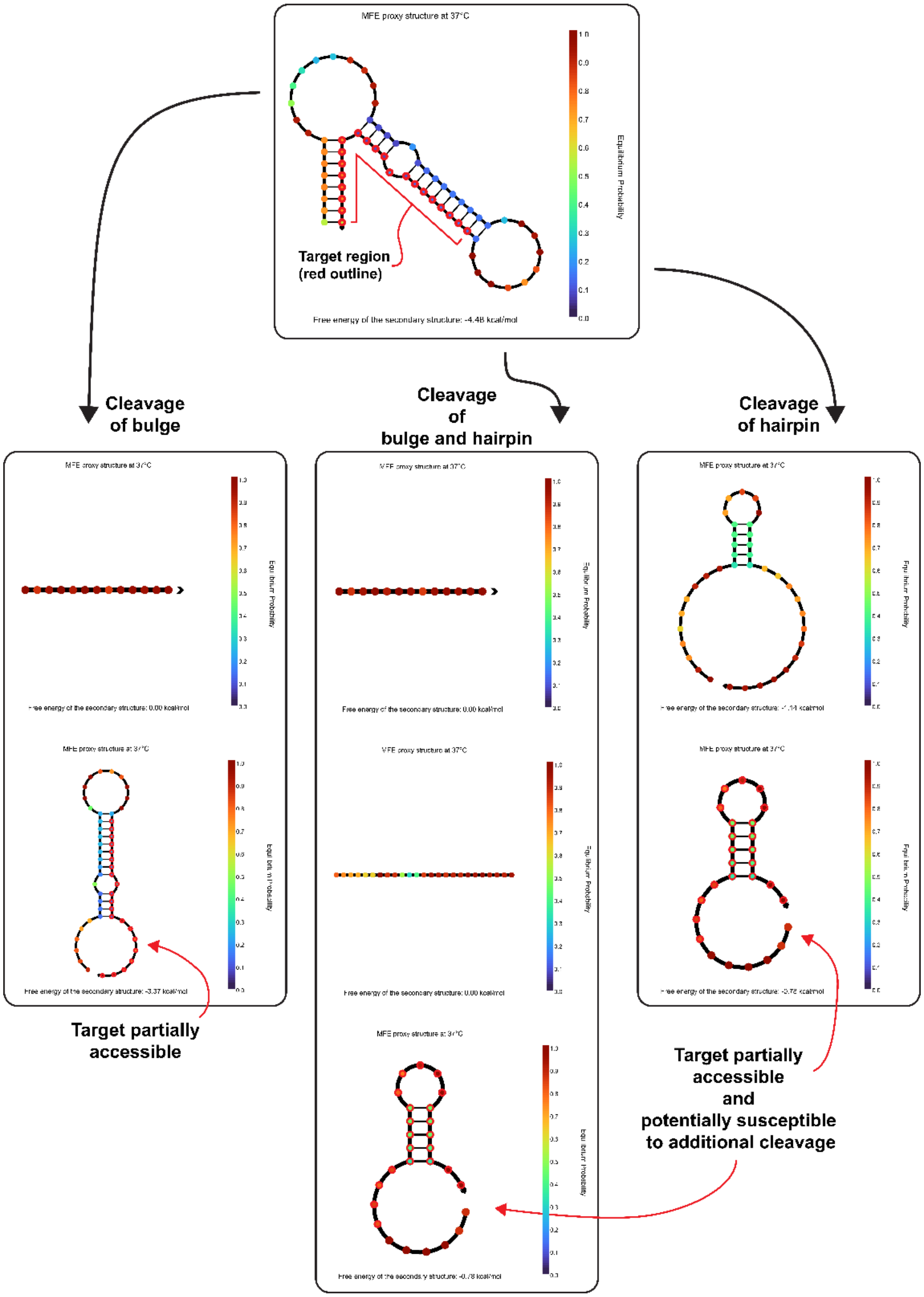


Figure S8. Potential cleavage process for a complex ANA. The ANA previously demonstrated by our group contains a bulge, hairpin, and loop in the minimum free energy (MFE) secondary structure. The target region for the RNP complex is outlined in red. After cleavage at one or both likely cleavage sites, each resulting set of strands has its own predicted MFE secondary structures, which may partially or fully expose the target. Further breakdown and interaction of these cleavage products may occur, complicating modeling of the ANA activation. Structures were modeled using the NUPACK web server with strand concentrations of 100 nM, temperature of 37°C, [$Na^+$] of 50 mM, and [$Mg^{++}$] of 10 mM.[1,2] Exact sequences are provided in Supplementary Table 2.

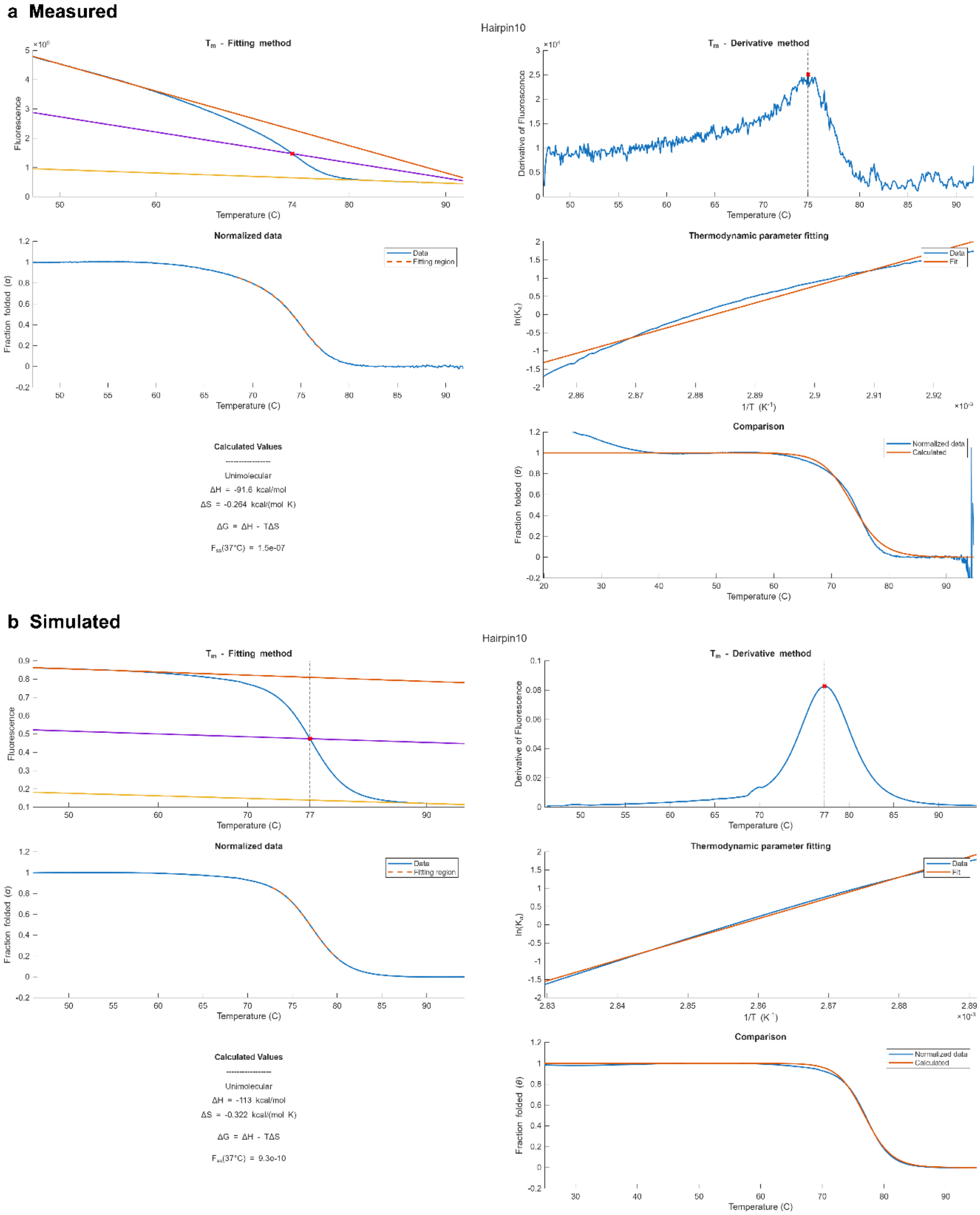


Figure S9. Estimation of free energy as a function of enthalpy and entropy for a unimolecular hairpin structure, following the method outlined by Mergny and Lacroix.[3] To estimate the enthalpy and entropy, the melting curve is normalized to the range of [0,1] using linear fits for the upper and lower linear regions. The melting temperature occurs where the midline of these fits intersects the melting curve. After normalization, the roughly linear region near the melting point is transformed to the form $\ln(K_a)$ vs. $1/T$ and fit to the linear form of the van't Hoff equation to obtain the enthalpy and entropy values. This process is shown for (a) measured and (b) simulated melting curves. Melting curves were measured according to Supplementary Note 2 and simulated using the NUPACK web server with strand concentrations of 100 nM, temperature of 37°C, [$Na^+$] of 50 mM, and [$Mg^{++}$] of 10 mM.[1,2] Exact sequences are provided in Supplementary Table 2.

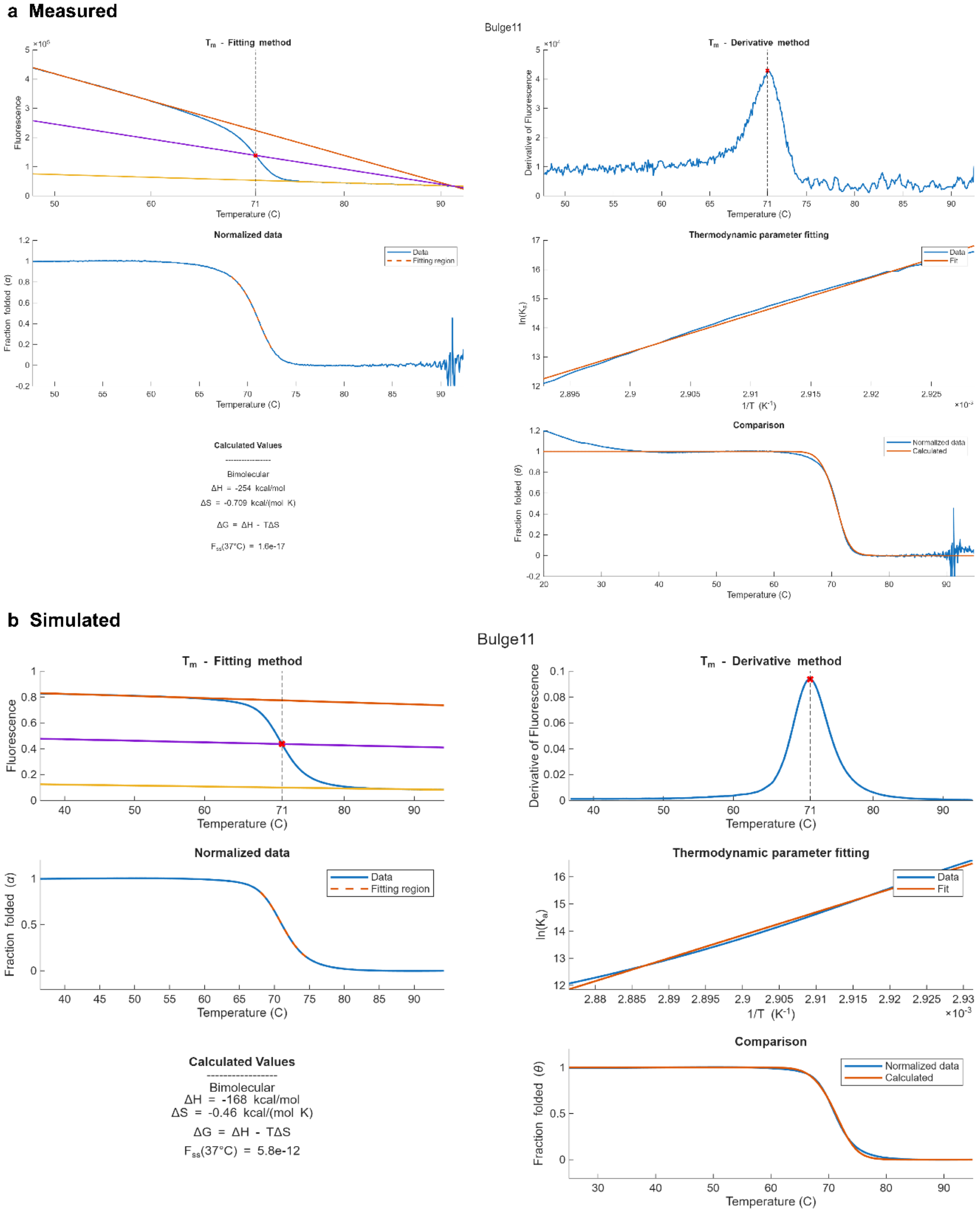


Figure S10. Estimation of free energy as a function of enthalpy and entropy for a unimolecular hairpin structure, following the method outlined by Mergny and Lacroix.[3] To estimate the enthalpy and entropy, the melting curve is normalized to the range of [0,1] using linear fits for the upper and lower linear regions. The melting temperature occurs where the midline of these fits intersects the melting curve. After normalization, the roughly linear region near the melting point is transformed to the form $\ln(K_a)$ vs. $1/T$ and fit to the linear form of the van't Hoff equation to obtain the enthalpy and entropy values. This process is shown for (a) measured and (b) simulated melting curves. Melting curves were measured according to Supplementary Note 2 and simulated using the NUPACK web server with strand concentrations of 100 nM, temperature of 37°C, [$Na^+$] of 50 mM, and [$Mg^{++}$] of 10 mM.[1,2] Exact sequences are provided in Supplementary Table 2.

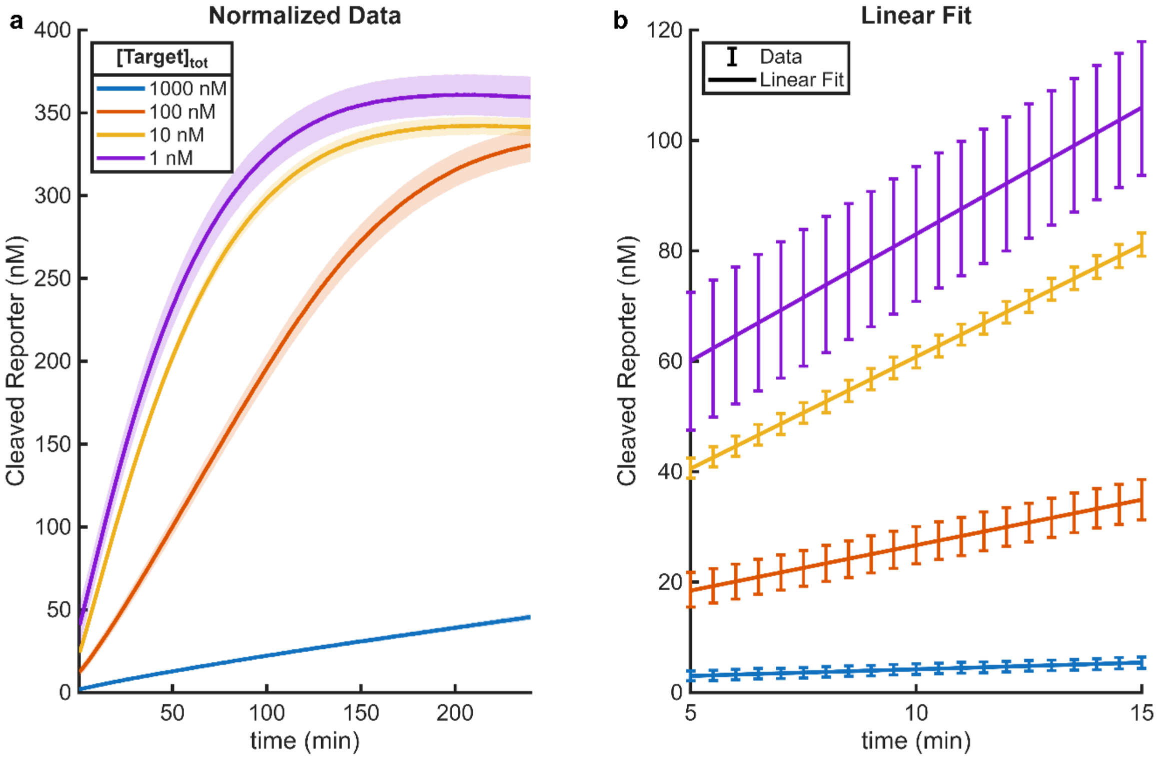


Figure S11. Excess target experiment. a) Cleaved reporter concentration data over the entire 4-hour experiment, with varying unblocked target concentration. The concentration of activated Cas enzyme in each case was approximately 1 nM. The shaded region represents the mean plus or minus the standard error (n=4). b) Selected data over the linear region of the reaction, with a linear fit applied. The error bars represent the mean plus or minus the standard error.

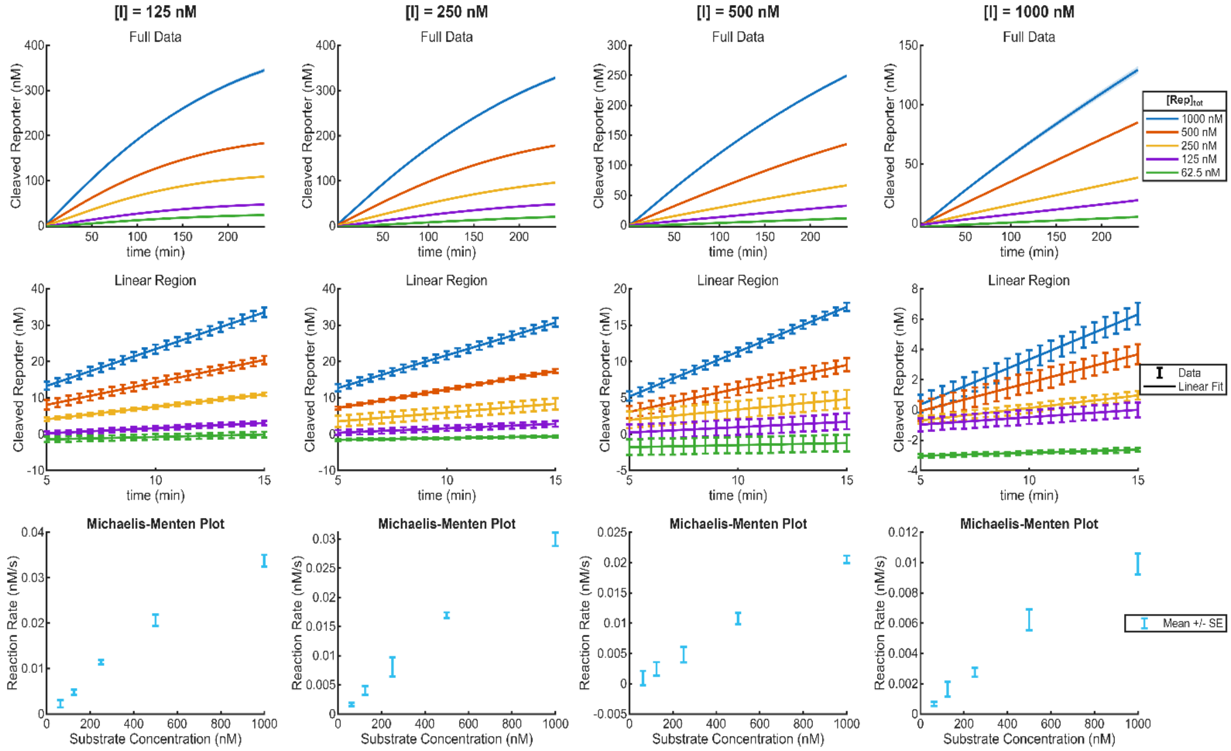


Figure S12. Full results of inhibition experiment. Each inhibitor concentration results in a complete Michaelis-Menten experiment (top row). Shaded regions indicate the value plus or minus the standard error (n=4). The region of roughly linear increase is isolated from the normalized and calibrated data and a linear fit is applied, with error bars representing the data plus or minus the standard error (middle row). Michaelis-Menten plots can then be generated for each inhibitor concentration, with the error bars indicating slope of the fit plus or minus the standard error of the linear fit (bottom row). The Michaelis-Menten plots for high inhibitor concentrations appear nearly linear, indicating that a much higher substrate concentration is required to reach the maximum reaction rate in the presence of high inhibitor concentration.

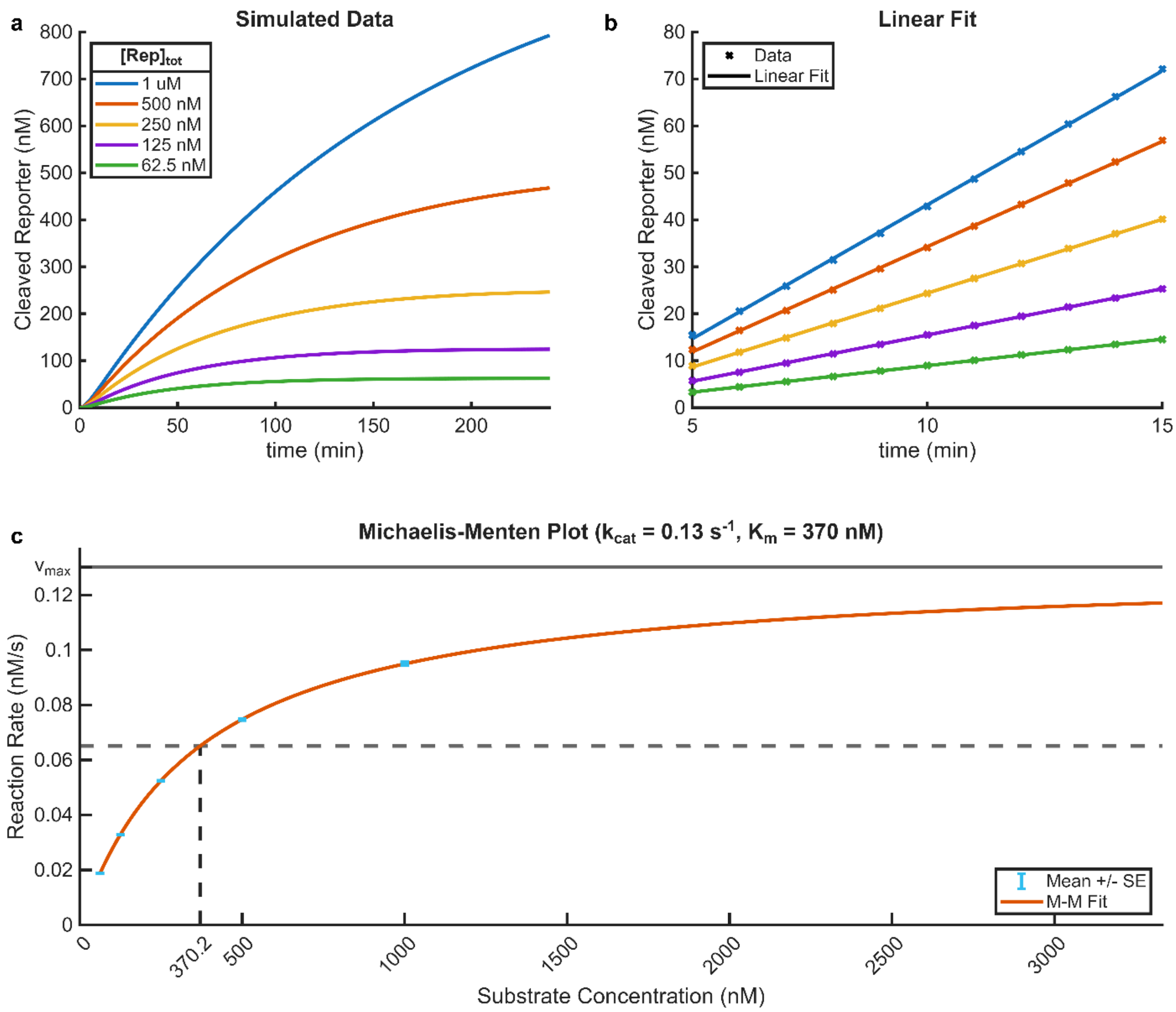


Figure S13. Measurement of Michaelis-Menten parameters from simulated data. a) Cleaved reporter concentration over the entire 4-hour experiment, with varying total substrate (reporter) concentration. b) Selected data over the linear region of the reaction, with a linear fit applied. No error bars are provided for the simulated data. c) Michaelis-Menten plot (reaction rate versus substrate concentration), with the plotted Michaelis-Menten fit. The error bars represent the standard error associated with the linear fits from (b), which is minimal for the simulated data.

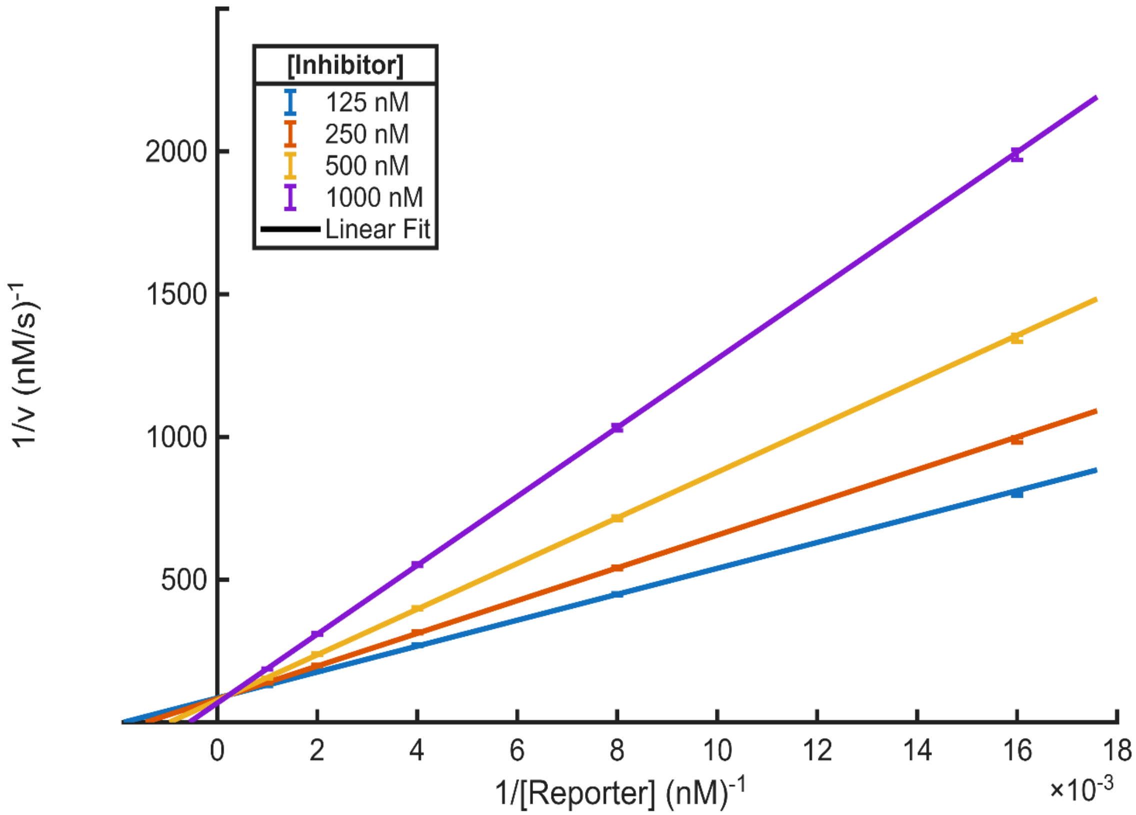


Figure S14. Simulated Lineweaver-Burk plot, repeated with $k_{cat}$ for trans-cleavage set as 0.01 $s^{-1}$, at the low end of the reported range for Cas12.[4]

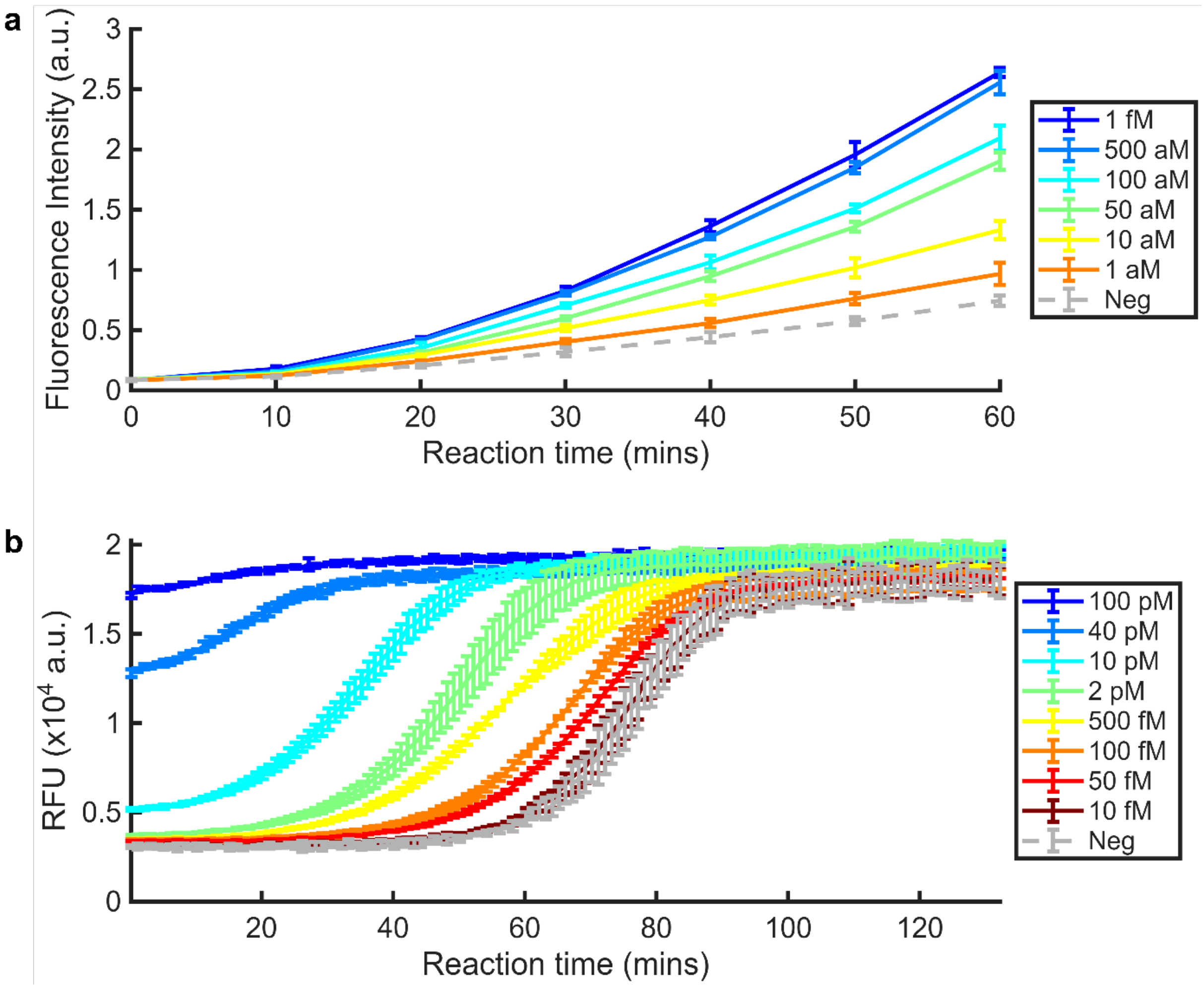


Figure S15. Reproduced data demonstrating the different reaction regimes. a) Prolonged exponential reaction demonstrated by Deng et al. with AutoCAR-1, reproduced from raw data.[5] b) Delayed sigmoidal amplification curve demonstrated by Sun et al. with split activators, reproduced from raw data.[6]

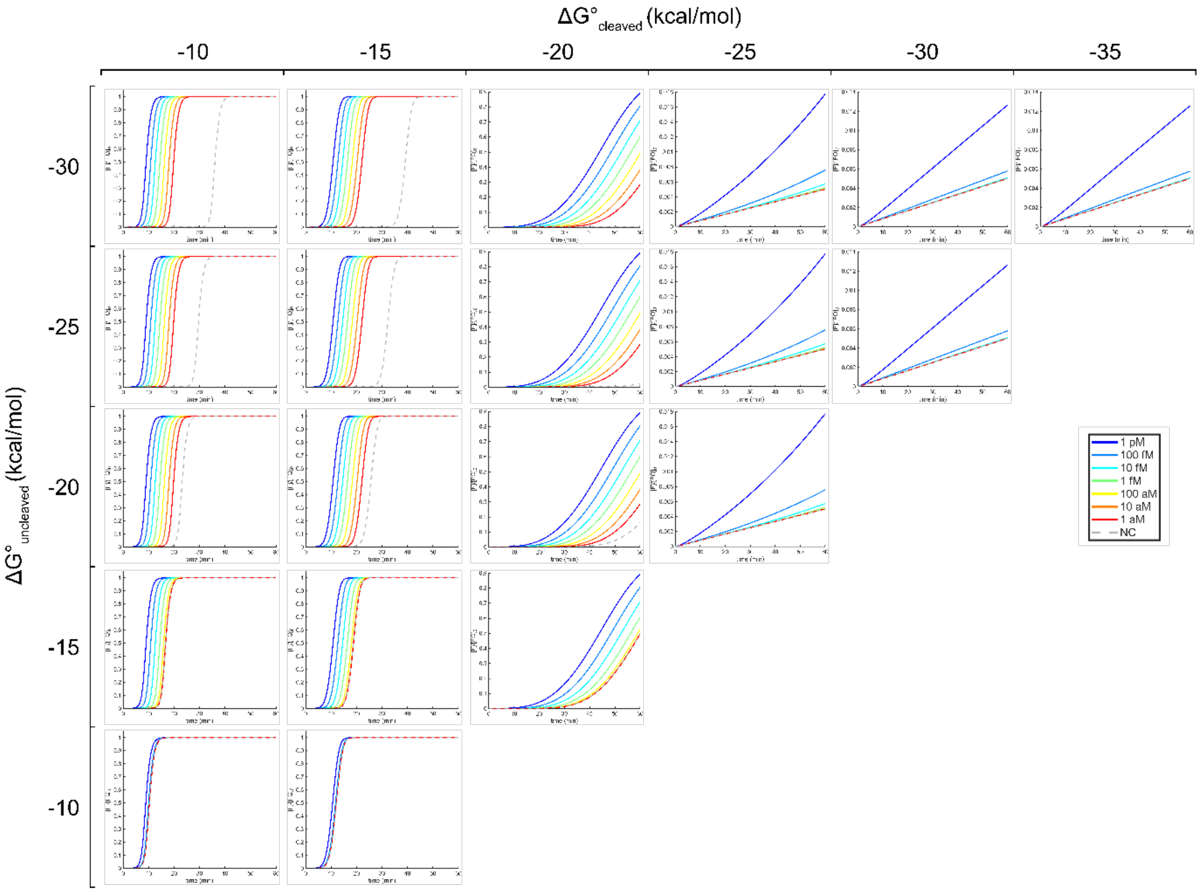


Figure S16. Systematic variation of $\Delta G^{\circ}_{uncleaved}$ and $\Delta G^{\circ}_{cleaved}$ ($[E2]$ ≈ 10 nM, $[ANA]$ ≈ 50 nM, $[FQ]$ = 100 nM, $k_{cat}$ = 1 $s^{-1}$). This demonstrates that $\Delta G^{\circ}_{uncleaved}$ primarily controls the background amplification and $\Delta G^{\circ}_{cleaved}$ primarily controls the signal amplification. However, high background dominates the behavior of the reaction when $\Delta G^{\circ}_{uncleaved}$ is high.

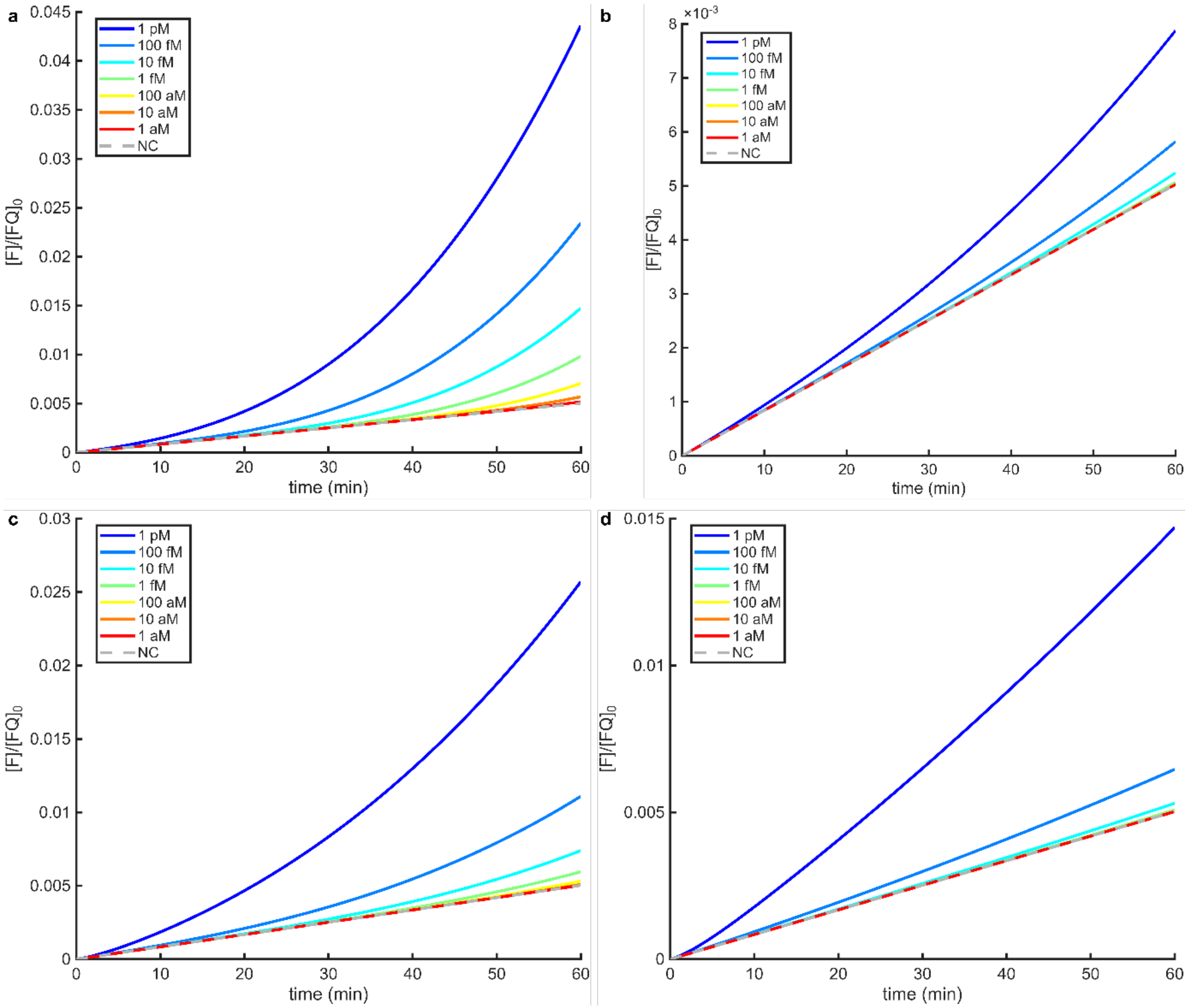


Figure S17. Prolonged exponential behavior under variable $k_{cat}$ and $\Delta\Delta G^{\circ}$ ($[E2] \approx$ 10 nM, $[ANA] \approx$ 50 nM, $[FQ] =$ 100 nM, $\Delta G^{\circ}_{uncleaved} =$ -26 kcal/mol). 1) $\Delta\Delta G^{\circ}$ = 4 kcal/mol and $k_{cat}$ = 0.5 s$^{-1}$. b) $\Delta\Delta G^{\circ}$ = 4 kcal/mol and $k_{cat}$ = 0.1 s$^{-1}$. c) $\Delta\Delta G^{\circ}$ = 2 kcal/mol and $k_{cat}$ = 1 s$^{-1}$. d) $\Delta\Delta G^{\circ}$ = 0 kcal/mol and $k_{cat}$ = 1 s$^{-1}$. The apparent sensitivity is observed to decrease as $\Delta\Delta G^{\circ}$ and $k_{cat}$ decrease.

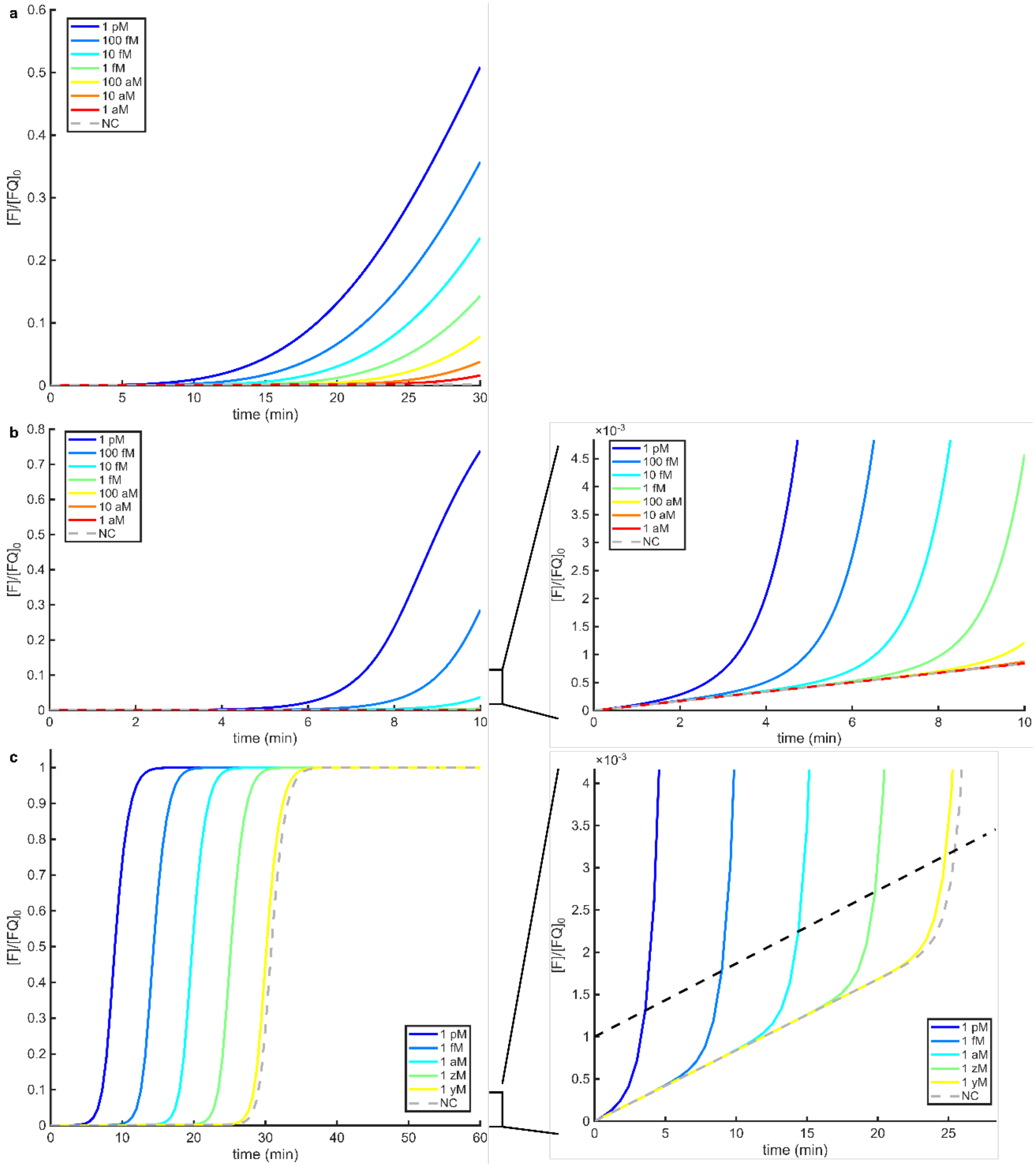


Figure S18. Investigation of the $\Delta\Delta G^{\circ}$ required to maintain a 1 aM limit of detection while reducing the detection time ($[E2] \approx$ 10 nM, $[ANA] \approx$ 50 nM, $[FQ] =$ 100 nM, $k_{cat} =$ 1 s$^{-1}$, $\Delta G^{\circ}_{uncleaved} =$ -26 kcal/mol). A 60-minute reaction requires $\Delta\Delta G^{\circ} \approx 4$ kcal/mol to distinguish between 1 aM and background. a) To achieve similar performance in 30 minutes, $\Delta\Delta G^{\circ}$ must be increase to approximately 7 kcal/mol. b) Increasing $\Delta\Delta G^{\circ}$ to an arbitrarily large number does not achieve 1 aM detection within 10 minutes in this scheme. Closer examination reveals that the limit of detection is on the order of 1 fM at 10 minutes, assuming >0.1% of reporter must be cleaved for detection. c) Repeating this observation reveals the theoretically optimal time for detection with the selected $\Delta G^{\circ}_{hairpin}$. The dashed line indicates where the cleaved reporter concentration is greater than the background by 0.1% of the reporter concentration (0.1 nM).

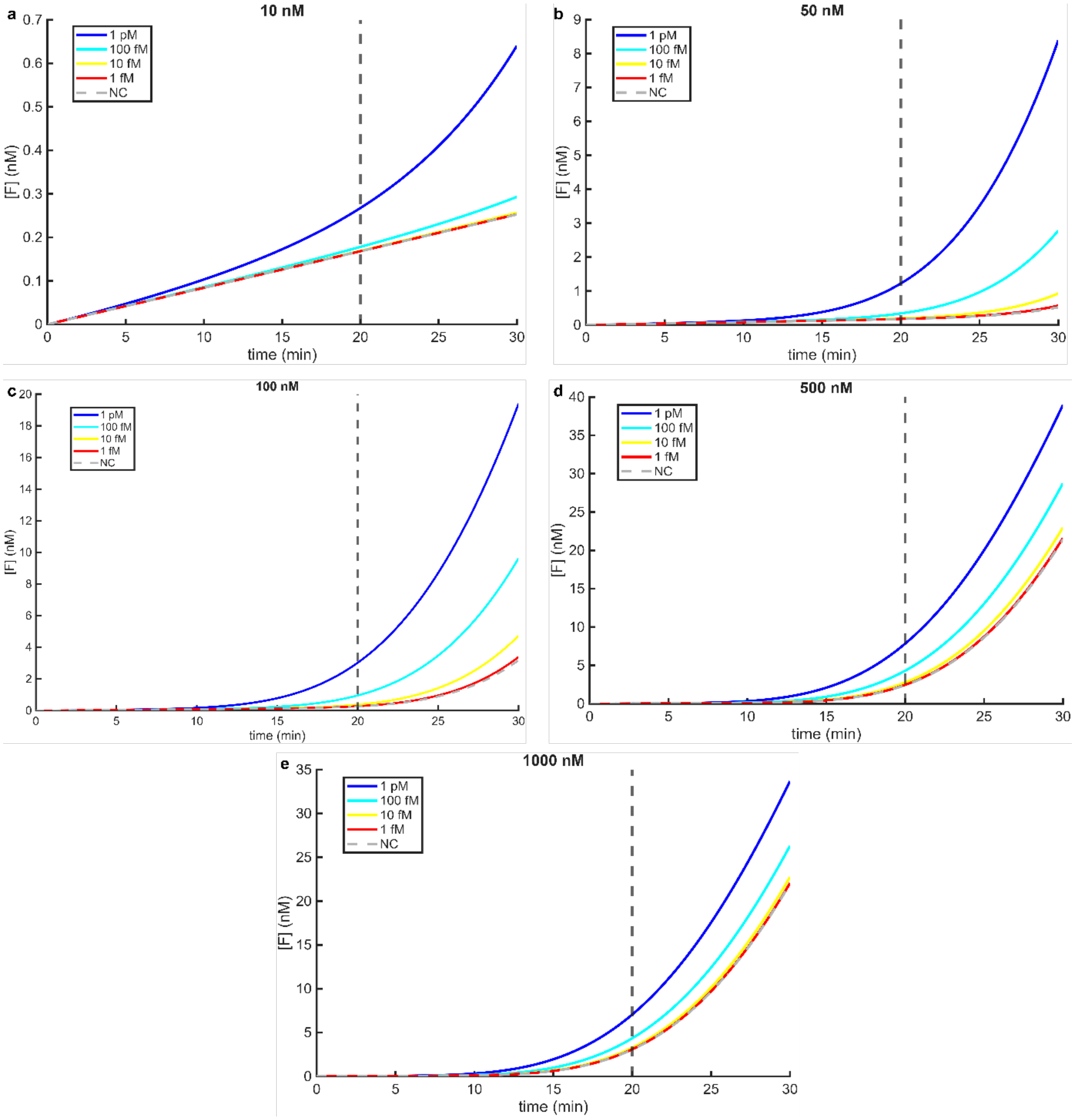


Figure S19. Individual reactions in the ANA concentration single variable optimization experiment ($\Delta G^{\circ}_{uncleaved}$ = -13 kcal/mol, $\Delta G^{\circ}_{cleaved}$ = -17 kcal/mol, $[E2] \approx$ 10 nM, $[FQ]$ = 100 nM).

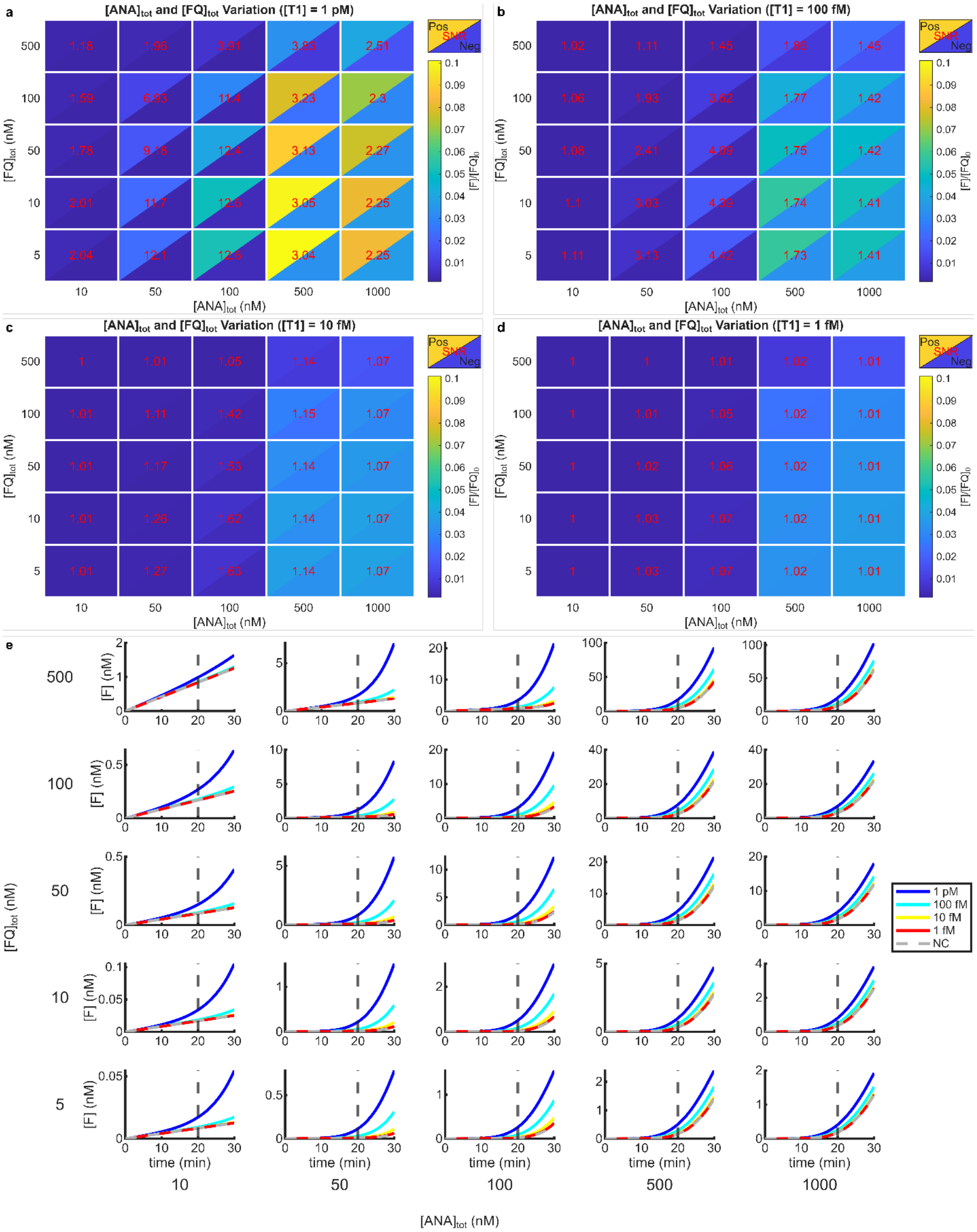


Figure S20. Complete results from the ANA and reporter concentration dual variable optimization experiment ($\Delta G^{\circ}_{uncleaved}$ = -13 kcal/mol, $\Delta G^{\circ}_{cleaved}$ = -17 kcal/mol, $[E2] \approx$ 10 nM).

#### Supplementary Notes

#### Supplementary Note 1: Full Reaction Schemes

##### Hairpin (Single-component ANA)

This reaction is based on the ANA design HairpinOpt. The full sequence is provided in Supplementary Table 2.

Main reactions

| | | |
|---|---|---|
| RNP1 Formation: | $E + gRNA1 \rightleftharpoons E1_{inact}$ | (1) |
| RNP2 Formation: | $E + gRNA2 \rightleftharpoons E2_{inact}$ | (2) |
| RNP1 Activation (by T1)*: | $E1_{inact} + T1 \rightleftharpoons E1_{inact}T1 \rightarrow E1_{act} + DNA_9 + DNA_{15}$ | (3) |
| RNP1 Inactivation*: | $E1_{act} \rightarrow E1_{inact} + DNA_{26} + DNA_{32}$ | (4) |
| ANA Equilibrium (Uncleaved): | $ANA_{annealed} \rightleftharpoons ANA_{melted}$ | (5) |
| ANA Equilibrium (Cleaved): | $ANA_{cleaved} \rightleftharpoons T2 + T2C$ | (6) |
| RNP2 Activation (by T2)*: | $E2_{inact} + T2 \rightleftharpoons E2_{inact}T2 \rightarrow E2_{act} + DNA_4$ | (7) |
| RNP2 Inactivation*: | $E2_{act} \rightarrow E2_{inact} + DNA_{22}$ | (8) |
| RNP2 Activation (by $ANA_{melted}$)*: | $E2_{inact} + ANA_{melted} \rightleftharpoons E2_{inact}ANA_{melted} \rightarrow E2_{act} + DNA_{30}$ | (9) |
| $ANA_{annealed}$ Trans-cleavage (by E1): | $E1_{act} + ANA_{annealed} \rightleftharpoons E1_{act}ANA_{annealed} \rightarrow E1_{act} + ANA_{cleaved}$ | (10) |
| $ANA_{annealed}$ Trans-cleavage (by E2): | $E2_{act} + ANA_{annealed} \rightleftharpoons E2_{act}ANA_{annealed} \rightarrow E2_{act} + ANA_{cleaved}$ | (11) |
| Reporter Trans-cleavage (by E1): | $E1_{act} + FQ \rightleftharpoons E1_{act}FQ \rightarrow E1_{act} + F + Q$ | (12) |
| Reporter Trans-cleavage (by E2): | $E2_{act} + FQ \rightleftharpoons E2_{act}FQ \rightarrow E2_{act} + F + Q$ | (13) |
| Reporter degradation: | $FQ \rightarrow F + Q$ | (14) |

Component degradation

| | | |
|---|---|---|
| T2C Trans-cleavage (by E1)*: | $E1_{act} + T2C \rightleftharpoons E1_{act}T2C \rightarrow E1_{act} + DNA_3 + DNA_{23}$ | (15) |
| T2C Trans-cleavage (by E2)*: | $E2_{act} + T2C \rightleftharpoons E2_{act}T2C \rightarrow E2_{act} + DNA_3 + DNA_{23}$ | (16) |
| $ANA_{melted}$ Trans-cleavage (by E1)*: | $E1_{act} + ANA_{melted} \rightleftharpoons E1_{act}ANA_{melted} \rightarrow E1_{act} + DNA_3 + DNA_{49}$ | (17) |
| $ANA_{melted}$ Trans-cleavage (by E2)*: | $E2_{act} + ANA_{melted} \rightleftharpoons E2_{act}ANA_{melted} \rightarrow E2_{act} + DNA_3 + DNA_{49}$ | (18) |
| T1 Trans-cleavage (by E1)*: | $E1_{act} + T1 \rightleftharpoons E1_{act}T1 \rightarrow E1_{act} + DNA_3 + DNA_{38}$ | (19) |
| T1 Trans-cleavage (by E2)*: | $E2_{act} + T1 \rightleftharpoons E2_{act}T1 \rightarrow E2_{act} + DNA_3 + DNA_{38}$ | (20) |
| T2 Trans-cleavage (by E1)*: | $E1_{act} + T2 \rightleftharpoons E1_{act}T2 \rightarrow E1_{act} + DNA_3 + DNA_{23}$ | (21) |
| T2 Trans-cleavage (by E2)*: | $E2_{act} + T2 \rightleftharpoons E2_{act}T2 \rightarrow E2_{act} + DNA_3 + DNA_{23}$ | (22) |
| Q Trans-cleavage (by E1)*: | $E1_{act} + Q \rightleftharpoons E1_{act}Q \rightarrow E1_{act} + DNA_2 + DNA_3$ | (23) |
| Q Trans-cleavage (by E2)*: | $E2_{act} + Q \rightleftharpoons E2_{act}Q \rightarrow E2_{act} + DNA_2 + DNA_3$ | (24) |

*Releases ssDNA fragments

##### Rates

These are the default rates set in the model.

1) $k_f = 2.9 \times 10^6\ M^{-1}\mathrm{s}^{-1}, k_r = 1.6 \times 10^{-6}\ \mathrm{s}^{-1}$.[7]
2) $k_f = 2.9 \times 10^6\ M^{-1}\mathrm{s}^{-1}, k_r = 1.6 \times 10^{-6}\ \mathrm{s}^{-1}$.[7]
3) $k_f = 1.1 \times 10^8\ M^{-1}\mathrm{s}^{-1}, k_r = 5.9 \times 10^{-6}\ \mathrm{s}^{-1}, k_{cat} = 5.1 \times 10^{-3}\ \mathrm{s}^{-1}$.[8]
4) $k_{inact} = 6 \times 10^{-6}\ s^{-1}$.[8]
5) Solved by calculateEquilibriumRates function.
6) Solved by calculateEquilibriumRates function.
7) $k_f = 1.1 \times 10^8\ M^{-1}\mathrm{s}^{-1}, k_r = 5.9 \times 10^{-6}\ \mathrm{s}^{-1}, k_{cat} = 5.1 \times 10^{-3}\ \mathrm{s}^{-1}$.[8]
8) $k_{inact} = 6 \times 10^{-6}\ s^{-1}$.[8]
9) $k_f = 1.1 \times 10^8\ M^{-1}\mathrm{s}^{-1}, k_r = 5.9 \times 10^{-6}\ \mathrm{s}^{-1}, k_{cat} = 5.1 \times 10^{-3}\ \mathrm{s}^{-1}$.[8]
10) $K_M = 294 \times 10^{-9}\ M, k_{cat} = 0.136\ s^{-1}, k_f = 10^8\ M^{-1}s^{-1}$ (Measured, Supplementary Figure 1)
11) $K_M = 294 \times 10^{-9}\ M, k_{cat} = 0.136\ s^{-1}, k_f = 10^8\ M^{-1}s^{-1}$ (Measured, Supplementary Figure 1)
12) $K_M = 294 \times 10^{-9}\ M, k_{cat} = 0.136\ s^{-1}, k_f = 10^8\ M^{-1}s^{-1}$ (Measured, Supplementary Figure 1)
13) $K_M = 294 \times 10^{-9}\ M, k_{cat} = 0.136\ s^{-1}, k_f = 10^8\ M^{-1}s^{-1}$ (Measured, Supplementary Figure 1)
14) $k_{rep} = 1.4 \times 10^{-6}\ s^{-1}$.[9]

15-24) $K_M = 294 \times 10^{-9}\ M, k_{cat} = 0.136\ s^{-1}, k_f = 10^8\ M^{-1}s^{-1}$ (Measured, Supplementary Figure 1)

**Species Descriptions**
Key species are named. Species such as $E1_{inact}T1$ indicate enzyme-substrate complexes. Species in the format $DNA_n$ indicate a single-stranded DNA fragment of length n.
$E$ – Cas enzyme (without gRNA)
$gRNA1$ – gRNA for the assay target-specific RNP complex
$E1_{inact}$ – Inactive assay target-specific RNP complex, before target recognition
$gRNA2$ – gRNA for the ANA target-specific RNP complex
$E2_{inact}$ – Inactive ANA target-specific RNP complex, before target recognition
$T1$ – Assay target
$E1_{act}$ – Active assay target-specific RNP complex, after target recognition
$ANA_{annealed}$ – ANA in the annealed form (inaccessible to Cas enzyme)
$ANA_{melted}$ – ANA in the melted form (accessible to Cas enzyme)
$ANA_{cleaved}$ – ANA in the cleaved form (dissociates into T2 and T2C)
$T2$ – ANA target
$T2C$ – Complement to ANA target
$E2_{act}$ - Active ANA target-specific RNP complex, after target recognition
$FQ$ – Reporter molecule
$F$ – Fluorescent component of cleaved reporter molecule
$Q$ – Quencher component of cleaved reporter molecule

**DNA Fragments**
The intention of the DNA terms is to approximate the creation of single-stranded DNA fragments that may act as reaction inhibitors. The process for approximating the lengths was as follows.
For Cas12a cis-cleavage, the target strand is reported to be cleaved approximately 21-23 nucleotides from the 5' end of the spacer (moving towards the 5' end of the target strand).[10,11] Non-target strand cleavage is reported to occur 15 nucleotides from the 5' end of the spacer. The PAM-distal fragments are released while the PAM-proximal fragments are retained in the active enzyme.[12] The target strand used in this model has a length of 41 and the complement is complement is the same length. This results in 9- and 15-nt fragments that are released immediately after cis-cleavage (Reaction 3), and 26- and 32-nt fragments that are retained in the active enzyme and released only upon inactivation (Reaction 4). The ANA considered here is has a length of 52 nucleotides and is cleaved into approximately 26-nt strands (Supplementary Table 2). After cis-cleavage of T2, the released fragment has a length of 4 nucleotides (Reaction 7) and the 22-nt fragment is retained until inactivation (Reaction 8). Cis-cleavage of the melted ANA releases a fragment with a length of 30 nucleotides (Reaction 9) and retains the same 22-nt fragment.
For component degradation during trans-cleavage, the single-stranded DNA species are assumed to be cleaved starting from the end of the strand, as discussed in the main text. This results in one small fragment (3 nucleotides) and one large fragment (variable length, depending on the species). The variable fragments have lengths of 23, 49, 38, 23, and 2 nucleotides based on the length of T2C, $ANA_{melted}$, T1, T2, and Q, respectively (Reactions 15-24). The degradation of reaction components is enabled by default in the code but can be disabled by setting the constant nonspecific_degradation to false.
The RNA cleavage during RNP complex formation is not considered, as ssRNA is not considered a trans-cleavage inhibitor in this model.

**Detailed Reaction Information**
The following describes the rationale for the reaction list:
RNP Complex Formation
RNP complexes must be formed as the enzyme processes the gRNA (reactions 1 and 2). This happens via a two-step process of pre-gRNA binding and cleavage to form the mature RNP complex.[7] In our model, this reaction serves to convert the enzyme ($E$) to the inactive RNP complex ($E1_{inact}$ and $E2_{inact}$). Sinan et al. measured this process in AsCas12a and found that the binding was extremely tight, with a dissociation constant of approximately 0.6 pM. This is significantly stronger than previously reported dissociation constants, which used less sophisticated measurement techniques.[13,14] The constants reported by Sinan et al. indicate that the characteristic time for the binding is on the order of seconds

when the concentration of the limiting species is on the order of 10 nM (typical of one-pot assays) and less than a second when the concentration is on the order of 1 µm (typical of assays with a separate complex formation step). The half-life for the cleavage, on the other hand is approximately 5 minutes. Once the mature gRNA is formed, the authors found that the binding kinetics were essentially unchanged. The measured constants indicate a dissociation half-life of approximately five days. They also found similar kinetics in LbCas12a and FnCas12a, indicating that this behavior is conserved among Cas12a enzymes, perhaps to ensure the gRNA remains bound while the enzyme searches for the target.

Because it is common to pre-complex the enzyme and gRNA, forming mature gRNA before the assay begins, we ignore the cleavage step in this model and consider only the binding kinetics. We also ignore the production of cleaved ~16-20-nt single-stranded RNA during gRNA processing, although this may be a consideration in Cas13-based assays where these could act as reaction inhibitors. Sinan et al. also observed little effect of sequence variation in the guide region on binding, indicating that the kinetics do not vary widely for properly designed gRNA. This is consistent with the observation by Moon and Liu that a full-sized gRNA did not significantly displace an already-processed one.[15] For this reason, we do not consider any affinity difference for gRNA1 and gRNA2.

Cis-cleavage

The next mechanism is cis-cleavage (reactions 3, 7, and 9), which is responsible for converting the inactive enzyme ($E1_{inact}$ and $E2_{inact}$) into its active, trans-cleaving form ($E1_{act}$ and $E2_{act}$). Cis-cleavage can be modeled as a Michaelis-Menten process and has been reported to occur with typical reaction time scales on the order of 100 seconds at low target concentrations.[16] Consequently, cis-cleavage time has been considered as negligible in single-CRISPR-Cas assays, where processing occurs primarily at the start of the reaction and is rapid compared to the overall reaction time scale. However, cis-cleavage in the autocatalytic system continues throughout the reaction as additional ANA is processed, so the effect of the cis-cleavage rate is far more significant. Because the time scale of autocatalytic reactions is much shorter, the initial target cis-cleavage step may also make up a significant portion of the reaction. Strohkendl et al. reported $k_{on}$ and $k_{off}$ values of $1.1 \times 10^8\ M^{-1}s^{-1}$ and $5.9 \times 10^{-6}\ s^{-1}$, respectively, for AsCas12a.[8] Further, they reported that target strand cleavage occurred with a $k_{cat}$ value of $5.1 \times 10^{-3}\ s^{-1}$, 10-fold slower than non-target strand cleavage. CRISPR-Cas-based systems utilizing Cas12 have been designed with double-stranded targets containing a protospacer adjacent motif (PAM) and single-stranded targets, where a PAM region is unnecessary.[10,17] In either case, the target strand cleavage is rate-limiting. Thus, we consider a single catalysis step with the rate reported for target strand cleavage.

Trans-cleavage

The central mechanism of this reaction is trans-cleavage, which is responsible for reactions 10-13 and 15-24. Trans-cleavage occurs after activation of the enzyme and all single-stranded DNA (ssDNA) species in the reaction are considered as substrates. While single-stranded RNA (ssRNA) is also reported to be cleaved by Cas12a, the rate is significantly lower than ssDNA.[18] Because of the reduced rate and the significantly reduced presence of ssRNA in the reaction modeled here, this mechanism is neglected. This is the key process determining the performance of single-CRISPR-Cas diagnostics.[16] Because of this, Huyke et al. measured the trans-cleavage $K_M$ and $k_{cat}$ values of many Cas12 and Cas13 variants, finding that $K_M$ fell between the orders of 100 and 1000 nM and $k_{cat}$ fell between the orders of 0.01 and 1 $s^{-1}$. This aligns with our measured values of $K_M = 294\ nM$ and $k_{cat} = 0.136\ s^{-1}$ (Supplementary Figure 1, protocol in Supplementary Note 2) which are used by default in the model. To determine the exact rate constants, it is necessary to provide a forward or reverse rate constant for trans-cleavage. The other constant can then be calculated using the relationship:

$$K_M = \frac{k_r + k_{cat}}{k_f} \qquad (SN1-1)$$

To the authors' knowledge, no value has been directly reported for either the forward or reverse constant. A reasonable upper bound for the forward rate is $\sim 10^{10}\ M^{-1}s^{-1}$, which is near the maximum value reported for diffusion-limited enzymes.[19] For $K_M$ on the order of 100 nM and $k_{cat}$ on the order of 0.1 $s^{-1}$, according to equation SN1-1, $k_r \approx k_{cat}$ when $k_f \approx 2 \times 10^6\ M^{-1}s^{-1}$. For values of $k_f$ significantly greater than this:

$$k_r = K_M k_f - k_{cat} \approx K_M k_f \qquad (SN1-2)$$

So:

$$K_M \approx K_D = \frac{k_r}{k_f} \qquad (SN1-3)$$

This is the rapid equilibrium regime, and the approximate trans-cleavage rate:

$$v = \frac{k_{cat}E_0[S]}{K_M + [S]} \approx \frac{k_{cat}E_0[S]}{K_D + [S]} \qquad (SN1-4)$$

depends only on the ratio of $k_f$ to $k_r$ (assuming the total enzyme $E_0$ and substrate $[S]$ concentrations are maintained), showing little effect of the individual choice of the forward and reverse rates. Thus, values between $10^6$ and $10^{10}$ should behave similarly in terms of the trans-cleavage rate. For our simulation, we select a value of $10^8\ M^{-1}s^{-1}$, approximately the middle of this regime when $10^6\ M^{-1}s^{-1} < k_f < 10^{10} M^{-1}s^{-1}$. This aligns roughly with the forward rate reported for cis-cleavage. Cas12 kinetics vary based on the thermodynamic parameters of the target and guide nucleic acids, but this effect is minor when the targets are designed within an optimal free energy range.[20] While the guide and target selection affect the kinetics, the base composition of the trans-cleavage target do not vary significantly depending on the substrate.[18] Thus, we assume that the trans-cleavage kinetics do not vary significantly for different substrates. Reactions 15-24 are modeled based on the same trans-cleavage kinetics.

<u>Enzyme Inactivation</u>

Once activated, it is possible for the enzyme to become inactive (reactions 4 and 8) via the dissociation of target DNA with a half-life of approximately 32 hours.[8] This process is modeled as irreversible. For an irreversible reaction:

$$k_{inact} = \frac{\ln(2)}{t_{1/2}} = 6 \times 10^{-6}\ s^{-1} \qquad (SN1-5)$$

where $t_{1/2}$ is the reaction half-life, in seconds. This reaction is negligible over the time course of most diagnostic applications.

<u>Non-specific reporter degradation</u>

Avaro et al. observed that the single-stranded reporters used frequently in CRISPR-Cas systems undergo enzyme-independent degradation, which is important in low-background assays.[9] This can be modeled as an irreversible process (reaction 14), with $k_{rep} = 1.4 \times 10^{-6} s^{-1}$.

<u>ANA Equilibrium</u>

The final major reactions (reactions 5 and 6) relate to the equilibrium between the blocked and unblocked autocatalytic nucleic acid (ANA) molecule before and after cleavage. This is the most significant parameter in determining assay performance and a major point of optimization, as the design of the ANA can vary widely and is almost entirely independent of the target. Each of these reactions are modeled as a simple two-state equilibrium, with the forward and reverse rates determined by the change in free energy for annealing, which can be modeled or measured. While the complement strand produced by ANA cleavage acts as a ssDNA inhibitor, it is treated separately so that the equilibrium of the cleaved ANA can be tracked properly. The inhibitory effect is captured in its trans-cleavage degradation (reactions 15 and 16).

In practice, the uncleaved ANA equilibrium can be estimated simply using a melting curve. However, the derivative method used commonly to estimate the melting temperature ($T_m$) is not sufficient for determining the value of $\Delta G°$, which is necessary to calculate the rate constants (Supplementary Note 3).[3] This method assigns the melting temperature as the peak of the derivative of the melting curve, which should roughly correspond to the point where the nucleic acid is 50% annealed in a two-state equilibrium. However, it does not derive any information about the rate of change throughout the rest of the melting curve. Consequently, the derivative method can approximate the temperature at which 50% is annealed, but not the fraction annealed at any other temperature. Because the assay temperature tends to be significantly lower than $T_m$ for the ANA used in these assays, this does not give reliable information about the equilibrium at the assay temperature. The method described by Mergny and Lacroix instead utilizes the entire melting curve to estimate the value of the standard enthalpy change ($\Delta H°$) and standard entropy change ($\Delta S°$) of the reaction, resulting in a temperature-dependent value for the standard Gibbs free energy change:

$$\Delta G°(T) = \Delta H° - T\Delta S°$$

which allows for estimation of the equilibrium away from the melting temperature.[3] Notably, because this method is based on the van't Hoff equation, it assumes that the melting and annealing follow a certain established equilibrium (such as $A \leftrightarrow B$) and that $\Delta H°$ and $\Delta S°$ are temperature independent. The validity of these assumptions may be tenuous far from the melting temperature, so the implied values derived in these cases should be interpreted cautiously. Melting curves for a simple hairpin and bulge ANA, as well as estimation of these thermodynamic parameters, are provided in Supplementary Figures 8 and 9 and show reasonable agreement for the fits obtained from the measured and simulated data. Measurement of these parameters for cleaved ANA molecules is complicated by the presence of off-target cleavage products after exposure to the nuclease. Certain strands may be purified for melting curve analysis via electrophoretic methods, but the isolated products no longer accurately reflect the strands present in the actual assay, so the value of this data is limited.

To determine the forward and reverse rates exactly, it is necessary to obtain some information about the time scale at which the nucleic acid moves towards its equilibrium state. This can be achieved with perturbation and relaxation experiments, typically by creating a step change in either the temperature or the concentration of a certain species and observing the return to equilibrium. The characteristic time of this return for small hairpins (4-6 nt loop and <10 bp stem) has been reported to be as short as microseconds.[21] However, others have argued that millisecond time scales are necessary for formation of complete hairpins, with the microsecond transitions reflecting intermediate states.[22] The default time scale chosen for this model is milliseconds to match the more conservative figures. The exact formulation used to obtain the forward and reverse rate constants here are detailed in Supplementary Note 3. In practice, the functions **calculateEquilibriumRatesUnimolecular** and **calculateEquilibriumRatesBimolecular** are provided to approximate the forward and reverse rate constants of single-strand and dual-strand ANAs, respectively, based on the standard free energy change and time constant.

#### Supplementary Note 2: Kinetic Experimental Protocols

##### Materials

The Cas12 enzyme (Alt-R LbCas12a (Cpf1) Ultra) was obtained from Integrated DNA Technologies (IDT). All nucleic acid sequences and the reporter were synthesized by IDT and resuspended in Tris-EDTA (TE) buffer as specified by IDT. ROX Passive Reference Dye (12223012) and UltraPure 1M Tris-HCl Buffer (15567027) and NaCl (5M, RNase-free, AM9760G) were purchased from ThermoFisher Scientific. Magnesium Chloride ($MgCl_2$) Solution (B9021S) and NEBuffer r2.1 (B6002S) were purchased from New England Biolabs. Nuclease-free water (NFW, 11-05-01-04) was purchased from IDT. EvaGreen Dye (20X, 31000) was purchased from Biotium.

##### Sequences

| Strand Name | Sequence (5'→3', Target Region Underlined) | Length (nt) |
|---|---|---|
| gRNA | UAAUUUCUACUAAGUGUAGAUGGAUAUACGAUAUAUAUAUAU | 42 |
| Target | AGGGGCGTGCATATATATATATCGTATATCCGTTGGTGACG | 41 |
| Inhibitor | ACTTTAGATCGTTATATAACGCTAACTATGA | 31 |
| Reporter | /56-FAM/TTTTTT/3IABkFQ/ | 6 |

##### Standard Michaelis-Menten Experiment

RNPT1 mix was prepared according to:

| RNPT1 Mix | | | |
|---|---|---|---|
| Reagent | [Stock] (M) | [Desired] (M) | Volume (μL) |
| Cas12 | 5.00E-07 | 5.00E-09 | 2.00 |
| gRNA | 5.00E-07 | 6.25E-09 | 2.50 |
| MgCl2 | 2.50E-02 | 5.00E-03 | 40.00 |
| NFW | | | 155.50 |

and incubated for 20 minutes at room temperature. A serial dilution of 1X, 0.5X, 0.25X, 0.125X, and 0.0625X reporter was created (1000 nM-62.5 nM final concentration). A reaction mix was prepared with the following components (temporarily omitting the reporter and RNPT1 mix):

| Reaction Mix (Total) | | | |
|---|---|---|---|
| Reagent | [Stock] (M) | [Desired] (M) | Volume (μL) |
| Reporter | 5.00E-06 | 1.00E-06 | ~~60.00~~ |
| ROX | 5.00E-05 | 5.00E-07 | 3.00 |
| NEB Buffer | 1.00E+01 | 1.00E+00 | 30.00 |
| MgCl2 | 2.50E-02 | 2.50E-03 | 30.00 |
| RNPT1 Mix | 5.00E-09 | 1.00E-09 | ~~60.00~~ |
| Target | 1.00E-07 | 5.00E-09 | 15.00 |
| NFW | | | 102.00 |

The reaction mix was split into five tubes, at which point the reporter dilution was added followed by the RNPT1 mix. Each tube was split into four 10 μL replicates and transferred to a QuantStudio 3 Real-Time PCR System (ThermoFisher Scientific). It was held at 37°C for 4 hours, with measurements taken every 30 seconds. Data was processed according to the previously described normalization and calibration procedure to obtain cleaved reporter concentrations.

**Excess Target Experiment**
The RNPT1 mix was prepared with the same method as the standard Michaelis-Menten experiment. A serial dilution of target was created with concentrations of 10 µM, 1 µM, 100 nM, and 10 nM for later dilution to final concentrations of 1 µM, 100 nM, 10 nM, and 1 nM. The reaction mix was prepared with the following components (temporarily omitting the target and RNPT1 mix):

| Reaction Mix (Total) | | | |
|---|---|---|---|
| **Reagent** | **[Stock] (M)** | **[Desired] (M)** | **Volume (µL)** |
| Reporter | 5.00E-06 | 5.00E-07 | 25.00 |
| ROX | 5.00E-05 | 5.00E-07 | 2.50 |
| NEB Buffer | 1.00E+01 | 1.00E+00 | 25.00 |
| MgCl2 | 2.50E-02 | 2.50E-03 | 25.00 |
| RNPT1 Mix | 5.00E-09 | 1.00E-09 | ~~50.00~~ |
| Target | 1.00E-05 | 1.00E-06 | ~~25.00~~ |
| NFW | | | 97.50 |

The reaction mix was split into four group and the target dilution was added for each group. Finally, the RNPT1 mix was added and each group. Each group was split into four 10 µL replicates and transferred to a QuantStudio 3 Real-Time PCR System (ThermoFisher Scientific). It was held at 37°C for 4 hours, with measurements taken every 30 seconds. Data was processed according to the previously described normalization and calibration procedure to obtain cleaved reporter concentrations.

**Inhibition Experiment**
RNPT1 mix was prepared according to:

| RNPT1 Mix | | | |
|---|---|---|---|
| **Reagent** | **[Stock] (M)** | **[Desired] (M)** | **Volume (µL)** |
| Cas12 | 5.00E-07 | 5.00E-09 | 3.00 |
| gRNA | 5.00E-07 | 6.25E-09 | 3.75 |
| MgCl2 | 2.50E-02 | 5.00E-03 | 60.00 |
| NFW | | | 233.25 |

and incubated at room temperature. Two serial dilutions were created. The inhibitor was diluted into four tubes of 10 µM, 5 µM, 2.5 µM, and 1.25 µM, for later dilution to final concentration of 1 µM, 500 nM, 250 nM, and 125 nM. The reporter was similarly diluted into five tubes of 5 µM, 2.5 µM, 1.25 µM, 625 nM, and 312.5 nM for later dilution to a final concentration of 500 nM, 250 nM, 125 nM, 62.5 nM, and 31.25 nM. A reaction mix was prepared with the following components (temporarily omitting the inhibitor, reporter, and RNPT1 mix):

| Reaction Mix (Total) | | | |
|---|---|---|---|
| **Reagent** | **[Stock] (M)** | **[Desired] (M)** | **Volume (µL)** |
| Reporter | 5.00E-06 | 1.00E-06 | ~~260.00~~ |
| ROX | 5.00E-05 | 5.00E-07 | 13.00 |
| NEB Buffer | 1.00E+01 | 1.00E+00 | 130.00 |
| MgCl2 | 2.50E-02 | 2.50E-03 | 130.00 |
| RNPT1 Mix | 5.00E-09 | 1.00E-09 | ~~260.00~~ |
| Target | 1.00E-07 | 1.00E-09 | 13.00 |
| Inhibitor | 1.00E-05 | 1.00E-06 | ~~130.00~~ |
| NFW | | | 364.00 |

The reaction mix was then distributed into 20 tubes and reporter and inhibitor serial dilutions were added to obtain each combination of the two concentrations. The samples were placed on ice to slow any reaction and then the RNPT1 mix was added to initiate the reaction. Each tube was split into four 10 µL replicates while on ice and then transferred to a QuantStudio 3 Real-Time PCR System. It was held at 37°C for 4 hours, with measurements taken every 30 seconds. Data was processed according to the previously described normalization and calibration procedure to obtain cleaved reporter concentrations.

**Melting Curve Measurement**

For melting curve measurements, the ANA mixtures were prepared according to:

| ANA Mix (Hairpin) | | | |
|---|---|---|---|
| **Reagent** | **[Stock] (M/X)** | **[Desired] (M/X)** | **Volume (µL)** |
| SYBR | 2.00E+01 | 1.00E00 | 2.50 |
| ANA | 1.00E-05 | 1.00E-06 | 5.00 |
| Tris HCl | 1.00E+00 | 1.08E-02 | 0.54 |
| NaCl | 5.00E+00 | 5.20E-02 | 0.52 |
| MgCl2 | 2.50E-02 | 1.39E-02 | 27.80 |
| NFW | | | 13.64 |

| ANA Mix (Bulge) | | | |
|---|---|---|---|
| Complement-to-Target Ratio | | 1.2 | |
| **Reagent** | **[Stock] (M/X)** | **[Desired] (M/X)** | **Volume (µL)** |
| EvaGreen | 2.00E+01 | 1.00E00 | 2.50 |
| ANA Target | 1.00E-05 | 1.00E-06 | 5.00 |
| ANA Complement | 1.00E-05 | 1.20E-06 | 6.00 |
| Tris HCl | 1.00E+00 | 1.08E-02 | 0.54 |
| NaCl | 5.00E+00 | 5.20E-02 | 0.52 |
| MgCl2 | 2.50E-02 | 1.39E-02 | 27.80 |
| NFW | | | 7.60 |

The samples were split into four 10 µL replicates and placed in a QuantStudio 7 Pro Real-Time PCR System (ThermoFisher Scientific). The samples were heated to 95°C, held for 4 minutes, and then transitioned to 20°C at a rate of 0.1°C/s and held there for 20 seconds. For the melting curve, the sample was heated from 20°C to 95°C at a rate of 0.02°C/s, with measurements taken approximately every 0.1°C.

#### Supplementary Note 3: ANA Rate Constants

**Unimolecular ANA**

For a unimolecular ANA described by the equilibrium:

$$ANA_{melted} \rightleftharpoons ANA_{annealed}$$

we can assume that we start with a concentration of ANA ($[ANA]_{tot}$) that is entirely the melted state to obtain:

$$\frac{d[ANA]_{annealed}}{dt} = k_f[ANA_{melted}] - k_r[ANA_{annealed}] = k_f[ANA]_{tot} - (k_f + k_r)[ANA]_{annealed}$$

which results in a solution of the form:

$$[ANA_{annealed}](t) = \frac{k_f[ANA]_{tot}}{k_f + k_r}\left(1 - e^{-(k_f+k_r)t}\right)$$

This is a first-order exponential relaxation with the time constant:

$$\tau = \frac{1}{k_f + k_r}$$

More generally, for the system returning to equilibrium from any perturbation, the form is:

$$[ANA_{annealed}](t) = \frac{k_f[ANA]_{tot} + k_r[ANA_{annealed}]_0 e^{-(k_f+k_r)t}}{k_f + k_r}$$

where $[ANA_{annealed}]_0$ is the initial concentration of $[ANA_{annealed}]$ after the perturbation. If we assume that the reaction has returned a certain fraction of the way to equilibrium $\beta$ ($\beta \in [0,1]$, where 0 is the perturbed state and 1 is the equilibrium state) within a time $t_\beta$, we can solve:

$$\beta = 1 - e^{-(k_f+k_r)t_\beta}$$

to obtain:

$$k_f + k_r = k_{tot} = -\frac{\ln(1-\beta)}{t_\beta}$$

From the equilibrium constant for the reaction:

$$K_{eq} = \frac{k_f}{k_r} = e^{-\Delta G^\circ_{anneal}/RT}$$

we can restate:

$$k_f = \left(-\frac{\ln(1-\beta)}{t_\beta}\right)\Big/\left(1 + \frac{1}{K_{eq}}\right) = \frac{K_{eq}}{1 + K_{eq}}k_{tot}$$

$$k_r = \frac{1}{1 + K_{eq}}k_{tot}$$

This is how the forward and reverse rate constants are calculated in the **calculateEquilibriumRatesUnimolecular** function.

**Bimolecular ANA**

The process for estimating the rate constants for a bimolecular equilibrium is more complex. For a bimolecular ANA (formed by association of a target strand $T$ and compliment strand $C$) with the equilibrium:

$$ANA \rightleftharpoons T + C$$

the rate of change for $ANA$ is:

$$\frac{d[ANA]}{dt} = k_r[T][C] - k_f[ANA]$$

In the case where we start with fully melted nucleic acids at equal concentration ($[T]_{tot} = [C]_{tot}$):

$$[T] = [T]_{tot} - [ANA]$$
$$[C] = [C]_{tot} - [ANA]$$

Then:

$$\frac{d[ANA]}{dt} = k_r([T]_{tot} - [ANA])([C]_{tot} - [ANA]) - k_f[ANA]$$
$$= k_r([T]_{tot} - [ANA])^2 - k_f[ANA]$$

Unlike the one-component case, this is non-linear. To create a linear form, we need to assume a near-equilibrium perturbation where the behavior of the system is approximately linear. At a point near equilibrium ($\delta[ANA] \ll ANA_{eq}$):

$$[ANA] = [ANA]_{eq} + \delta[ANA]$$

the rate of change for this quantity is:

$$\frac{d\left([ANA]_{eq} + \delta[ANA]\right)}{dt} = k_r\left([T]_{tot} - [ANA]_{eq} - \delta[ANA]\right)^2 - k_f\left([ANA]_{eq} + \delta[ANA]\right)$$

The squared term can be expanded as:

$$\begin{aligned}\left([T]_{tot} - [ANA]_{eq} - \delta[ANA]\right)^2 &= [T]_{tot}^2 - [ANA]_{eq}[T]_{tot} - [T]_{tot}(\delta[ANA]) - [ANA]_{eq}[T]_{tot} + [ANA]_{eq}^2 + [ANA]_{eq}(\delta[ANA]) \\ &\quad - (\delta[ANA])[T]_{tot} + (\delta[ANA])[ANA]_{eq} + (\delta[ANA])^2 \\ &= \left([T]_{tot} - [ANA]_{eq}\right)^2 - 2\left([T]_{tot} - [ANA]_{eq}\right)(\delta[ANA]) + (\delta[ANA])^2\end{aligned}$$

Removing the non-linear term $(\delta[ANA])^2$ and substituting results in:

$$\begin{aligned}\frac{d\left([ANA]_{eq} + \delta[ANA]\right)}{dt} &= \frac{d(\delta[ANA])}{dt} \\ &= k_r\left([T]_{tot} - [ANA]_{eq}\right)^2 - 2k_r\left([T]_{tot} - [ANA]_{eq}\right)(\delta[ANA]) - k_f[ANA]_{eq} - k_f(\delta[ANA])\end{aligned}$$

From the original rate, we can say that exactly at equilibrium:

$$\frac{d[ANA]}{dt} = 0 = k_r\left([T]_{tot} - [ANA]_{eq}\right)^2 - k_f[ANA]_{eq}$$

or:

$$k_r\left([T]_{tot} - [ANA]_{eq}\right)^2 = k_f[ANA]_{eq}$$

Therefore:

$$\frac{d(\delta[ANA])}{dt} = -\left(2k_r\left([T]_{tot} - [ANA]_{eq}\right) + k_f\right)(\delta[ANA])$$

This results in:

$$(\delta[ANA])(t) = (\delta[ANA])(0)e^{-k_{tot}t}$$

where:

$$k_{tot} = 2k_r\left([T]_{tot} - [ANA]_{eq}\right) + k_f$$

and:

$$\tau = \frac{1}{2k_r\left([T]_{tot} - [ANA]_{eq}\right) + k_f}$$

As in the unimolecular ANA case, we will consider:

$$k_{tot} = -\frac{\ln(1-\beta)}{t_\beta}$$

and:

$$K_c = \frac{k_f}{k_r} = e^{\Delta G^\circ_{anneal}/RT}$$

Note that the free energy change here is the opposite of the unimolecular case because the equilibrium is defined with the annealed ANA on the left.

In this case, the equilibrium constant is defined as:

$$K_{eq} = \frac{a_T a_C}{a_{ANA}} = \frac{[T]_{eq}[C]_{eq}}{[ANA]_{eq}}/C^\circ \rightarrow K_c = \frac{[T]_{eq}[C]_{eq}}{[ANA]_{eq}} = K_{eq}C^\circ$$

and the standard state concentration $C^\circ$ is 1 M. Consequently:

$$k_f = \frac{K_c}{K_c + 2\left([T]_{tot} - [ANA]_{eq}\right)}k_{tot}$$

$$k_r = \frac{1}{K_c + 2\left([T]_{tot} - [ANA]_{eq}\right)}k_{tot}$$

Thus, finding the forward and reverse rates requires solving for the equilibrium concentration of ANA. Because we started with equal concentrations of $T$ and $C$, we can solve for the equilibrium concentrations with the quadratic:

$$K_c = \frac{[T]_{eq}[C]_{eq}}{[ANA]_{eq}} = \frac{\left([T]_0 - [ANA]_{eq}\right)^2}{[ANA]_{eq}}$$

$$[ANA]_{eq}{}^2 - (2[T]_0 + K_c)[ANA]_{eq} + [T]_0^2 = 0$$

$$[ANA]_{eq} = \frac{2[T]_0 + K_c \pm \sqrt{(2[T]_0 + K_c)^2 - 4[T]_0^2}}{2} = \frac{2[T]_0 + K_c \pm \sqrt{K_c(4[T]_0 + K_c)}}{2}$$

The root can be simplified as such:

$$\sqrt{(2[T]_0 + K_c)^2 - 4[T]_0^2} = \sqrt{K_c(4[T]_0 + K_c)}$$

This quantity is always greater than zero because both $K_c$ and $[T]_0$ are positive. As a result, when the positive root is used

$$[ANA]_{eq} = [T]_0 + \frac{K_c \pm \sqrt{K_c(4[T]_0 + K_c)}}{2} > [T]_0$$

This is not physically reasonable for our conditions, so only the negative root need to be considered:

$$[ANA]_{eq} = \frac{2[T]_0 + K_c - \sqrt{K_c(4[T]_0 + K_c)}}{2}$$

The solution obtained from this root can be substituted in the equations for the forward and reverse rate constants. This is exactly the approach used in the **calculateEquilibriumRatesBimolecular** function. Because the time constant here depends on the concentration, this function must take a concentration value as an input; for approximating the rate constants, the total ANA concentration present in the reaction is a reasonable choice. The values of the forward and reverse rate constants are not particularly sensitive to changes in concentration within the same order of magnitude.

This process can be expanded to an n-component system, but this involves solving an nth degree polynomial for the equilibrium concentration. The near-equilibrium linear approximation used here is based on the method demonstrated in [23].

#### Supplementary Note 4: Model Structure

##### Model Setup and Operation

In the project directory, there are two major types of files: experiment functions and ordinary differential equation (ODE) reaction scheme functions. In addition, there is a private sub-directory with several helper functions

Each reaction scheme requires an ODE generator function (e.g. **ODEGeneratorHairpin(**user_constants**)**) for the basic hairpin ANA reaction) that defines the individual reactions, rate constants, and fragmentation behavior for a particular reaction scheme. The individual reactions are defined as a cell array of reaction strings, with reaction strings in the format:

```
'12) E1_{act} + FQ <-> E1_{act}FQ -> E1_{act} + F + Q’
```

Reactions may be reversible, irreversible, or Michaelis-Menten type. There are two functions (**buildSpeciesList**, **buildODEExpressions**) that convert the reaction cell array into a series of ODEs. Each reaction must have an accompanying set of rate constants. The functions **calculateEquilibriumRatesUnimolecular** and **calculateEquilibriumRatesBimolecular** are available to assist in calculating the equilibrium rate constants for one-component and two-component ANA, respectively. **calculateAlphaUnimolecular**, **calculateAlphaBimolecular**, and **calculateAlphaNmolecular** are available for calculating the $\alpha$ values for ANA, although this is not used directly in the ODE solver. **buildAffinityVector** and **buildFragmentationMatrix** are used to calculate the coefficients used in the single-stranded DNA fragmentation process. The ODE generator function takes a user_constants parameter that allows the user to temporarily modify the constants used in the ODE generator function without modifying the ODE generator function directly. The output of the ODE generator function is an ODE object that can be used with MATLAB ODE solvers like ode15s.

To run an experiment with the model, the user first generates the ODE function. This is done by calling the desired ODE generator function, such as:

[ode_function,names] = ODEGeneratorHairpin();

To modify a constant such as the ANA standard free energy change, the ODE function is called with the user_constants argument:

```
mod_constants.dG_uncleaved = -14;   % kcal/mol
mod_constants.dG_cleaved = -10; % kcal/mol
[ode_function,names] = ODEGeneratorHairpin(mod_constants);
```

Other constants that can be modified include the cis- and trans-cleavage rate constants, the size range of simulated DNA fragments, and whether, non-specific degradation of reaction components should be considered. The full list of constants can be observed in the ODE generator function. ODE solver options must then be generated. These set the behavior of the MATLAB ODE solver. Throughout the experiments used in this manuscript, the options are set as:

```
min_conc = 1e-21; % minimum species concentration that needs to be considered
tol_mod = 1; % 1 default
min_threshold = tol_mod*min_conc;   %meaningful value cutoff
model_options = odeset('NonNegative',1:numel(y0),'RelTol', 1e-9*tol_mod,
'AbsTol', min_threshold);
```

These options are found to generally result in stable solutions and can maintain species concentrations down to approximately one molecule in a 1 mL volume.

The user then sets the initial conditions for the reaction. These are set using the **buildInitialConditions** function, which takes as an argument a Map object containing a mapping of the reaction species to their initial concentrations. If a reaction species is not specifically named, the concentration is presumed to be zero. This function outputs a vector of concentrations that is an input to the built-in MATLAB ODE solvers. For example:

```
react_init_conc = containers.Map();
react_init_conc('T1') = 1e-15; % nM
react_init_conc('FQ') = 100e-9; % nM
y0_complete = buildInitialConditions(react_init_conc,names);
```

This function can also generate the initial concentrations from multiple input tubes. For example, pre-incubation of the RNP complexes and ANA, as is typical in some experimental protocols, are simulated in the **prepareStandardExperiment** function. This simulates each incubation and returns the final concentration of each tube:

```
[y_end_preRNP1, y_end_preRNP2, y_end_preANA] =
prepareStandardExperiment(ode_function,names,model_options,plot_preinc);
```

**buildInitialConditions** can take these tube concentrations, a vector containing the dilution fractions for each tube, and a mapping of concentrations for the new species added at this step and generate the complete initial concentrations for the next step:

```
[y_end_preRNP1, y_end_preRNP2, y_end_preANA] =
prepareStandardExperiment(ode_function,names,model_options,plot_preinc);
react_init_conc = containers.Map();
react_init_conc('T1') = 1e-15; % nM
react_init_conc('FQ') = 100e-9; % nM
dilution_fractions = [0.1, 0.1, 0.04];
y0_complete =
buildInitialConditions(react_init_conc,names,{y_end_preRNP1,y_end_preRNP2,y_en
d_preANA},dilution_fractions);
```

A dilution fraction of 0.1, for example, would dilute the species present in the RNP1 tube from 100 nM to 10 nM. This allows the user to easily set up fairly complicated experiments and simulate multi-step experimental protocols.

Once the ODE generator function, model options, and initial concentrations are set, the ode function can be run. This uses a standard approach for MATLAB:

```
t_react = 30; % Reaction time (min)
t_int = 0.1;  % Time interval (min)
[t,y] = ode15s(@(t, y)
ode_function(t,y),(0:t_int:t_react)*60,y0_complete,model_options);
```

The reaction results can then be plotted using the **plotAllTabs** function, which plots a summary of the model results and several tabs tracking the concentration of individual species and single-stranded DNA fragmentation. This plotting function calls **plotReactionSummaryHairpin**, which is specific to the hairpin reaction scheme, as the name of the plotted species can change. To facilitate easy manipulation of the data from the reaction, **selectData** can isolate the data for a specific species by the name of the species, without any need for tracking the indices of the many species. For example:

```
F = selectData('F',names,y);
```

selects the cleaved reporter concentraton from the reaction data. Two helper functions, **cushionAxes** and **formatSI** are used in plotting. The process for running a basic reaction is outlined in the **basicDemo** subfunction within the **AutoCRISPRKinetics** function, which runs all of the code used to generate data and plots for this manuscript.

In addition to the scheme and helper functions described above ,there are several functions for preparing, running, and plotting multi-reaction experiments. These include **runStandardExperiment**, which runs and plots a single multi-concentration CRISPR autocatalysis reaction and **runVariableOptimization**, which runs multi-concentration experiments over a range of values for species concentrations, constants, and/or dilution fractions. The results of this can be plotted using **plotSingleVariableOptimization** or **plotDualVariableOptimization** to visualize *in silico* optimization experiments. The **test** function contains test/demonstration examples for most of the other functions. Individual demonstrations can be uncommented to demonstrate the operation of each of the functions.

For a user to create a new reaction scheme that is significantly different from the hairpin ANA scheme, they can edit a copy of **ODEGeneratorHairpin** to include the appropriate reactions and constants. If the user wants to use the **plotAllTabs** function, to visualize the reaction, a fig_struct must be created for the reaction scheme; this simply specifies the names of the tabs and the species that should be plotted in each. To match this, a copy of **plotReactionSummaryHairpin** can optionally be modified to include the most relevant species. If the summary function is omitted from the arguments for **plotAllTabs**, the summary tab will not be generated but the rest of the tabs will be generated as specified by the fig_struct.

**Function Documentation**
The function documentation is included for each of the functions used in the model and experiments. The information provided here is also provided in the function documentation and is accessible via the MATLAB help function. The full MATLAB code is provided as specified in the main text.

**Experiment Functions**

**AutoCRISPRKinetics()** – The main driver function for all experiments in the paper. The driver function calls subfunctions for each experiment. These subfunction calls can be uncommented to run a desired set of experiments.

**test()** – This file is intended for: (1) holding standard test/demo code for the various private functions and (2) allowing the user to test the various functions in the private folder, as they are not accessible from the console.

**Scheme Functions**

[ode_function,names] = **ODEGeneratorHairpin(**user_constants**)** - Generates a function capturing the CRISPR autocatalysis reaction for implementation with an ODE solver. Because the generation of these reaction systems requires many reaction-specific considerations, the function must be modified to implement other reaction systems. However, individual constants may be modified via the user_constants input.
Syntax:
[ode_function,names] = ODEGeneratorHairpin(user_constants)
Inputs:
user_constants (optional) - Struct containing independent constants to be modified from the default value defined in this function.
Outputs:
ode_function - The function generated for the reaction system, which has the syntax dydt = ode_function(t,y) for use with MATLAB ODE solvers.
names - Cell array containing the ordered names of species handled by the ODE function (as strings). This is necessary for indexing purposes and is passed to selectData and buildInitialConditions.

parent = **plotReactionSummaryHairpin(**t,y,names,fig_title,parent**)** - This function plots a summary tab of the CRISPR autocatalysis reaction described by ODEGeneratorHairpin. It is specific to the autocatalytic reaction, as other reactions may have different significant species. However, it can serve as a template for other summary functions.
Inputs:
t - Time data produced by ODE solver.
y - Species data produced by ODE solver.
names - Cell array containing the ordered names of species handled by the ODE function (as strings). This is necessary for indexing purposes and is passed to selectData and buildInitialConditions.
fig_title - Figure title as string.
parent (optional) - Parent object for figure, typically either an existing figure or tab. If not given, creates new figure.
Outputs:
parent - Figure handle or parent object if it is a tab.

**Private/Helper Functions**

These functions are contained in a private folder. As a result, they can be called from the functions in the "CRISPRAutocatalysisSimulation" directory but not from the command line.

<u>For ODE function generation:</u>
scalar_species = **buildSpeciesList(**reaction_strings,vector_species**)** – Extracts ordered list of scalar species from reaction list.

Syntax:

scalar_species = buildSpeciesList(reaction_strings,vector_species)

Inputs:

reaction_strings - Cell array of reaction strings in the format'#) A + B -> C', '#) A <-> B', or '#) A + B <->C -> D'.

vector_species (optional) - Cell array of strings with the names of the vector species, for exclusion.

Outputs:

scalar_species - Cell array of unique scalar species names (strings) in order of first appearance, excluding vector species (species matching pattern 'Name_#', like DNA fragments).

exprs = **buildODEExpressions(**reaction_strings,vector_species**)** - Builds symbolic ODE expressions for each of the species in the provided reaction strings.

Syntax:

exprs = buildODEExpressions(reaction_strings,names)

Inputs:

reaction_strings - Cell array of reaction strings in the format '#) A + B -> C', '#) A <-> B', or '#) A + B <-> C -> D'

names - Cell array of species names (strings) in order

Outputs:

exprs - Cell array of symbolic expressions (as strings) for each species' rate of change, to be evaluated by the ODE function

A=**buildAffinityVector(**max_DNA_length,min_DNA_length,type**)** - Generates affinity vector for DNA fragments of type 'flat' (for length-independent) or 'length' (for length-dependent).

Syntax:

A = buildAffinityVector(max_DNA_length,min_DNA_length,'flat')

Inputs:

max_DNA_length - Maximum DNA fragment length to model (in bases)

min_DNA_length - Minimum DNA fragment length to model (in bases)

type - String specifying affinity model:

'flat' - Length-independent binding (uniform affinity)

'length' - Length-dependent binding (affinity scales with fragment length)

Outputs:

A - Column vector of relative binding affinities for each fragment size, where the index corresponds to fragment length (max to min). The minimum length fragment always has zero affinity.

Example:

A = buildAffinityVector(10,2,'flat'); Returns: [1; 1; 1; 1; 1; 1; 1; 1; 0]

M_F = **buildFragementationMatrix(**max_DNA_length,min_DNA_length,region**)** - Generates fragmentation matrix for DNA cleavage.

Syntax:

M_F = buildFragmentationMatrix(max_DNA_length,min_DNA_length)

M_F = buildFragmentationMatrix(max_DNA_length,min_DNA_length,region)

Inputs:

max_DNA_length - Maximum DNA fragment length to model (in bases)

min_DNA_length - Minimum DNA fragment length that can be cleaved (in bases)

region (optional) - Two-element vector [n1,n2] specifying directional cleavage region. n1 and n2 are the bases at the end of the region, so an input of [2,3] will result in a single cleavage site between bases 2 and 3. If not provided or empty, assumes uniform random cleavage at any position along the fragment

Outputs:

M_F - Fragmentation matrix of size (max_DNA_length - min_DNA_length + 1) × (max_DNA_length - min_DNA_length + 1) where entry M_F(i,j) represents the probability that a fragment of length (max_DNA_length-i+1) results from cleaving a fragment of length (max_DNA_length-i+1). Each column for fragments where cleavage is able to occur sums to 2 (two fragments produced per cleavage event); otherwise, columns sum to 1

For calculating rate constants/alpha:

[k_forward,k_reverse] = **calculateEquilibriumRatesUnimolecular(**dG,beta,t_beta,T_C**)** - Calculates forward and reverse rate constants for equilibrium from free energy and timescale for reactions of the form A<->B. dG should be negative for spontaneous reaction in forward direction.

Syntax:

    [k_forward, k_reverse] = calculateEquilibriumRatesUnimolecular(dG,beta,t_beta,T_C)

Inputs:

- dG - Standard Gibbs free energy change (kcal/mol)
- beta - Fraction of equilibrium completion at time t_beta (e.g., 0.99)
- t_beta - Time to reach beta fraction of equilibrium (seconds)
- T_C - Temperature (deg Celsius)

Outputs:

- k_forward - Forward rate constant (1/s)
- k_reverse - Reverse rate constant (1/s)

[k_forward,k_reverse] = **calculateEquilibriumRatesBimolecular(**dG,beta,t_beta,T_C,C0**)** - Calculates forward and reverse rate constants for equilibrium from free energy and timescale for reactions of the form A <-> B_1 + B_2. dG should be negative for spontaneous reaction in forward direction.

Syntax:

    [k_forward, k_reverse] = calculateEquilibriumRatesBimolecular(dG,beta,t_beta,T_C,C0)

Inputs:

- dG - Standard Gibbs free energy change (kcal/mol)
- beta - Fraction of equilibrium completion by time t_beta (e.g., 0.99)
- t_beta - Time to reach beta fraction of equilibrium (seconds)
- T_C - Temperature (deg Celsius)
- C0 - Initial concentration of introduced species (approximated as [ANA]_tot)

Outputs:

- k_forward - Forward rate constant k_f (1/s)
- k_reverse - Reverse rate constant k_r (1/(M*s))

[alpha, alpha_complement] = **calculateAlphaUnimolecular(**dG,T_C**)** - Calculates equilibrium fraction in blocked state, assuming unimolecular equilibrium (e.g. A_melt <-> A_anneal). dG input should be negative for a spontaneous forward reaction. For a unimolecular reaction, alpha is concentration-independent. Both alpha and the complement must be returned to maintain numerical precision when alpha ~ 1, which is the case for highly stable binding reactions.

Syntax:

    [alpha, alpha_complement] = calculateAlpha(dG, T_C)

Inputs:

- dG - Standard Gibbs free energy change (kcal/mol)
- T_C - Temperature (deg Celsius)

Outputs:

- alpha - Equilibrium fraction in folded state (dimensionless)
- alpha_complement - Equilibrium fraction in unfolded state

[alpha, alpha_complement] = **calculateAlphaBimolecular(**dG,T_C,C_T,C_C**)** - Calculates equilibrium fraction in blocked state for bimolecular equilibrium (T + C <-> TC). dG input should be negative for a spontaneous forward reaction. Both alpha and the complement must be returned to maintain numerical precision when alpha ~ 1, which is the case for highly stable binding reactions.

Syntax:

    [alpha, alpha_complement] = calculateAlphaBimolecular(dG, T_C, C_T, C_C)

Inputs:

- dG - Standard Gibbs free energy change (kcal/mol)
- T_C - Temperature (deg Celsius)
- C_T - Initial concentration of T (M)
- C_C - Initial concentration of C (M)

Outputs:
alpha - Equilibrium fraction of T in bound state (dimensionless)
alpha_complement - Equilibrium fraction of T in unbound state (1-alpha)

[alpha, alpha_complement] = **calculateAlphaNmolecular(**dG,T_C,C_T,C_C**)** - Calculates equilibrium fraction in blocked state for n-molecular equilibrium (T + C_1 + ... C_n <-> TC). dG input should be negative for a spontaneous forward reaction. Both alpha and the complement must be returned to maintain numerical precision when alpha ~ 1, which is the case for highly stable binding reactions.

Syntax:
[alpha, alpha_complement] = calculateAlphaNmolecular(dG,T_C,C_T,C_C)

Inputs:
dG - Standard Gibbs free energy change (kcal/mol)
T_C - Temperature (deg Celsius)
C_T - Initial concentration of T (M)
C_C - Initial concentration of C_1 through C_n as vector (M)

Outputs:
alpha - Equilibrium fraction of T in bound state (dimensionless)
alpha_complement - Equilibrium fraction of T in unbound state (1-alpha)

<u>For preparing experiments and interacting with data:</u>

y0 = **buildInitialConditions(**initial_conc, names, y0_init, dilution**)** - Builds initial condition concentration vector for ODE solver.

Syntax:
y0 = buildInitialConditions(initial_conc, names, y0_init [opt], dilution [opt])

Inputs:
initial_conc - containers.Map with species names (strings) as keys and concentrations as values. Keys may contain arbitrary characters, (e.g., initial_conc('ANA_{anneal}') = 100e-9) names - Cell array of unique species names in the order expected by the ODE function (produced by the ODE generator function)
y0_init (optional) - Column vector of default initial concentrations or a cell array of multiple column vectors. If not provided, values default to zero. If multiple vectors are provided, values are summed for each species. Values in initial_conc are added to y0_init values.
dilution (optional) - Row vector of dilution fractions for initial concentrations, with length the same size as y0_init cell array. Dilution fractions (reciprocal of dilution factor) should be fractions between 0 and 1. Provided y0_init concentrations are multiplied by the corresponding dilution factor. Defaults to ones array.

Outputs:
y0 - Column vector of initial concentrations matching the order in names. Values are summed from all y0_init vectors and initial_conc.

y_mod = **selectData(**species,names,y**)** - Selects data from ode solution using species name. This enables easy selection of appropriate data without tracking species indices.

Syntax:
y_mod = selectData(species,names,y)

Inputs:
species - Cell array of names (as strings) for selected species. If name starts with $, it will search for any that start with the string. If name ends with $, it will search for any that contain that string.
names - 1xn cell array containing ordered species names as strings. Generated by the ODE generator function for a certain reaction.
y - Data output from ode solver, with each column corresponding to a species.

Outputs:
y_mod - Data for the selected species, with each column corresponding to a species.

<u>For plotting:</u>

fig = **plotAllTabs(**t,y,names,fig_title,fig_struct,plotSummaryFunction**)** - Plots tabbed layout of autocatalysis reaction. This includes a first tab with a summary, which requires a plotting function specific to the reaction. The following tabs are defined by fig_struct. The last tab shows the fragmentation information.

Syntax:

fig = plotAllTabs(t,y,fig_title,fig_struct,plotSummaryFunction)

Inputs:

t - Time vector (seconds) from ODE solver

y - Concentration matrix (M) from ODE solver, with size [timepoints x species]

names - Cell array of species names corresponding to columns of y

fig_title - String for the main figure title

fig_struct - Structure defining species tabs with fields:

.tabs - Cell array of tab names

.species - Cell array of cell arrays, each containing species name patterns (supports $ expressions as specified by selectData)

.species_labels - Cell array of cell arrays with display labels

.plot_fragmentation - Boolean for option of plotting fragmentation process

plotSummaryFunction (optional) - Function handle for first summary tab with input structure: @(t, y, names, fig_title, parent)

Outputs:

fig - Figure handle for the created tabbed plot

**cushionAxes(**ax,percent**)** - Increase y-axis limits on all y-axis rulers (left and right) by percent.

Syntax:

cushionAxes(ax,percent)

Inputs:

ax - Axis object to cushion

percent - Value (in ) to expand the axes by (+/-). Defaults to 10.

Outputs:

None

labels = **formatSI(**values, unit**)** - Converts numerical values into labels utilizing common SI prefixes.

Syntax:

labels = formatSI(values, unit)

Inputs:

values - Numerical values to be converted, as a list.

unit - Unit for values (as string), to be appended to label (e.g. 'M').

Outputs:

labels - Cell array of generated labels corresponding to values.

<u>For preparing, running, and plotting multi-reaction experiments:</u>

[y_end_preRNP1, y_end_preRNP2, y_end_preANA] =
**prepareStandardExperiment(**ode_function,names,model_options,plot_preinc,tube_options**)** - This function runs the standard pre-experimental protocol, including RNP1 complex formation, RNP2 complex formation, and ANA incubation and generates outputs representing the equilibrium concentration in each tube.

Inputs:

ode_function - The function generated for the reaction system, which has the syntax dydt = ode_function(t,y) for use with MATLAB ODE solvers.

names - Cell array containing the ordered names of species handled by the ODE function (as strings). This is necessary for indexing purposes and is passed to selectData and buildInitialConditions.

model_options - ODE solver options argument, passed directly to ODE solver. If left empty, uses default ODE options.

plot_preinc - Boolean value indicating whether to plot incubation steps.

tube_options - Optional mapping of parameters to override default times and concentrations. Paramters are t_react_preRNP1, t_react_preRNP2, t_react_preANA, n_points, E_t1, gRNA1, E_t2, gRNA2, and ANA_{tot}.

Outputs:
  y_end_preRNP1 - y data from the final time point of the RNP1 complex formation step, to be used when initiating subsequent reaction.
  y_end_preRNP2 - y data from the final time point of the RNP2 complex formation step, to be used when initiating subsequent reaction.
  y_end_preANA - y data from the final time point of the ANA annealing step, to be used when initiating subsequent reaction.

parent =
  **runStandardExperiment(**init_conc,target_concs,t_react,ode_function,names,ode_options,fig_title,fig_struct_opt,parent**)** - This function runs and plots a single experiment between multiple concentrations, as in a typical CRISPR autocatalysis experiment. Plots fraction of reporter cleaved.

Inputs:
  init_conc - Initial concentrations for non-target species. Dilution fractions can be specified as a 3x1 matrix with key 'dilution'.
  target_concs - Row vector containing the target concentrations for the experiment, typically including a negative group ([T1] = 0).
  t_react - Time period for reaction (in min).
  ode_function - The function generated for the reaction system, which has the syntax dydt = ode_function(t,y) for use with MATLAB ODE solvers.
  names - Cell array containing the ordered names of species handled by the ODE function (as strings). This is necessary for indexing purposes and is passed to selectData and buildInitialConditions.
  ode_options - ODE solver options argument, passed directly to ODE solver. If left empty, uses default ODE options.
  fig_title - Figure title as string.
  fig_struct_opt - Structure defining species tabs with fields:
    .tabs - Cell array of tab names
    .species - Cell array of cell arrays, each containing species name patterns (supports $ expressions as specified by selectData)
    .species_labels - Cell array of cell arrays with display labels
    .plot_fragmentation - Boolean for option of plotting fragmentation process
    *If fig_struct_opt is empty, individual concentration groups will not have results plotted
  parent (optional) - Parent object for figure, typically either an existing figure or tab. If not given, creates new figure.

Outputs:
  parent - Figure handle or parent object if it is a tab.

[t,data] = **runVariableOptimization(**constants,init_conc,t_react,conditions**)** - This function runs experiments over a range of values for variable inputs and returns the data for in silico optimization experiments.

Inputs:
  constants - Modified constants structure to be held constant across all experiments. If empty, uses default ode constants
  init_conc - Initial concentrations map to be held constant across all experiments. At minimum, a reporter and target concentration need to be specified here or as the variable parameter. Dilution fractions can be specified as a 3x1 matrix with key 'dilution'. Multiple target concentrations may be run by specifying init_conc('T1') as a vector of concentrations.
  t_react - Reaction time, at the end of which (by default) the fluorescence is measured (minutes).
  conditions - Structure defining experimental conditions (modifying n variables over m conditions) with fields:
    .var_types - Cell array (length n) of variable type to be modified, either 'constant','concentration', or 'dilution', as these are handled differently.
    .var_names - Cell array (length n) of variable name to be modified, as a string. (e.g. 'E_t1' or 'dG_uncleaved'). Should be empty string if var_types is 'dilution'.

.values - Cell array (length n) with col vectors (length m) of modified values. In the case of var_type = dilution, the modified values are instead in a mx3 matrix.
.labels (unused) - Cell array (length n) of labels for each condition.

Outputs:
t - Vector containing time points (in min) where ode was evaluated.
data - Array of structures containing the data generated by the experiment. The size of the array is the number of conditions (n). Each structure in the array contains the fields:
.data_matrix_pos - Matrix output from ode solver with dimension num_points x num_species x num_target_concs for positive sample
.data_matrix_neg - Matrix output from ode solver with dimension num_points x num_species for negative sample
.names - Cell array containing the ordered names of species handled by the ODE function (as strings).

parent = **plotSingleVariableOptimization(**conditions,t,data,t_sample,fig_title,fig_struct_opt,parent**)** - This function plots the results of runVariableOptimization over a single axis (corresponding to each condition) for in silico optimization experiments.

Inputs:
conditions - Structure defining experimental conditions (modifying n variables over m conditions) with fields:
.var_types - Cell array (length n) of variable type to be modified, either 'constant','concentration', or 'dilution', as these are handled differently.
.var_names - Cell array (length n) of variable name to be modified, as a string. (e.g. 'E_t1' or 'dG_uncleaved'). Should be empty string if var_types is 'dilution'.
.values - Cell array (length n) with col vectors (length m) of modified values. In the case of var_type = dilution, the modified values are instead in a mx3 matrix.
.labels - Cell array (length n) of labels for each condition.
t - Vector containing time points (in min) where ode was evaluated.
data - Array of structures containing the data generated by the experiment. The size of the array is the number of conditions (n). Each structure in the array contains the fields:
.data_matrix_pos - Matrix output from ode solver with dimension num_points x num_species x num_target_concs for positive sample
.data_matrix_neg - Matrix output from ode solver with dimension num_points x num_species for negative sample
.names - Cell array containing the ordered names of species handled by the ODE function (as strings)
t_sample - The time at which the signal and SNR is calculated, in min. If not empty, defaults to last time point.
fig_title - Figure title as string.
fig_struct_opt - Structure defining species tabs with fields:
.tabs - Cell array of tab names
.species - Cell array of cell arrays, each containing species name patterns (supports $ expressions as specified by selectData)
.species_labels - Cell array of cell arrays with display labels
.plot_fragmentation - Boolean for option of plotting fragmentation process
*If fig_struct_opt is empty, individual concentration groups will not have results plotted
parent (optional) - Parent object for figure, typically either an existing figure or tab. If not given, creates new figure.

Outputs:
parent - Figure handle or parent object if it is a tab.

parent = **plotDualVariableOptimization(**conditions,t,data,t_sample,fig_title,fig_struct_opt,parent**)** - This function plots the results of runVariableOptimization over two axes (corresponding to each condition) for in silico optimization experiments.

Inputs:

conditions - Structure defining experimental conditions (modifying n variables over m conditions) with fields:
 .var_types - Cell array (length n) of variable type to be modified, either 'constant','concentration', or 'dilution', as these are handled differently.
 .var_names - Cell array (length n) of variable name to be modified, as a string. (e.g. 'E_t1' or 'dG_uncleaved'). Should be empty string if var_types is 'dilution'.
 .values - Cell array (length n) with col vectors (length m) of modified values. In the case of var_type = dilution, the modified values are instead in a mx3 matrix.
 .labels - Cell array (length n) of labels for each condition.
t - Vector containing time points (in min) where ode was evaluated.
data - Array of structures containing the data generated by the experiment. The size of the array is the number of conditions (n). Each structure in the array contains the fields:
 .data_matrix_pos - Matrix output from ode solver with dimension num_points x num_species x num_target_concs for positive sample
 .data_matrix_neg - Matrix output from ode solver with dimension num_points x num_species for negative sample
 .names - Cell array containing the ordered names of species handled by the ODE function (as strings)
t_sample - The time at which the signal and SNR is calculated, in min. If not empty, defaults to last time point.
fig_title - Figure title as string.
fig_struct_opt - Structure defining species tabs with fields:
 .tabs - Cell array of tab names
 .species - Cell array of cell arrays, each containing species name patterns (supports $ expressions as specified by selectData)
 .species_labels - Cell array of cell arrays with display labels
 .plot_fragmentation - Boolean for option of plotting fragmentation process
 *If fig_struct_opt is empty, individual concentration groups will not have results plotted
parent (optional) - Parent object for figure, typically either an existing figure or tab. If not given, creates new figure.

Outputs:
 parent - Figure handle or parent object if it is a tab.

#### Supplementary Note 5: ANA Thermodynamics

**Background**
Chemical equilibria are typically described by $K_{eq}$, which is intrinsic to a reaction for a certain temperature. However, physical operation of CRISPR-Cas-based assays depends more directly on the equilibrium concentrations of signal-initiating species; these cannot be determined directly from $K_{eq}$ for some systems. This is the motivation for use of the parameter $\alpha$, defined as the fraction of ANA in the annealed or blocked state in the initial equilibrium. This value enables simplifications that would otherwise be difficult to generalize across systems with different ANA designs. Like $K_{eq}$, $\Delta G^\circ$ is intrinsic for a specific reaction, such as the annealing of a particular DNA secondary structure. Therefore, it can be easily converted to $K_{eq}$ and serves a similar purpose. Using the $\Delta G^\circ_{anneal}$ value for a single secondary structure has important limitations, however, as large or complex nucleic acid structures exist in many states at equilibrium. The predicted $\Delta G^\circ$ value for the entire ensemble of states can be used instead, but this value does not correspond to a specific reaction between two or three states. For common types of ANA, this note details the relationship between $\alpha$, $K_{eq}$, and $\Delta G^\circ$.

**Hairpin ANA**
For a hairpin ANA, the definition of $\alpha$ is straightforward because in the equilibrium:

$$ANA_{melted} \rightleftharpoons ANA_{annealed}$$

both species are the same molecule, but in different states. Therefore, $\alpha$ is defined as:

$$\alpha = \frac{[ANA_{annealed}]_{eq}}{[ANA_{tot}]_{eq}} = \frac{[ANA_{annealed}]_{eq}}{[ANA_{annealed}]_{eq} + [ANA_{melted}]_{eq}}$$

when the system is at equilibrium. If we define $K_{eq}$ for this system:

$$K_{eq} = \frac{[ANA_{annealed}]_{eq}}{[ANA_{melted}]_{eq}}$$

we can observe the following relationship:

$$K_{eq} = \frac{\alpha}{1-\alpha}, \alpha = \frac{K_{eq}}{1+K_{eq}}$$

For large $K_{eq}$ ($\alpha \approx 1$), calculating $\alpha$ directly can lead to loss of precision. Instead, it is better to define:

$$\alpha_c = 1 - \alpha$$

and calculate:

$$\alpha_c = \frac{1}{1+K_{eq}}$$

as an intermediate value. Because $K_{eq}$ is intrinsic to the reaction and $\alpha$ can be stated entirely in terms of $K_{eq}$, $\alpha$ is also intrinsic for hairpin ANAs conforming to this equilibrium.

For each hairpin ANA, the value of $\Delta G^\circ_{anneal}$ for a certain temperature (the value typically reported by nucleic acid analysis software for a specific secondary structure) can be converted to $K_{eq}$ via the equation:

$$K_{eq} = e^{-\Delta G^\circ_{anneal}/RT} = e^{\Delta G^\circ_{melt}/RT}$$

For a nucleic acid that spontaneously anneals to a high degree (highly negative $\Delta G^\circ_{anneal}$), therefore, $K_{eq}$ will be very large, indicating an equilibrium dominated by the annealed component. To convert directly from $\Delta G_{anneal}$ to $\alpha$:

$$\alpha = \frac{e^{-\Delta G^\circ_{anneal}/RT}}{1+e^{-\Delta G^\circ_{anneal}/RT}} = \frac{1}{1+e^{\Delta G^\circ_{anneal}/RT}}$$

The value of $\alpha$ for this type of system can be calculated with the **calculateAlphaUnimolecular** function.

**Two-component ANA**
For a two-component ANA, the equilibrium can be defined as:

$$T + C \rightleftharpoons ANA$$

where $T$ is the target-containing strand and $C$ is the complementary or partially complementary blocking strand. In this case, $\alpha$ is defined as:

$$\alpha = \frac{[ANA]_{eq}}{[T]_0} = \frac{[ANA]_{eq}}{[ANA]_{eq} + [T]_{eq}}$$

This can be approached in two ways depending on the starting conditions for achieving equilibrium.

Case 1: Starting with target and complement

Assuming the system is initiated by adding only some concentration of target ($[T]_0$) and complement ($[C]_0$), $K_{eq}$ for this system is defined as:

$$K_{eq} = \frac{[ANA]_{eq}}{[T]_{eq}[C]_{eq}} = \frac{\alpha[T]_0}{([T]_0 - \alpha[T]_0)[C]_{eq}}$$

Thus, the relationship between the two parameters is:

$$K_{eq} = \frac{\alpha}{[C]_{eq}(1-\alpha)}, \alpha = \frac{K_{eq}}{1/[C]_{eq} + K_{eq}}$$

Based on the species conservation for $T$ and $C$:

$$[C]_0 = [C]_{eq} + [ANA]_{eq}$$

it can be stated:

$$[C]_{eq} = [C]_0 - [ANA]_{eq} = [C]_0 - \alpha[T]_0$$

resulting in:

$$K_{eq} = \frac{\alpha}{([C]_0 - \alpha[T]_0)(1-\alpha)}, \alpha = \frac{K_{eq}([C]_0 - \alpha[T]_0)}{1 + K_{eq}([C]_0 - \alpha[T]_0)}$$

If we express the complement in terms of the target concentration:

$$[C]_0 = \kappa[T]_0$$

then:

$$K_{eq} = \frac{\alpha}{[T]_0(\kappa-\alpha)(1-\alpha)}, \alpha = \frac{K_{eq}[T]_0(\kappa-\alpha)}{1 + K_{eq}[T]_0(\kappa-\alpha)}$$

While $K_{eq}$ is intrinsic to the reaction, $\alpha$ is extrinsic for two-component ANA designs because it depends on the initial concentrations used. While it is trivial to convert form $\alpha$ to $K_{eq}$, the reverse requires solving the quadratic equation:

$$\left(K_{eq}[T]_0\right)\alpha^2 - \left(1 + K_{eq}[T]_0(\kappa+1)\right)\alpha + K_{eq}\kappa[T]_0 = 0$$

which is solved by:

$$\alpha = \frac{1 + K_{eq}[T]_0(\kappa+1) - \sqrt{\left(1 + K_{eq}[T]_0(\kappa+1)\right)^2 - 4K_{eq}^2\kappa[T]_0^2}}{2K_{eq}[T]_0}$$

which results in a value between zero and one. Like before, this can result in a loss of precision if $\alpha$ is close to 1. Because of this, it is better to first calculate $\alpha_c$:

$$\alpha_c = \frac{1}{1 + K_{eq}[T]_0(\alpha_c + \kappa - 1))}$$

This results in the quadratic equation:

$$\left(K_{eq}[T]_0\right)\alpha_c^2 + \left(1 + K_{eq}[T]_0(\kappa-1)\right)\alpha_c - 1 = 0$$

which is solved by:

$$\alpha_c = \frac{-1 - K_{eq}[T]_0(\kappa-1) + \sqrt{\left(1 + K_{eq}[T]_0(\kappa-1)\right)^2 + 4K_{eq}[T]_0}}{2K_{eq}[T]_0}$$

In terms of $\Delta G^\circ_{anneal}$, the value is:

$$\alpha_c = \frac{-1 - [T]_0(\kappa-1)e^{-\Delta G^\circ_{anneal}/RT} + \sqrt{\left(1 + [T]_0(\kappa-1)e^{-\Delta G^\circ_{anneal}/RT}\right)^2 + 4[T]_0 e^{-\Delta G^\circ_{anneal}/RT}}}{2[T]_0 e^{-\Delta G^\circ_{anneal}/RT}}$$

Case 2: Starting with ANA

If we instead initiate the system by introducing a certain amount of ANA with no other components, as would occur when cleaving a hairpin ANA:

$$[ANA]_0 = [C]_{eq} + [ANA]_{eq}$$

In this case:

$$\alpha = \frac{[ANA]_{eq}}{[ANA]_0}$$

Therefore:

$$[C]_{eq} = [T]_{eq} = [ANA]_0 - [ANA]_{eq} = [ANA]_0 - \alpha[ANA]_0 = (1-\alpha)[ANA]_0$$

Substituting these in the definitions for $K_{eq}$:

$$K_{eq} = \frac{[ANA]_{eq}}{[T]_{eq}[C]_{eq}} = \frac{\alpha[ANA]_0}{(1-\alpha)^2[ANA]_0^2}$$

$$K_{eq} = \frac{\alpha}{(1-\alpha)^2[ANA]_0}, \alpha = \frac{K_{eq}(1-\alpha)[ANA]_0}{1 + K_{eq}(1-\alpha)[ANA]_0}$$

allows us to solve a quadratic for $\alpha$:

$$\left([ANA]_0 K_{eq}\right)\alpha^2 - \left(2[ANA]_0 K_{eq} + 1\right)\alpha + [ANA]_0 K_{eq} = 0$$

as:

$$\alpha = 1 + \frac{1 - \sqrt{4k+1}}{2k}$$

where:

$$k = [ANA]_0 K_{eq} = [ANA]_0 e^{-\Delta G^\circ_{anneal}/RT}$$

and:

$$\alpha_c = 1 - \alpha = \frac{\sqrt{4k+1} - 1}{2k}$$

The value of $\alpha$ for this type of system can be calculated with the **calculateAlphaBimolecular** function.

**n-component ANA**

For any ANA with a target strand and n complement components, the equilibrium is:

$$T + C_1 + \cdots + C_n \rightleftharpoons ANA$$

where $T$ is the target-containing strand and each $C_i$ is a distinct partially complementary blocking strand. This case is most relevant when a complement may be digested via trans-cleavage into multiple components that may remain bound to the target. In this case, $\alpha$ is defined as:

$$\alpha = \frac{[ANA]_{eq}}{[T]_0} = \frac{[ANA]_{eq}}{[ANA]_{eq} + [T]_{eq}}$$

assuming the system is initiated by adding only some concentration of target ($[T]_0$) and the respective complements. $K_{eq}$ for this system is defined as:

$$K_{eq} = \frac{[ANA]_{eq}}{[T]_{eq} \prod_{i=1}^{n} [C_i]_{eq}}$$

Thus, the relationship between the two parameters is:

$$K_{eq} = \frac{\alpha}{\prod_{i=1}^{n} [C_i]_{eq} (1-\alpha)}, \alpha = \frac{K_{eq}}{1/\prod_{i=1}^{n} [C_i]_{eq} + K_{eq}}$$

From the species conservation for $T$ and $C$:

$$[C_i]_0 = [C_i]_{eq} + [ANA]_{eq}$$

it can be stated:

$$[C_i]_{eq} = [C_i]_0 - [ANA]_{eq} = [C_i]_0 - \alpha[T]_0$$

If we express the complement in terms of the target concentration:

$$[C_i]_0 = \kappa_i [T]_0$$

then:

$$[C_i]_{eq} = [T]_0(\kappa_i - \alpha)$$

This results in:

$$K_{eq} = \frac{\alpha}{[T]_0^n (1-\alpha) \prod_{i=1}^{n} (\kappa_i - \alpha)}$$

$$\alpha = \frac{K_{eq}[T]_0^n \prod_{i=1}^{n} (\kappa_i - \alpha)}{1 + K_{eq}[T]_0^n \prod_{i=1}^{n} (\kappa_i - \alpha)}, \alpha_c = \frac{1}{1 + K_{eq}[T]_0^n \prod_{i=1}^{n} (\alpha_c + \kappa_i - 1)}$$

If we assume that each $\kappa_i$ is the same ($\kappa$), for example because each complement is released with a fixed 1:1 stoichiometry, then:

$$K_{eq} = \frac{\alpha}{[T]_0^n(\kappa-\alpha)^n(1-\alpha)}, \alpha = \frac{K_{eq}[T]_0^n(\kappa-\alpha)^n}{1+K_{eq}[T]_0^n(\kappa-\alpha)^n}, \alpha_c = \frac{1}{1+K_{eq}[T]_0^n(\alpha_c+\kappa-1)^n}$$

In either case, finding $\alpha$ and $\alpha_c$ involves solving higher-order polynomials and should be approached with numerical methods. The value of $\alpha$ for this type of system can be calculated with the **calculateAlphaNmolecular** function.

#### Supplementary Note 6: Gel Experimental Protocol

**Materials**

The Cas12 enzyme (Alt-R LbCas12a (Cpf1) Ultra) was obtained from Integrated DNA Technologies (IDT). All nucleic acid sequences were synthesized by IDT and resuspended in Tris-EDTA (TE) buffer as specified by IDT. 1M Tris-HCl Buffer (15567027) and NaCl (5M, RNase-free, AM9760G) were purchased from ThermoFisher Scientific. Magnesium Chloride ($MgCl_2$) Solution (B9021S) and NEBuffer r2.1 (B6002S) were purchased from New England Biolabs (NEB). Nuclease-free water (NFW, 11-05-01-04) and 10/60 Ladder (51-05-15-01) were purchased from IDT. 10X TBE Buffer (1610733), ammonium persulfate (APS, 1610700), and TEMED (1610800) were purchased from Bio-Rad. Gel Loading Dye Purple (6X, B7024S) was purchased from NEB. SYBR Green I Nucleic Acid Gel Stain (S7563) was purchased from ThermoFisher Scientific.

**Sequences**

| Strand Name | Sequence (5'→3', Target Region Underlined) | Length (nt) |
|---|---|---|
| gRNA | UAAUUUCUACUAAGUGUAGAUGGAUA<br>UACGAUAUAUAUAUAU | 42 |
| Target | AGGGGCGTGCATATATATATATCGTAT<br>ATCCGTTGGTGACG | 41 |

**CRISPR-Cas Reaction**

The CRISPR-Cas enzyme was activated by preparing the mix below and incubation for 30 min at RT:

| Active RNPT1 Mix | | | |
|---|---|---|---|
| **Reagent** | **[Stock] (M)** | **[Desired] (M)** | **Volume (µL)** |
| Cas12 | 5.00E-07 | 1.00E-07 | 24.00 |
| gRNA | 5.00E-07 | 1.00E-07 | 24.00 |
| Target | 1.00E-06 | 1.00E-07 | 12.00 |
| MgCl2 | 2.50E-02 | 5.00E-03 | 24.00 |
| NFW | | | 36.00 |

The ANA for a given experiment was prepared according to:

| NA Mix (Two-component ANA) | | | |
|---|---|---|---|
| Complement-to-Target Ratio | | 1.2 | |
| **Reagent** | **[Stock] (M)** | **[Desired] (M)** | **Volume (µL)** |
| Target | 1.00E-04 | 5.00E-06 | 2.50 |
| Comp | 1.00E-04 | 6.00E-06 | 3.00 |
| Tris HCl | 1 | 2.00E-02 | 1.00 |
| NaCl | 5 | 5.00E-02 | 0.50 |
| MgCl2 | 2.50E-02 | 1.00E-02 | 20.00 |
| NFW | | | 23.0 |

The ANA mix intended for direct visualization in the gel (without annealing) was reserved and diluted to match the final concentration of the reaction groups (4 µL NA mix, 2 µL $MgCl_2$, and 14 µL NFW). The rest of the NA mix was annealed at 95°C for 4 min and then returned to room temperature at a rate of 0.1°C/s. ANA mix was similarly reserved after annealing and diluted to match the final concentration of the reaction groups.

Reaction mix was prepared with the following components (temporarily omitting the active RNPT1 mix):

| Reaction Mix | | | |
|---|---|---|---|
| Reagent | [Stock] (M/X) | [Desired] (M) | Volume (µL) |
| NEB Buffer | 1.00E+01 | 1.00E+00 | 18.00 |
| MgCl2 | 2.50E-02 | 2.50E-03 | 18.00 |
| Active RNPT1 Mix | 1.00E-07 | 5.00E-08 | 90.00 |
| NA Mix | 5.00E-06 | 1.00E-06 | 36.00 |
| NFW | | | 18.00 |

The mixture was split into four tubes, each containing 16 µL. 4 µL of NA mix was added to the first tube and it was placed in a heat block at 37°C. The remaining tubes and NA mix were held at 4°C. At each subsequent time point (1 and 2 hours later), NA mix was added to another tube and the tube was placed in the heat block. After three hours, NA mix was added to the final tube and all tubes were placed on ice to halt the reaction.

**Gel Electrophoresis**
High-resolution non-denaturing polyacrylamide gel electrophoresis (PAGE) experiments were prepared with 15% polyacrylamide in 10xTBE (89 mM Tris, 89 mM boric acid, 2 mM EDTA). Gel solution was prepared by mixing 3.75 mL Biorad 40% Acrylamide/Bis Solution, 5.25 mL degassed water, 1 mL 10X TBE 50 µL APS (10% w/v solution), and 5 µL TEMED. 10/60 Ladder from IDT was prepared according to the instructions by resuspending the lyophilized product in 100 µL TE buffer. 4 µL of the prepared 10/60 ladder was mixed with 6 µL water and 1 µL gel loading dye. After quenching the reactions, 2 µL of loading dye was added (with a ratio of 8 µl of ANA sample solution to 2 µl of loading dye) and the resulting solution was immediately loaded into the wells of the 15% acrylamide gel. Electrophoresis was carried out at room temperature under a constant voltage of 120 V for 1 hour and 40 minutes in the Mini-Protean Tetra Vertical Electrophoresis Cell with the PowerPac Basic Power Supply (Bio-Rad). Following electrophoresis, gels were stained for visualization by submerging in a solution prepared at a ratio of 5 µL 10x SYBR gel stain to 50 µL 1x TBE buffer and gently covered and agitated at 150 rpm for 40 min. Gels were imaged using a Bio-Rad Gel Doc XR+ gel documentation system.

**Complete Gel Results**
The complete gel electrophoresis result is included below, with the cropped image from the main text figure outlined. The lanes are: (1) empty, (2) ladder intended for higher molecular weight bands, (3) pre-annealing nucleic acid, (4) post-annealing nucleic acid, nucleic acid after (5) one hour, (6) two hours, (7) and three hours of cleavage, (8) 10/60 ssDNA ladder, (9) empty, and (10) empty.

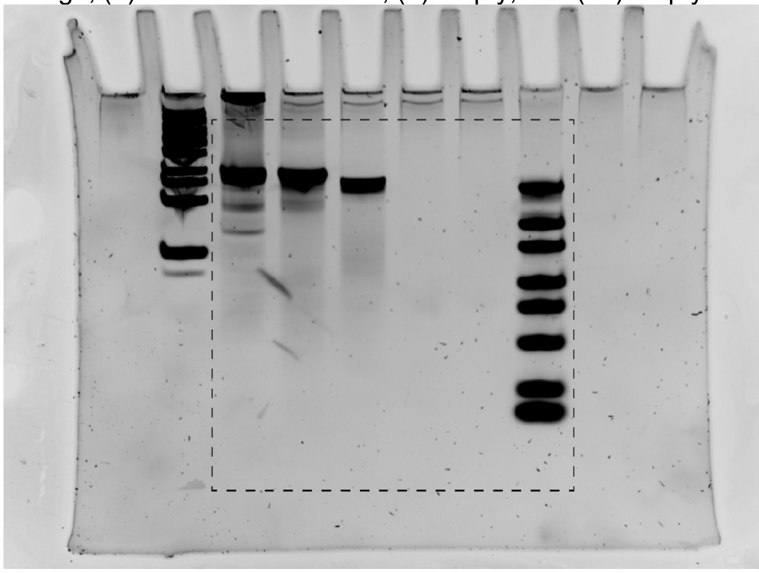

#### Supplementary Note 7: DNA Fragmentation Kinetics

**Basic Model**
Background ssDNA is subject to degradation by trans-cleavage. Because this occupies activated enzyme, it can present significant inhibition of reporter cleavage. The reaction for background cleavage of DNA fragments can be written as:

$$E + F_l \rightleftharpoons EF_l \rightarrow E + F_m + F_n$$

where a fragment of length $l$ is cleaved into two smaller fragments of length $m$ and $n$ and $m + n = l$. A fragment of the maximum length considered for the reaction ($L_{tot}$) can be cleaved until it is below some threshold length, $L_{th}$, below which fragments are no longer cleaved. We can consider the fragment concentrations for fragments between these limits as vectors:

$$E + \boldsymbol{F} \rightleftharpoons \boldsymbol{EF} \rightarrow E + \boldsymbol{F}$$

where:

$$\boldsymbol{F} = \begin{matrix} [F_{L_{tot}}] \\ [F_{L_{tot}-1}] \\ \vdots \\ [F_{L_{th}}] \end{matrix} \text{ and } \boldsymbol{EF} = \begin{matrix} [EF_{L_{tot}}] \\ [EF_{L_{tot}-1}] \\ \vdots \\ [EF_{L_{th}}] \end{matrix}$$

**Affinity Vector**
There are two general assumptions we could make about the affinity of the Cas enzyme for different fragments. For a site-specific interaction such as cis-cleavage, binding could depend on the number of sites in the fragment; as a result the fraction of enzyme bound would scale with fragment length. For non-specific interactions like Cas12a trans-cleavage, it is more reasonable to assume that binding is limited by encounters between the enzyme and DNA. Thus, binding is independent of length. To model either of these, we can modify the reverse rate constants in the reaction to reflect the tendency for the enzyme to stay bound to the fragment. The simpler case is length-independent binding. An affinity vector of size $(L_{tot} - L_{th} + 1) \times 1$ can be constructed:

$$\boldsymbol{A} = \begin{bmatrix} 1 \\ \vdots \\ 1 \\ 0 \end{bmatrix}$$

This can be thought of as maintaining the same rate constant $k_r$ for each of the fragments above length $L_{th}$. The final element is zero because fragments of length $L_{th}$ are considered not to bind.
For length-dependent binding, a different affinity vector can be generated. This is multiplied by $(L_{tot} - L_{th})/\Sigma\boldsymbol{L}$ so that $\Sigma\boldsymbol{A}$ is kept constant between the length-independent and length-dependent case. This keeps the effective $k_r$ for the entire reaction system the same as in the length-independent case:

$$\boldsymbol{L} = \begin{bmatrix} L_{tot} \\ L_{tot} - 1 \\ \vdots \\ L_{th} + 1 \\ 0 \end{bmatrix}, \boldsymbol{A} = \frac{(L_{tot} - L_{th})}{\Sigma\boldsymbol{L}}\boldsymbol{L}$$

The type of affinity vector can be selected in the code by changing the affinity_type constant between 'flat' (length-independent) and 'length' (length-dependent).

**Fragmentation Matrix**
To complete the description of this process, it is necessary to determine what fragment lengths will result from cleavage of a fragment with a certain length. Like before, there are two cases that can be considered. The first is that the cleavage occurs at a random location and the second is that the cleavage occurs near the end of the strand. For the more basic case, we can consider cleavage at a random location. It can be observed that for a DNA sequence of length $l$, there are $l - 1$ potential cleavage locations, resulting in fragments of lengths $(1, l - 1), (2, l - 2), \dots, (l - 1, 1)$ with each length from $1$ to $l$ appearing twice. Therefore, the probability of a certain cut length resulting from a cleavage event is:

$$P_n = 2/(l - 1)$$

with:

$$\sum_{n=1}^{l-1} P_n = 2$$

because each cleavage event doubles the number of fragments. All lengths below $L_{th}$ are treated as identical because their kinetic role in the reaction is the same. For each length from $L_{tot}$ to $L_{th}$, a fragmentation matrix ($\boldsymbol{M_F}$) of size $(L_{tot} - L_{th} + 1) \times (L_{tot} - L_{th} + 1)$ can be constructed with entries containing the probability that a fragment of length $L_{tot} - i + 1$ will result from splitting a fragment with length $L_{tot} - j + 1$:

$$\boldsymbol{M}_{i,j} = \begin{cases} 0 & \text{for} \quad i \leq j, i \neq L_{tot} - L_{th} + 1 \\ 1 & \text{for} \quad i = j = L_{tot} - L_{th} + 1 \\ 2/(L_{tot} - j) & \text{for} \quad i > j, i \neq L_{tot} - L_{th} + 1 \\ 2L_{th}/(L_{tot} - j) & \text{for} \quad i = L_{tot} - L_{th} + 1 \end{cases}$$

Because fragments must decrease in size from cleavage, this will be a lower triangular matrix. For a fragment of length five ($L_{tot} = 5$) with a threshold of two ($L_{th} = 2$), the fragmentation matrix would be:

$$\boldsymbol{M_F} = \begin{bmatrix} 0 & 0 & 0 & 0 \\ 1/2 & 0 & 0 & 0 \\ 1/2 & 2/3 & 0 & 0 \\ 1 & 4/3 & 2 & 1 \end{bmatrix}$$

Multiplying the fragmentation matrix $\boldsymbol{M_F}$ by the enzyme-substrate concentration matrix $\boldsymbol{EF}$ and $k_{cat}$ gives the rate of production of each size fragment by catalysis, ignoring binding/unbinding:

$$\boldsymbol{v_{cat}} = k_{cat} \boldsymbol{M_F EF}$$

Practically, this means that a fragment of length 5 (column 4) would result in cleavage products that are 50% length 2 or below, 25% length 3, and 25% length 4. All columns of $\boldsymbol{M_F}$ other than the last will sum to two because an extra fragment is produced in each cleavage event. The last column is insignificant because, as described in the previous section, none of the fragments at or below the length threshold should be bound to the enzyme.

It is likely that the trans-cleavage site is not entirely random but rather may proceed from the 3' end to a random cleavage location.[6,24] In this case, we can construct a fragmentation matrix that indicates cleavage near one end. We can define a region from the end where the cleavage is equally likely to occur:

$$r \in (n_1, n_2)$$

Each column of the matrix must be calculated individually in this case. An algorithm for this calculation is implemented in the **buildFragmentationMatrix** function. As an example, we can consider cleavage of a fragment of length 10, with a region between bases 2 and 5. In this case, the enzyme will cleave at bonds 2-3, 3-4, or 4-5, resulting in fragments of length 2 and 8, 3 and 7, or 4 and 6, respectively. Thus, the probability of a fragment of length 2, 3, 4, 6, 7, and 8 are each 1/6, while the probability of a fragment with length 5 is zero. However, because two fragments are produced in each cleavage event, each probability is doubled so that the sum of the fragments produced is two. For inputs of $L_{tot} = 10$, $L_{th} = 2$, and $r = (2,5)$, the resulting matrix is:

| | | | | | | | | |
|---|---|---|---|---|---|---|---|---|
| 0 | 0 | 0 | 0 | 0 | 0 | 0 | 0 | 0 |
| 0 | 0 | 0 | 0 | 0 | 0 | 0 | 0 | 0 |
| 1/3 | 0 | 0 | 0 | 0 | 0 | 0 | 0 | 0 |
| 1/3 | 1/3 | 0 | 0 | 0 | 0 | 0 | 0 | 0 |
| 1/3 | 1/3 | 1/3 | 0 | 0 | 0 | 0 | 0 | 0 |
| 0 | 1/3 | 1/3 | 1/3 | 0 | 0 | 0 | 0 | 0 |
| 1/3 | 1/3 | 2/3 | 2/3 | 2/3 | 1/3 | 0 | 0 | 0 |
| 1/3 | 1/3 | 1/3 | 2/3 | 2/3 | 2/3 | 0.5 | 0 | 0 |
| 1/3 | 1/3 | 1/3 | 1/3 | 2/3 | 1 | 1.5 | 2 | 1 |

The type of fragmentation matrix can be selected in the code by changing the fragmentation_range constant; an empty vector ([]) results in random cleavage while a vector with a range (e.g. [2,5]) results in region-specific cleavage.

**Implementation in Differential Equations**

With this, we can write the complete rate equations for the fragmentation system:

$$\frac{dE}{dt} = -k_f[E]\sum \boldsymbol{F} + \sum\left(\frac{k_r}{\boldsymbol{A}+\epsilon\boldsymbol{1}} + k_{cat}\boldsymbol{1}\right)\cdot \boldsymbol{EF}$$

$$\frac{d\boldsymbol{F}}{dt} = -k_f[E]\boldsymbol{F} + \left(\frac{k_r}{\boldsymbol{A}+\epsilon\boldsymbol{1}}\right)\cdot \boldsymbol{EF} + k_{cat}\boldsymbol{M_F}\boldsymbol{EF}$$

$$\frac{d\boldsymbol{EF}}{dt} = k_f[E]\boldsymbol{F} - \left(\frac{k_r}{\boldsymbol{A}+\epsilon\boldsymbol{1}} + k_{cat}\boldsymbol{1}\right)\cdot \boldsymbol{EF}$$

where multiplication by $\boldsymbol{1}$ indicates broadcasting the scalar quantity onto a length-matched column vector. The affinity vector is implemented as affinity-based unbinding by applying it to the reverse rate. This results in numerical breakdown when the affinity vector contains a zero. To avoid this, the steps that involve dividing by $\boldsymbol{A}$ instead divide by $\boldsymbol{A}+\epsilon$, which is trivial because MATLAB includes a built-in epsilon function. Note that the production of $\boldsymbol{F}$ via catalysis ($+k_{cat}\boldsymbol{M_F}\boldsymbol{EF}$) is not balanced by the consumption of $\boldsymbol{EF}$ in the last equation ($-k_{cat}\boldsymbol{EF}$). This is reasonable because each cleavage event will increase the total concentration of fragments. Specifically, the increase in concentration is:

$$\frac{d\boldsymbol{F}_{tot}}{dt} = \frac{d\boldsymbol{F}}{dt} + \frac{d\boldsymbol{EF}}{dt} = k_{cat}\boldsymbol{M_F}\boldsymbol{EF} - k_{cat}\boldsymbol{EF} = k_{cat}(\boldsymbol{M_F} - \boldsymbol{I})\boldsymbol{EF}$$

The total scalar change in concentration of fragments is:

$$\frac{dF_{tot}}{dt} = k_{cat}\sum_{i=1}^{L_{tot}-L_{th}} \boldsymbol{EF}$$

Because the last element of $\boldsymbol{EF}$ (the only one excluded from this sum) will be zero because of the zero binding probability, this is equivalent to:

$$\frac{dF_{tot}}{dt} = k_{cat}\sum \boldsymbol{EF}$$

In addition to changing the affinity vector and fragmentation matrix, the DNA length boundaries can be set by modifying the constant DNA_bounds. By default, for example, DNA_bounds is set to [49 2] to consider fragments as large as 49 nucleotides and consider fragments of length 2 or less completely degraded. Setting DNA_bounds to an empty vector turns off the DNA fragmentation mechanism entirely.

#### Supplementary Note 8: Exponential Phase Analytical Simplification

Using the principles typically applied to Michaelis-Menten systems, we can derive an analytical simplification for the cleaved reporter concentration during the exponential phase of the reaction. This analytical simplification is potentially valid only during the early exponential phase of the reaction because of the assumptions that consumption of inactivated E2, ANA, and reporter are negligible.

To start, we can assume that cis-cleavage occurs at a much higher rate than trans-cleavage. We will also assume that the cleaved ANA converts into the second enzyme target (T2) at a much higher rate than other phenomenon, and inactivation of the enzyme after cis-cleavage is negligible. A consequence of these assumptions is that any T1 and cleaved ANA immediately and irreversibly activates the respective RNP complex. We will also ignore other effects reflected in the complete system, like the interaction with background DNA. This simplifies the system to four reactions:

If the active RNP complex specific to the assay target ($[E1_{act}]$) cleaves the ANA, it immediately converts the ANA to T2, which results in an active secondary RNP ($[E2_{act}]$) while maintaining the first:

$$E1_{act} + ANA \rightleftharpoons E1_{act}ANA \rightarrow E1_{act} + E2_{act} \qquad (a)$$

Similarly, $[E2_{act}]$ can cleave the ANA and produce a second active $[E2_{act}]$.

$$E2_{act} + ANA \rightleftharpoons E2_{act}ANA \rightarrow 2E2_{act} \qquad (b)$$

Both complexes may also trans-cleave the reporter (FQ):

$$E1_{act} + FQ \rightleftharpoons E1_{act}FQ \rightarrow E1_{act} + F + Q \qquad (c)$$

$$E2_{act} + FQ \rightleftharpoons E2_{act}FQ \rightarrow E2_{act} + F + Q \qquad (d)$$

The complexity of solving this system is mostly due to the autocatalytic reaction (b), from which exponential behavior arises. We will assume that $k_f$, $k_r$, and $k_{cat}$ are the same for each trans-cleavage reaction.

The main equation to be solved is:

$$\frac{d[F]}{dt} = k_{cat}([E1_{act}FQ] + [E2_{act}FQ])$$

We can apply the quasi-steady-state assumption to the enzyme-substrate complexes:

$$\frac{d[E1_{act}ANA]}{dt} = \frac{d[E1_{act}FQ]}{dt} = \frac{d[E2_{act}ANA]}{dt} = \frac{d[E2_{act}FQ]}{dt} \approx 0$$

This allows us to simplify the relationships between the reaction species to obtain equations for the enzyme-substrate complexes, for example:

$$-(k_{cat} + k_r)[E1_{act}ANA] + k_f[E1_{act}][ANA] = 0$$

results in:

$$[E1_{act}ANA] = \frac{k_f[E1_{act}][ANA]}{k_{cat} + k_r} = \frac{[E1_{act}][ANA]}{K_M}$$

Similarly:

$$[E1_{act}FQ] = \frac{[E1_{act}][FQ]}{K_M}$$

$$[E2_{act}ANA] = \frac{[E2_{act}][ANA]}{K_M}$$

$$[E2_{act}FQ] = \frac{[E2_{act}][FQ]}{K_M}$$

Species conservation requires:

$$[E1_{act}]_{tot} = [E1_{act}] + [E1_{act}ANA] + [E1_{act}FQ]$$

$$[E2_{act}]_{tot} = [E2_{act}] + [E2_{act}ANA] + [E2_{act}FQ]$$

so:

$$[E1_{act}]_{tot} = [E1_{act}] + \frac{[E1_{act}][ANA]}{K_M} + \frac{[E1_{act}][FQ]}{K_M} = \frac{[E1_{act}](K_M + [ANA] + [FQ])}{K_M}$$

and:

$$[E1_{act}] = \frac{[E1_{act}]_{tot}K_M}{K_M + [ANA] + [FQ]}$$

By the same process:

$$[E2_{act}] = \frac{[E2_{act}]_{tot}K_M}{K_M + [ANA] + [FQ]}$$

Consequently, the concentrations of the other species are:

$$[E1_{act}ANA] = \frac{[E1_{act}]_{tot}K_M}{K_M + [ANA] + [FQ]}\frac{[ANA]}{K_M} = \frac{[E1_{act}]_{tot}[ANA]}{K_M + [ANA] + [FQ]}$$

$$[E1_{act}FQ] = \frac{[E1_{act}]_{tot}[FQ]}{K_M + [ANA] + [FQ]}$$

$$[E2_{act}ANA] = \frac{[E2_{act}]_{tot}[ANA]}{K_M + [ANA] + [FQ]}$$

$$[E2_{act}FQ] = \frac{[E2_{act}]_{tot}[FQ]}{K_M + [ANA] + [FQ]}$$

The rate of fluorophore generation is then:

$$\frac{d[F]}{dt} = k_{cat}([E1_{act}FQ] + [E2_{act}FQ]) = \frac{k_{cat}[FQ]}{K_M + [ANA] + [FQ]}([E1_{act}]_{tot} + [E2_{act}]_{tot})$$

While $[E1_{act}]_{tot}$ is constant, $[E2_{act}]_{tot}$ will increase through the exponential phase of the autocatalytic reaction. It is thus denoted as $[E2_{act}]_{tot}(t)$ and increases by the catalysis of (a) and (b) according to:

$$\frac{d[E2_{act}]_{tot}}{dt} = k_{cat}([E1_{act}ANA] + [E2_{act}ANA]) = \frac{k_{cat}[ANA]}{K_M + [ANA] + [FQ]}([E1_{act}]_{tot} + [E2_{act}]_{tot})$$

To reach a closed-form solution, we will assume that ANA and reporter are abundant enough that depletion is minimal at early time points, i.e.:

$$\frac{d[ANA]}{dt} = \frac{d[FQ]}{dt} \approx 0$$

We can then define constants:

$$C_1 = \frac{k_{cat}[ANA]}{K_M + [ANA] + [FQ]} \text{ and } C_2 = \frac{k_{cat}[FQ]}{K_M + [ANA] + [FQ]}$$

With:

$$\frac{d[E2_{act}]_{tot}}{dt} = C_1\left([E1_{act}]_{tot} + [E2_{act}]_{tot}(t)\right)$$

We will assume some constant amount of ANA is available at the start of the reaction due to its equilibrium dynamics. Due to the immediate cis-cleavage, this results in:

$$[E2_{act}]_{tot}(0) = [E2_{act}]_0$$

Then:

$$[E2_{act}]_{tot}(t) = ([E2_{act}]_0 + [E1_{act}]_{tot})e^{C_1 t} - [E1_{act}]_{tot}$$

and:

$$\frac{d[F]}{dt} = C_2\left([E1_{act}]_{tot} + [E2_{act}]_{tot}(t)\right) = C_2([E2_{act}]_0 + [E1_{act}]_{tot})e^{C_1 t}$$

Assuming the concentration of cleaved reporter is initially zero ($[F](0) = 0$):

$$\int_0^{F(t)} d[F] = \int_0^t C_2([E2_{act}]_0 + [E1_{act}]_{tot})e^{C_1 t}\,dt$$

$$[F](t) = C_2([E2_{act}]_0 + [E1_{act}]_{tot})\frac{e^{C_1 t} - 1}{C_1} = \frac{C_2}{C_1}([E2_{act}]_0 + [E1_{act}]_{tot})(e^{C_1 t} - 1)$$

$$= \frac{[FQ]}{[ANA]}([E2_{act}]_0 + [E1_{act}]_{tot})\left(e^{t/\tau} - 1\right)$$

where the time constant $\tau$ is:

$$\tau = \frac{1}{C_1} = \frac{K_M + [ANA] + [FQ]}{k_{cat}[ANA]}$$

As described elsewhere, we consider the initial concentration of available ANA due to its equilibrium to be $1 - \alpha$, where $\alpha$ is the fraction of ANA that is inaccessible at equilibrium. This allows for simplification to:

$$[F](t) = \frac{[FQ]}{[ANA]}\left((1-\alpha)[ANA] + [E1_{act}]_{tot}\right)\left(e^{t/\tau} - 1\right) = [FQ]\left((1-\alpha) + \frac{[E1_{act}]_{tot}}{[ANA]}\right)\left(e^{t/\tau} - 1\right)$$

Because $[E1_{act}]_{tot} \approx [T1]$ (assuming the concentration of target is less than the first enzyme, which is typical in diagnostics):

$$[F](t) = \frac{[FQ]}{[ANA]}\left((1-\alpha)[ANA] + [T1]\right)\left(e^{t/\tau} - 1\right)$$

Similarly:

$$[E2_{act}]_{tot}(t) = \big((1-\alpha)[ANA] + [T1]\big)e^{t/\tau} - [T1]$$

The reaction will only be in the exponential phase while the amount of $[E_{act}]_{tot}$ does not exceed the amount of corresponding RNP available in the reaction $[E]_{max}$. Calculation of this quantity depends on the reaction. Assuming that enzymes are pre-complexed with the gRNA results and the dissociation is very slow, there is a maximum amount of inactive enzyme ($[E1]_{tot}$ and $[E2]_{tot}$), which can be approximated as the minimum of the enzyme and gRNA concentration. The maximum concentration of active enzyme is then:

$$[E1_{act}]_{max} = \min([E1]_{tot}, [T1])$$
$$[E2_{act}]_{max} = \min([E2]_{tot}, [ANA])$$

with:

$$[E]_{max} = [E1_{act}]_{max} + [E2_{act}]_{max}$$

This is dominated by $[E2_{act}]_{max}$ for most assay conditions. Based on this, we can estimate the duration of the exponential phase. The reaction should remain exponential while:

$$[E1_{act}]_{tot} + [E2_{act}]_{tot}(t) \leq [E]_{max}$$
$$[E1_{act}]_{tot} + [E_2]_{tot}(t) = [T1] + \big((1-\alpha)[ANA] + [T1]\big)e^{t/\tau} - [T1] \leq [E]_{max}$$

Or:

$$t_{exp} \leq \tau \ln\left(\frac{[E]_{max}}{(1-\alpha)[ANA] + [T1]}\right)$$

After complete conversion of the enzyme, the fluorescence is:

$$[F] = \frac{[FQ]}{[ANA]}\big((1-\alpha)[ANA] + [T1]\big)\left(\frac{[E]_{max}}{(1-\alpha)[ANA] + [T1]} - 1\right)$$
$$= \frac{[FQ]([E]_{max} - (1-\alpha)[ANA] - [T1])}{[ANA]}$$

Therefore, the fraction of reporter cleaved during the exponential phase is:

$$f_{cl,exp} = \frac{[F]}{[FQ]} = \frac{[E]_{max} - (1-\alpha)[ANA] - [T1]}{[ANA]}$$

For most assay conditions, the numerator is dominated by $[E]_{max}$. This allows us to simplify this value to:

$$f_{cl,exp} \approx \frac{[E]_{max}}{[ANA]}$$

#### Supplementary Note 9: Linear Phase Analytical Simplification

After the initial exponential phase, autocatalytic CRISPR reactions can be seen to proceed with a linear rate. This is consistent with the expectation that the enzyme will reach maximum activation either due to depletion of the enzyme or ANA.

Starting from the same reactions as Supplementary Note 9:

$$E1_{act} + ANA \rightleftharpoons E1_{act}ANA \rightarrow E1_{act} + E2_{act} \quad (a)$$

$$E2_{act} + ANA \rightleftharpoons E2_{act}ANA \rightarrow 2E2_{act} \quad (b)$$

$$E1_{act} + FQ \rightleftharpoons E1_{act}FQ \rightarrow E1_{act} + F + Q \quad (c)$$

$$E2_{act} + FQ \rightleftharpoons E2_{act}FQ \rightarrow E2_{act} + F + Q \quad (d)$$

We can assume that after the initial exponential phase, the amount of $E1_{act}$ is equal to the target concentration ($[T1]$). We will assume that the maximum possible concentration of enzyme is activated. As discussed in Supplementary Note 8, this is approximated as:

$$[E]_{max} = \min([E1]_{tot}, [T1]) + \min([E2]_{tot}, [ANA])$$

No further activation of the enzyme can occur, so reactions (a) and (b) do not produce additional enzyme concentration. In the case where $[ANA] > [E2]_{tot}$, however, the enzyme is still occupied by cleaving remaining ANA. We can rewrite these generally as:

$$E_{act} + ANA \rightleftharpoons E_{act}ANA \rightarrow E_{act}$$

We can also assume that the catalytic efficiency with which each activated enzyme cleaves reporter is similar. Therefore, we can combine reactions (c) and (d) to:

$$E_{act} + FQ \rightleftharpoons E_{act}FQ \rightarrow E_{act} + F + Q$$

The system then nearly follows Michaelis-Menten kinetics, but with the ANA acting as a competitive substrate. Because the inhibitor cleavage may have different kinetics than the reporter cleavage, we can consider a distinct $k_{1cat}$, $K_{1M}$ for the ANA and $k_{2cat}$, $K_{2M}$ for the reporter. Briefly, by assuming quasi-steady-state behavior, we can state:

$$[E_{act}FQ] = \frac{[E_{act}][FQ]}{K_{2M}} \text{ and } [E_{act}ANA] = \frac{[E_{act}][ANA]}{K_{1M}}$$

Species conservation requires:

$$[E_{act}]_{tot} = [E_{act}] + [E_{act}ANA] + [E_{act}FQ]$$

Therefore:

$$[E_{act}] = [E_{act}]_{tot} - \frac{[E_{act}][ANA]}{K_{1M}} - \frac{[E_{act}][FQ]}{K_{2M}}$$

$$[E_{act}] = [E_{act}]_{tot} / \left(1 + \frac{[ANA]}{K_{1M}} + \frac{[FQ]}{K_{2M}}\right)$$

And:

$$[E_{act}FQ] = \frac{[E_{act}]_{tot}[FQ]}{K_{2M}\left(1 + \frac{[ANA]}{K_{1M}} + \frac{[FQ]}{K_{2M}}\right)} =$$

Resulting in:

$$\frac{d[F]}{dt} = k_{2cat}[E_{act}FQ] = \frac{k_{2cat}[E_{act}]_{tot}[FQ]}{K_{2M} + [FQ] + \frac{K_{2M}}{K_{1M}}[ANA]}$$

This can alternatively be stated as:

$$\frac{d[F]}{dt} = \frac{k_{cat}[E_{act}]_{tot}[FQ]}{\kappa K_M + [FQ]}$$

with:

$$\kappa = \left(1 + \frac{[ANA]}{K_I}\right)$$

where $k_{cat}$ and $K_M$ are the rates associated with the intended substrate and $K_I$ is the rate associated with the competitive substrate. To account for the fact that some assays have excess ANA and others do not, we can use a parameter:

$$[ANA]_{excess} = \max([ANA] - [E2]_{tot}, 0)$$

This results in a positive value if $[ANA] > [E2]_{tot}$ (excess ANA available) and a zero value if $[ANA] \leq [E2]_{tot}$. We can restate:

$$\frac{d[F]}{dt} = \frac{k_{cat}[E_{act}]_{tot}[FQ]}{\kappa K_M + [FQ]}$$

with:

$$\kappa = \left(1 + \frac{[ANA]_{excess}}{K_I}\right)$$

This result is similar to the standard Michaelis-Menten result:

$$\frac{d[F]}{dt} = \frac{k_{cat}[E_{act}]_{tot}[FQ]}{K_M + [FQ]}$$

but with the inhibitor modifying the effective $K_M$ of the intended substrate. If the competitive substrate is cleaved with the same kinetics as the main substrate ($K_M = K_I$, as we assume in our model), this simplifies to:

$$\frac{d[F]}{dt} = \frac{k_{cat}[E_{act}]_{tot}[FQ]}{K_M + [ANA]_{excess} + [FQ]}$$

This clearly demonstrates how excess ANA can reduce the effective rate.

#### Supplementary Note 10: Evaluation of Analytical Predictions

In our previous analytical simplifications, we predicted the exponential phase would occur during the time:

$$t \leq \tau \ln\left(\frac{[E]_{max}}{(1-\alpha)[ANA]+[T_1]}\right)$$

where:

$$\tau = \frac{K_M + [ANA] + [FQ]}{k_{cat}[ANA]}$$

It was predicted to reach the following fraction of reporter cleaved during the exponential phase:

$$f_{cl,exp} = \frac{[E]_{max}}{[ANA]}$$

During the linear phase, it was predicted to cleave reporter at the rate:

$$\frac{d[F]}{dt} = \frac{k_{cat}[E_{act}]_{tot}[FQ]}{K_M + [ANA]_{excess} + [FQ]}$$

To understand whether the analytical model can be helpful in analyzing and optimizing these reactions, we can test these predictions against the full model. This is achieved by running simulated experiments across a variety of conditions and comparing the predicted values to the values measured from the simulated data.

To measure these parameters, we use a simple approach. First, the derivative of the cleaved reporter concentration is calculated. For a roughly sigmoidal curve, the maximum of this derivative approximates the inflection point. The inflection point is where the reporter cleavage rate reaches its maximum value, which generally occurs when the last of the Cas enzyme is activated. This is demonstrated in Figure SN10-1.

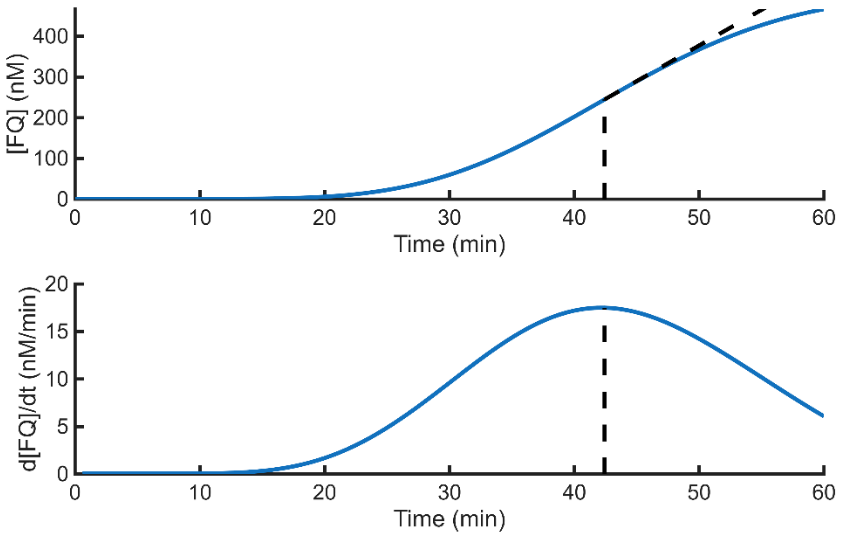


Figure SN10-1. Demonstration of the inflection point approximation.

Because the $[ANA]$, $[FQ]$, and $[E]_{max}$ are the primary modifiable parameters appearing in the equations, we will modify these.

<u>Experiment 1:</u>

First, we hold $[E2]$ constant and vary $[ANA]$ and $[FQ]$ for a typical hairpin ANA (Figure SN10-2).

a-b) The general trend for the predicted exponential time (a) and observed inflection point (b) are similar. Both show that increasing the concentration of ANA reduces the exponential phase time. This is a consequence of the ANA (both cleaved and uncleaved) driving signal amplification, which outweighs any inhibitory effect from excess ANA. The simulated data shows that increasing the reporter concentration slightly increases the duration of the exponential phase; this is primarily due to excess reporter inhibiting ANA cleavage. However, the magnitude of the effect is not fully reflected in the analytical simplification. Notably, the analytical simplification considerably underestimates the observed exponential time.

c-d) The analytical simplification predicts that increasing ANA concentration decreases the fraction cleaved (c). This does not hold in the simulated data (d).
d-e) The analytical model predicts that increasing the reporter concentration and decreasing the ANA concentration will both increase the slope in the linear phase (e). The simulated data shows this trend for the reporter but not for the ANA (f). This is likely because in the analytical model, cleaved ANA is assumed to immediately dissociate. In the simulation, the ANA dissociates much more slowly, resulting in a decreased cleavage rate at low ANA concentrations. The analytical simplification consistently overestimates the slope.

Experiment 2:

We can repeat the results from the first experiment but for an idealized ANA, which is approximated as $\Delta G^{\circ}_{uncleaved}$ = -30 and $\Delta G^{\circ}_{cleaved}$ = 100 (Figure SN10-3). This matches the simulation more closely to the ANA behavior assumed in the analytical model. With this modification, the trends from each of the analytical models match the simulated data almost exactly. This demonstrates that the key discrepancy between the analytical model and simulated data is the assumption about ANA behavior. The observed exponential phase duration is still substantially longer and the linear cleavage rate is lower than in the analytical predictions. This is likely because some reactions are assumed to be instantaneous in the analytical model while some sources of inhibition are ignored.

Experiment 3:

Next, we can hold $[FQ]$ constant and vary $[E2]$ and $[ANA]$ for a typical ANA (Figure SN10-4).
a-b) Like in experiment 1, the analytical model correctly predicts that increasing the ANA concentration increases the speed of the exponential phase. However, it also predicts that decreasing the enzyme concentration will slightly speed up the exponential phase because there is less enzyme to activate. In the simulation, we see the opposite, primarily because of the more accurately reflected ANA behavior.
c-d) The analytical model predicts that both increasing the ANA concentration and decreasing the enzyme concentration will decrease the fraction cleaved during the exponential phase. The simulated data shows this effect for the enzyme concentration but not for the ANA, and the overall effect is minor.
e-f) The analytical model predicts increasing linear cleavage rate with increasing ANA concentration and enzyme concentration in the range plotted here. This is generally observed in the simulated data. However, the analytical model overestimates the cleavage rate by more than ten times under these conditions.

Experiment 4:

Again, we can replicate this experiment with an idealized ANA (Figure SN10-5).
a-b) The predictions for the time of the exponential phase appear nearly identical, although the magnitude is different.
c-d) The simulated data for the cleaved fraction much more closely resembles the analytical model. However, the magnitude of the effect is reduced in the simulated data.
e-f) The cleavage rate data remains similar to before. While the general trend is predicted, the magnitude is overestimated several times in the analytical model.
As before, this demonstrates that the simplified ANA behavior is a significant source of the difference between the analytical model and full numerical model. However, other discrepancies result in a significant difference even for the idealized ANA. When possible, the full numerical model should be preferred over the analytical model.

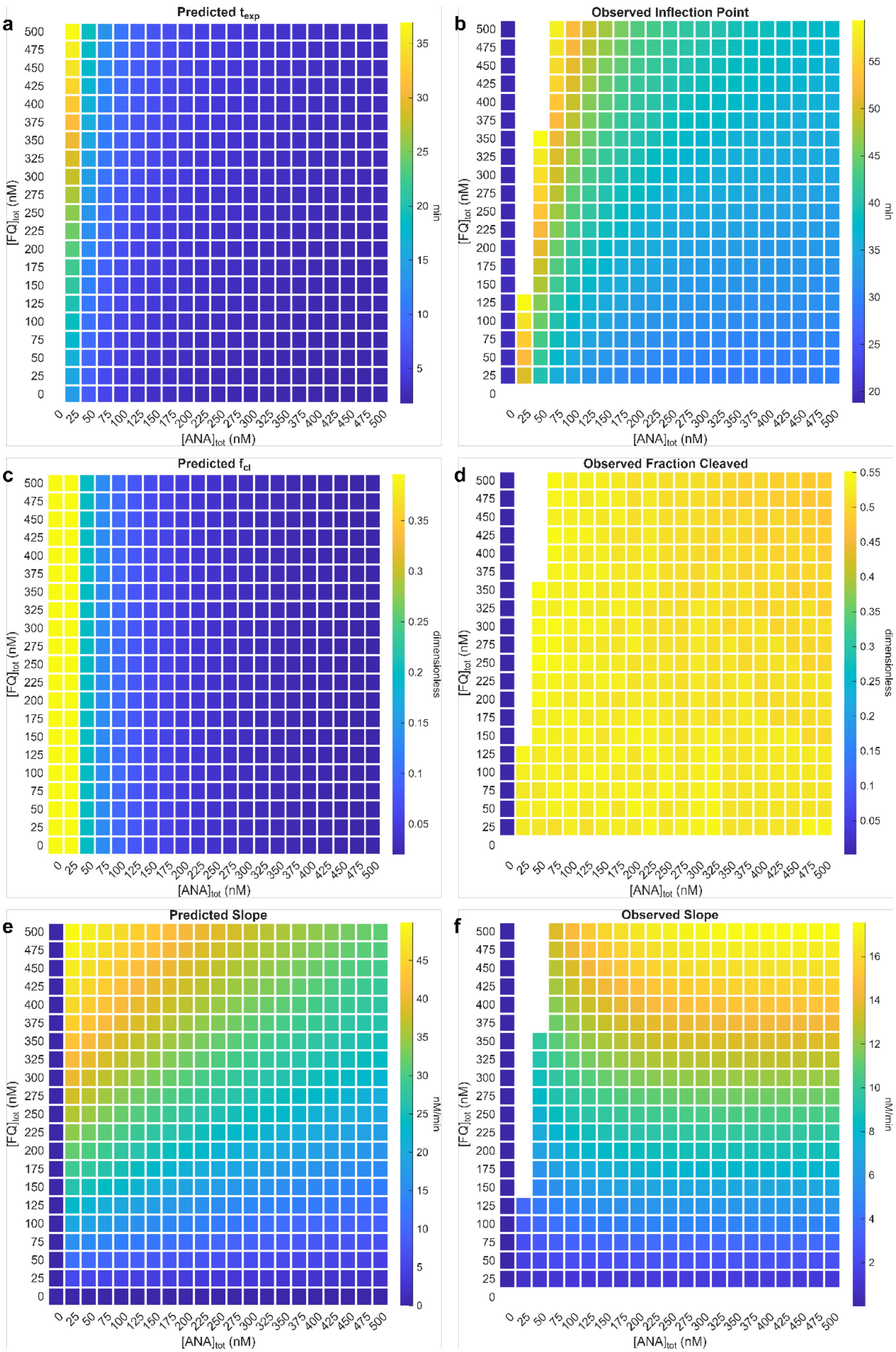


Figure SN10-2. Testing analytical simplifications for a typical ANA ($\Delta G^{\circ}_{uncleaved}$ = -13, $\Delta G^{\circ}_{cleaved}$ = -17) with $[E2] \approx 10$ nM.

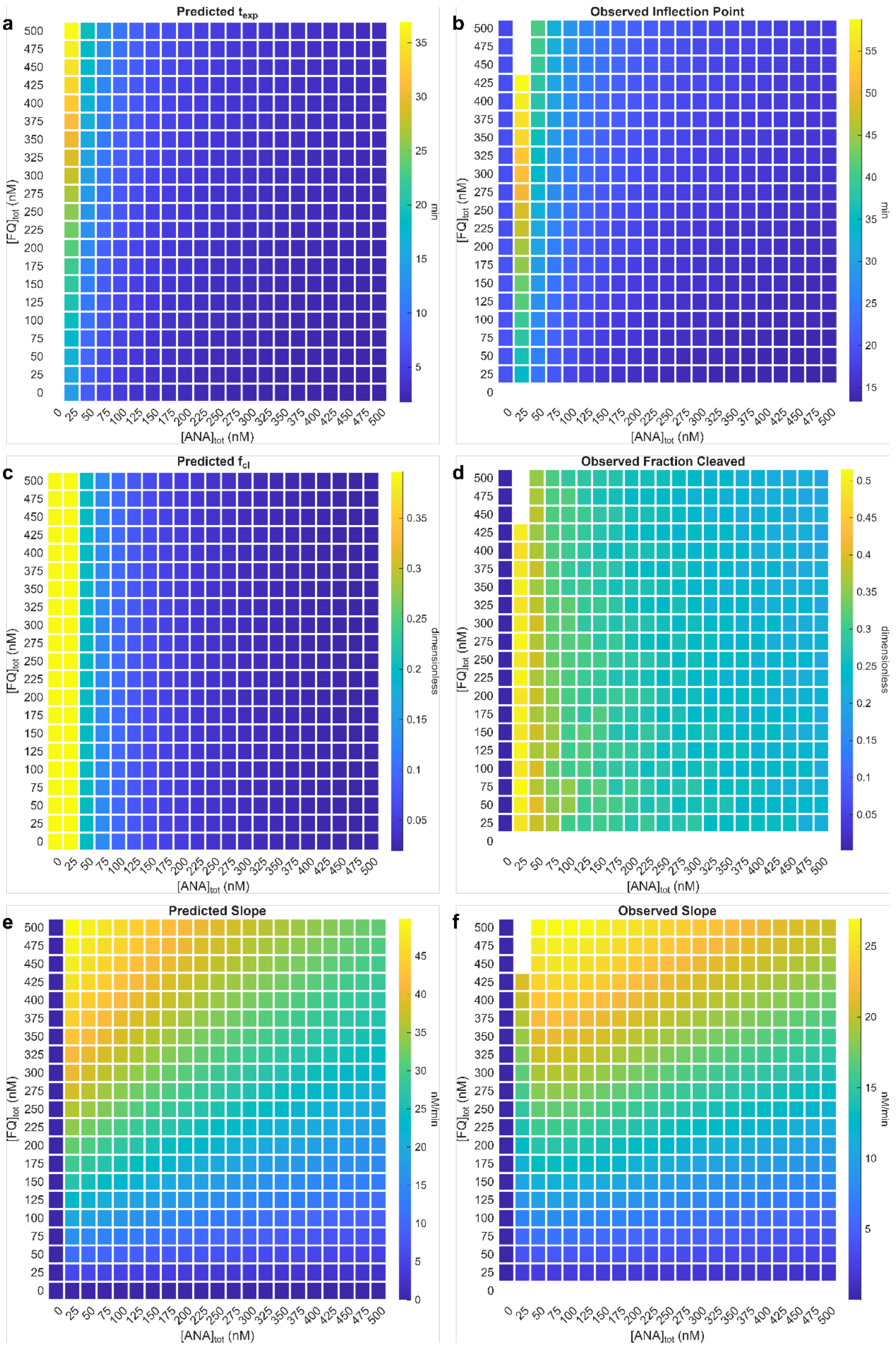


Figure SN10-3. Testing analytical simplifications for an idealized ANA ($\Delta G^{\circ}_{uncleaved}$ = -30, $\Delta G^{\circ}_{cleaved}$ = 100) with $[E2] \approx$ 10 nM.

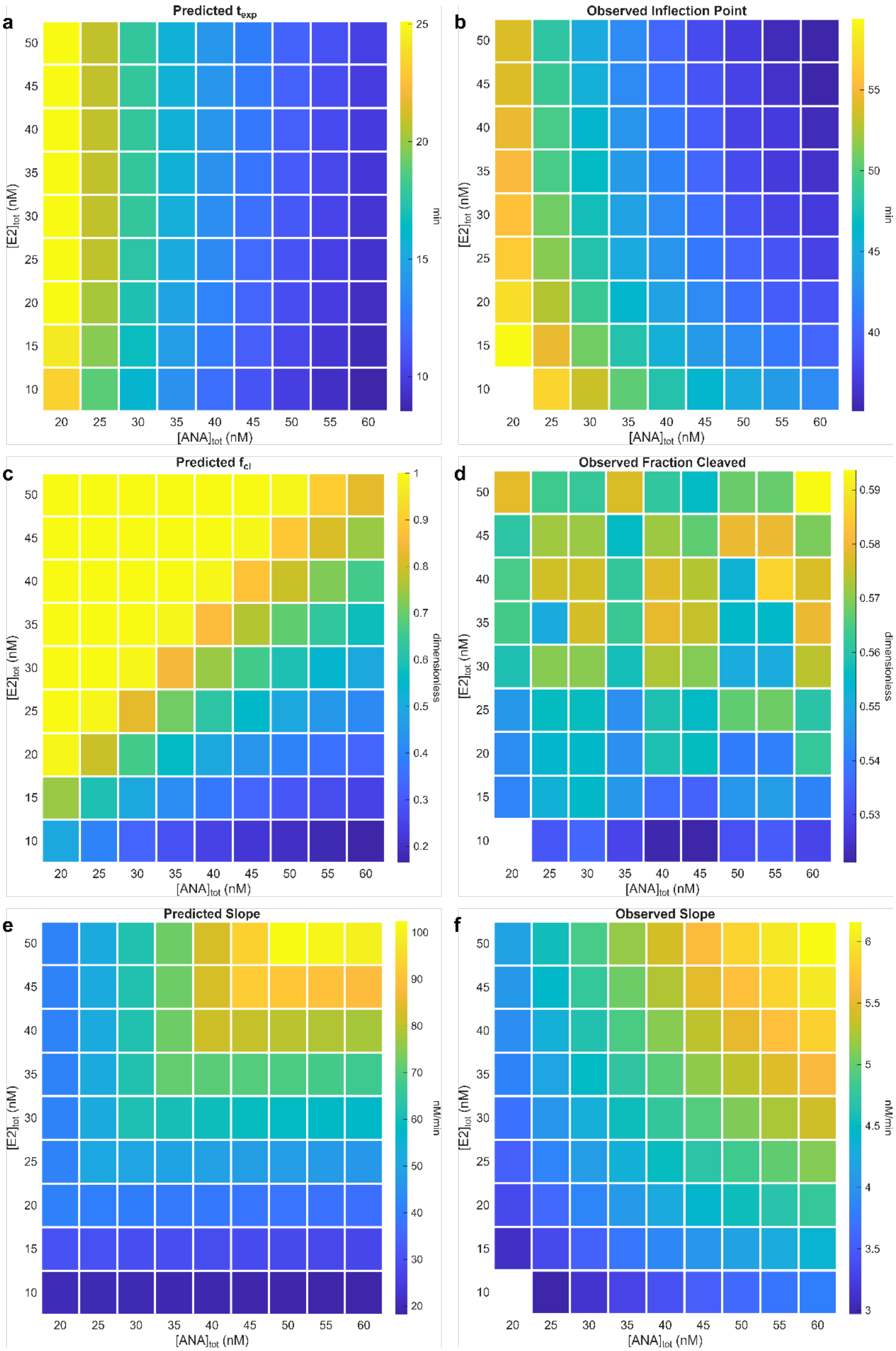


Figure SN10-4. Testing analytical simplifications for a typical ANA ($\Delta G^{\circ}_{uncleaved}$ = -13, $\Delta G^{\circ}_{cleaved}$ = -17) with $[FQ] \approx$ 100 nM.

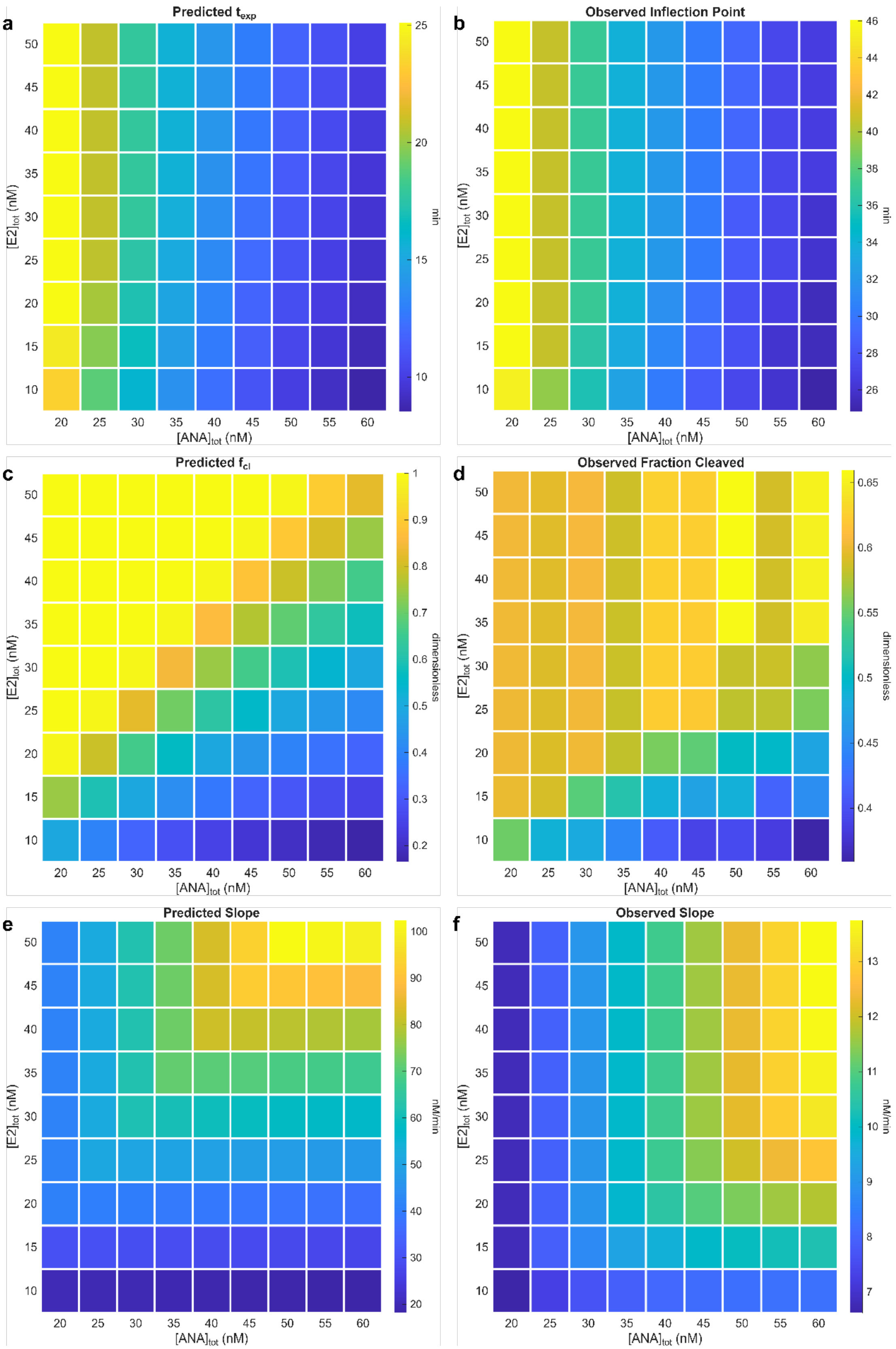


Figure SN10-5. Testing analytical simplifications for an idealized ANA ($\Delta G^{\circ}_{uncleaved}$ = -30, $\Delta G^{\circ}_{cleaved}$ = 100) with $[FQ] \approx$ 100 nM.

#### Supplementary Note 11: Data Calibration and Normalization Protocol

**Calibration**

The calibration procedure used here is based on the procedure described by Ramachandran and Santiago.[16] First, calibration curves were obtained. Briefly, the Cas12 enzyme (Alt-R LbCas12a (Cpf1) Ultra, obtained from IDT) pre-complexed with gRNA at a 5:3.3 ratio and diluted to a final enzyme concentration of 10 nM was activated with a final target concentration of 100 nM. This was used to cleave reporter (1 μM) for 12 hours at 37°C. The reporter used in all experiments is /56-FAM/TT TTT T/3IABkFQ/, synthesized by Integrated DNA Technologies (IDT). The uncleaved and cleaved reporter stocks were diluted into concentrations of 1000, 500, 250, 125, and 62.5 nM and split into three 10 μL replicates for each concentration. The fluorescent signal was measured for each sample in a QuantStudio3 Real-Time PCR System (ThermoFisher Scientific) for 10 minutes and the average signal was taken. This was used to produce linear fits (Figure SN11-1). This results in the following equation:

$$F(t) = m_{ucl}c_{ucl} + m_{cl}c_{cl}$$

Based on a total reporter concentration $[FQ]_{tot}$:

$$F(t) = m_{ucl}([FQ]_{tot} - c_{cl}) + m_{cl}c_{cl}$$

and:

$$c_{cl} = \frac{F(t) - m_{ucl}[FQ]_{tot}}{m_{cl} - m_{ucl}}$$

This allows for conversion from raw fluorescence to reporter concentration in quantitative experiments. The same machine was used for all subsequent experiments.

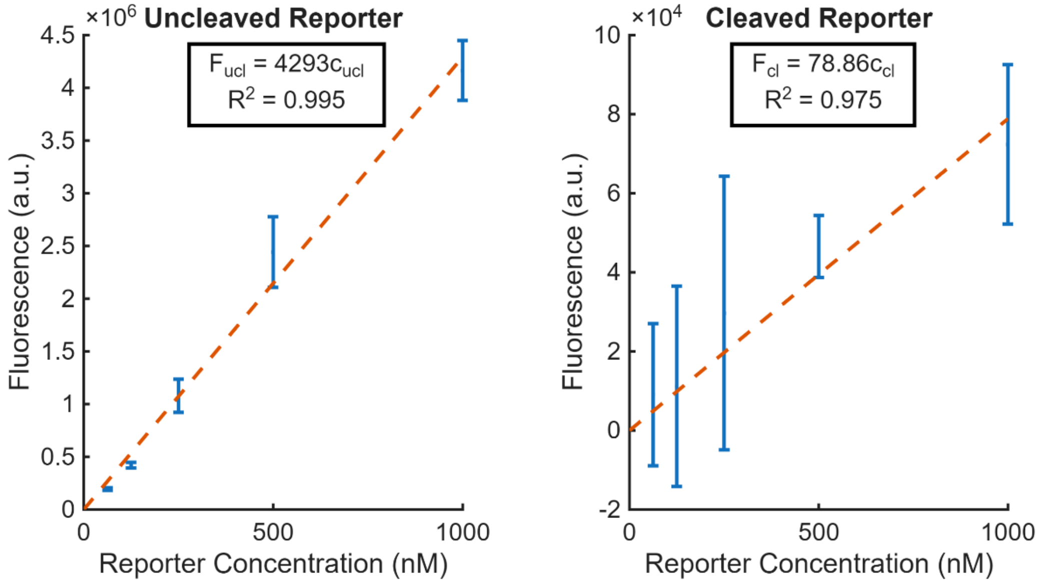


Figure SN11-1. Reporter calibration curves. Linear fits are generated for serial dilutions of uncleaved and cleaved reporter to allow for conversion from raw fluorescence to reporter concentration.

**Normalization**

It is necessary to correct for well-to-well fluorescence variation in fluorescence readers for quantitative experiments. Avaro and Santiago previously recommended a flat-field correction method that involves calibrating from a baseline measurement taken for each well before conducting the quantitative experiment.[25] Instead of this method, we have chosen to integrate a passive reference dye (ROX Reference Dye, 12223012 from ThermoFisher Scientific) in our assays. This is added during the same step when the reporter is added to the reaction mix at a final concentration of 500 nM. The primary benefit is that this repeats the calibration measurement with each experiment, correcting for variation in the readings over time. However, because the passive reference dye concentration is constant in the reaction mix, this method also has the potential to compensate for pipetting errors that affect the distribution of reaction mix. During each experiment, fluorescent measurements in the main channel are taken in parallel with a measurement of the passive reference. The normalization procedure is demonstrated in Figure SN11-2.

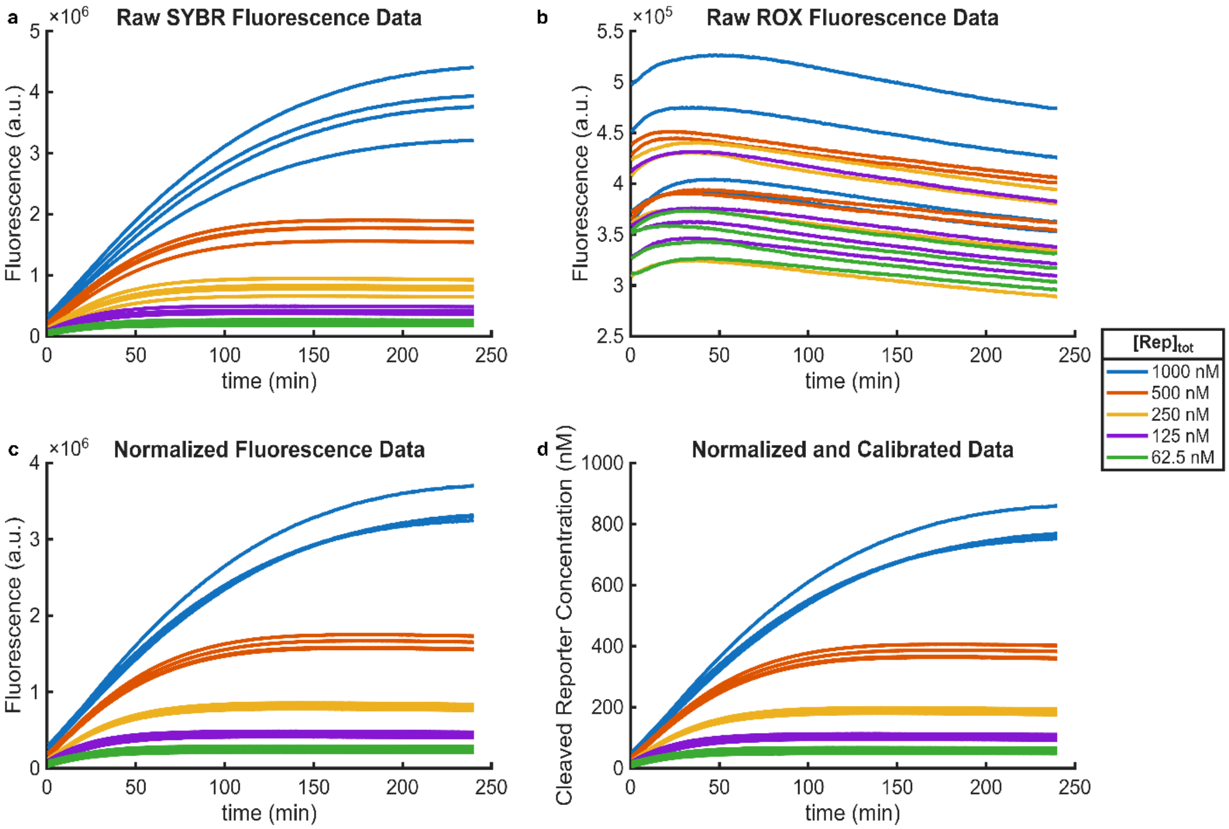


Figure SN11-2. Normalization using a passive reference dye. a) Raw SYBR fluorescence data. b) Raw ROX fluorescence data. c) Normalized fluorescence data using the normalization approach can be observed to reduce the variation between samples in the same group. d) Calibration is applied to convert the fluorescence signal to cleaved reporter concentration.

Figure SN11-2a and SN11-2b show the raw SYBR and ROX fluorescence data, respectively. The last 10 minutes of ROX data are averaged to find a baseline ROX value ($ROX_{well}$) for each sample well. These well values are averaged to obtain an experiment average ($\overline{ROX}$). The normalized signal is then calculated (Figure SN11-2c):

$$F'_{well}(t) = F_{well}(t)\frac{\overline{ROX}}{ROX_{well}}$$

This normalizes the fluorescence such that a well with ROX data matching the average passive reference value maintains the raw fluorescence value. In a well with increased ROX signal from this baseline, the raw fluorescence is decreased proportionally. In practice, this method is observed to reduce the variation in fluorescence within sample groups. After normalization, the calibration procedure is applied to obtain a concentration value (Figure S11-2d). Both of these procedures are implemented with MATLAB with helper functions in the DataAnalysis.m file included with the raw data from the experiments.

#### Supplementary Tables

#### Supplementary Table 1: CRISPR-Cas-based Autocatalytic Literature

| Publication | DOI | Journal | Authors | Publication Month & Year | ANA Description | CRISPR only? |
|---|---|---|---|---|---|---|
| A CRISPR-Cas autocatalysis-driven feedback amplification network for supersensitive DNA diagnostics (CONAN) | 10.1126/sciadv.abc7802 | *Sci. Adv.* | Shi et al. | Jan 2021 | Switchable caged gRNA | Yes |
| An aM-level sensitive cascade CRISPR-Dx system (ASCas) for rapid detection of RNA without pre-amplification | 10.1016/j.bios.2023.115248 | *Biosens. Bioelectron.* | Zhang et al. | Jun 2023 | Bulge DNA/RNA cascade probe | Yes |
| Asymmetric CRISPR enabling cascade signal amplification for nucleic acid detection by competitive crRNA | 10.1038/s41467-023-43389-7 | *Nat. Commun.* | Moon and Liu | Nov 2023 | Full/split crRNA competition mechanism | Yes |
| Topological barrier to Cas12a activation by circular DNA nanostructures facilitates autocatalysis and transforms DNA/RNA sensing (AutoCAR) | 10.1038/s41467-024-46001-8 | *Nat. Commun.* | Deng et al. | Mar 2024 | Circularized dsDNA | Yes† |
| An autocatalytic CRISPR-Cas amplification effect propelled by the LNA-modified split activators for DNA sensing | 10.1093/nar/gkae176 | *Nucleic Acids Res.* | Sun et al. | Apr 2024 | Split activator – hairpin and ssDNA with site-directed cleavage | Yes |
| A TdT-driven amplification loop increases CRISPR-Cas12a DNA detection levels | 10.1016/j.bios.2024.116464 | *Biosens. Bioelectron.* | Zwerus et al. | Oct 2024 | TdT-modifiable ssDNA | No |
| Preamplification-free ultra-fast and ultra-sensitive point-of-care testing via LwaCas13a (PASSPORT) | 10.1016/j.bios.2024.116400 | *Biosens. Bioelectron.* | Zeng et al. | Sep 2024 | U-rich single-loop hairpin | Yes |
| Amplification-free, OR-gated CRISPR-Cascade reaction for pathogen detection in blood samples | 10.1073/pnas.2420166122 | *PNAS* | Lim et al. | Mar 2025 | Dual-loop hairpin | Yes |
| Ultrasensitive detection of clinical pathogens through a target-amplification-free collateral-cleavage-enhancing CRISPR-CasΦ tool | 10.1038/s41467-025-59219-x | *Nat. Commun.* | Chen et al. | Apr 2025 | Dual-loop hairpin | Yes |
| CRISPR anti-tag-mediated room-temperature RNA detection using CRISPR/Cas13a | 10.1038/s41467-025-64205-4 | *Nat. Commun.* | Moon et al. | Oct 2025 | Anti-tag single-loop hairpin | Yes |
| Cascade CRISPR/cas Enables More Sensitive Detection of Toxoplasma gondii and Listeria monocytogenes than Single CRISPR/cas | 10.3390/microorganisms13081896 | *Microorganisms* | Chen et al. | Aug 2025 | Single-loop hairpin | Yes |
| Autocatalytic Cas13a biosensor enabled by RNA-nanocircles for ultrasensitive RNA detection | 10.1038/s44328-025-00067-6 | *npj Biosens.* | Deng et al. | Jan 2026 | RNA nanocircle | Yes |
| Dumbbell-shaped DNA topology drives self-sustaining CRISPR/Cas12a exponential amplification for ultrasensitive monitoring of DNA methyltransferase activity | 10.1016/j.talanta.2025.128841 | *Talanta* | Xiang et al. | Feb 2026 | DNA dumbbell with methylation-mediated cleavage | No |
| Bulge DNA-driven CRISPR/Cas12a dynamic activation circuit enables highly sensitive and versatile biosensing | 10.1016/j.bios.2026.118412 | *Biosens. Bioelectron.* | Wei et al. | April 2026 | Bulge DNA | No |

†Additional enzyme required for generation of ANA but not during assay

#### Supplementary Table 1: CRISPR-Cas-based Autocatalytic Literature (Continued)

| Publication | Limit of detection | Reaction time | Reaction volume (µL) | Temperature (°C) | Cas Subspecies | Cas Enzyme Concentration (nM) | ANA Concentration (nM) | Reporter Concentration (nM) |
|---|---|---|---|---|---|---|---|---|
| A CRISPR-Cas autocatalysis-driven feedback amplification network for supersensitive DNA diagnostics (CONAN) | 5 aM§ | 4 hours | 20 | 37 | LbCas12a | 1000 | 1000 | 1000 (Integrated w/ ANA) |
| An aM-level sensitive cascade CRISPR-Dx system (ASCas) for rapid detection of RNA without pre-amplification | 1 aM | 20 min | 25 | 33 | LwaCas13a & LbCas12a | 10 & 10# | 100 | 1000 |
| Asymmetric CRISPR enabling cascade signal amplification for nucleic acid detection by competitive crRNA | 100 fM | 1 hour | 20 | 37 | LbCas12a | 100 | 5-40* | 500 |
| Topological barrier to Cas12a activation by circular DNA nanostructures facilitates autocatalysis and transforms DNA/RNA sensing (AutoCAR) | 1 aM§ | 15 min | 100 | RT | LbCas12a | ~17 | ~173 | ~173 (Integrated w/ ANA) |
| An autocatalytic CRISPR-Cas amplification effect propelled by the LNA-modified split activators for DNA sensing | 4.7 fM§ | 50 min | 20 | 37 | LbCas12a | 50 | 50 & 200* | 200 (Integrated w/ ANA) |
| A TdT-driven amplification loop increases CRISPR-Cas12a DNA detection levels | 1 pM | 1 hour | 25 | 37 | LbCas12a | 50 | 25* | 400 |
| Preamplification-free ultra-fast and ultra-sensitive point-of-care testing via LwaCas13a (PASSPORT) | ~0.17 aM§ (100 copies/mL) | 5 min | 50 | 37 | LwaCas13a | 252 | 200 | 500 |
| Amplification-free, OR-gated CRISPR-Cascade reaction for pathogen detection in blood samples | 1 aM§ | 10 min | 11 | 33 | LbCas12a | 3.3 & 5.5# | 100 | 500 |
| Ultrasensitive detection of clinical pathogens through a target-amplification-free collateral-cleavage-enhancing CRISPR-CasΦ tool | 0.18 aM§ | 30-60 min | 100 | 37 | CasΦ (Cas12j) | 30 | 10 | 250 |
| CRISPR anti-tag-mediated room-temperature RNA detection using CRISPR/Cas13a | 10 am§ | 60 min | 20 | RT | LwaCas13a | 10 | 1 & 5 | 500 |
| Cascade CRISPR/cas Enables More Sensitive Detection of Toxoplasma gondii and Listeria monocytogenes than Single CRISPR/cas | 10 fM§ | 30 min | 20 | 30 | AsCas12a | 25 & 25# | 50 | 500 |
| Autocatalytic Cas13a biosensor enabled by RNA-nanocircles for ultrasensitive RNA detection | 1 aM | 15 min | 100 | 37 | LwaCas13a | 40 | 120 | 120 (Integrated w/ ANA) |
| Dumbbell-shaped DNA topology drives self-sustaining CRISPR/Cas12a exponential amplification for ultrasensitive monitoring of DNA methyltransferase activity | $6.37 \times 10^{-4}$ U/mL‡ | 60 min | 200 | 37 | LbCas12a | 150 | 200* | 200 |
| Bulge DNA-driven CRISPR/Cas12a dynamic activation circuit enables highly sensitive and versatile biosensing | 14 CFU/mL | 1 hr 5 min | 50 | 37, 25 | LbCas12a | 200 | 100 | 5000 |

§Multiple reaction formats – values are chosen for basic autocatalytic format with lowest LOD

#Assay pre-complexes target-detection and signal-amplification Cas enzymes separately.

*Multiple ANA components

‡Assay measures enzyme activity rather than concentration directly

### Supplementary Table 2: Nucleic Acid Sequences and Free Energies

| Strand Name | | Sequence (5'→3', Target Region Underlined) | Length (nt) | ΔG°* (kcal/mol) |
|---|---|---|---|---|
| Complex ANA (Uncleaved) | | GGATATACTTTTTATTTTTGATAAATATATATATTTTTTATTTTTATATATATATATCGTATATCC | 65 | -4.48 |
| Complex ANA (Cleaved at bulge) | Strand A | GGATATACTTTTT | 13 | 0.00 |
| | Strand BC | ATTTTTGATAAATATATATATTTTTATTTTTATATATATATATCGTATATCC | 52 | -3.37 |
| Complex ANA (Cleaved at bulge and hairpin) | Strand A | GGATATACTTTTT | 13 | 0.00 |
| | Strand B | ATTTTTGATAAATATATATATTTTTAT | 27 | 0.00 |
| | Strand C | TTTTATATATATATATCGTATATCC | 25 | -0.78 |
| Complex ANA (Cleaved at hairpin) | Strand AB | GGATATACTTTTTATTTTTGATAAATATATATATTTTTAT | 40 | -1.14 |
| | Strand C | TTTTATATATATATATCGTATATCC | 25 | -0.78 |
| Hairpin10 (Uncleaved) | | CGGCGGATATACGATATATATATATAATAATTAATATATATATATATCGTATATCCGCCG | 60 | -21.68 |
| Hairpin10 (Cleaved) | Target | TTAATATATATATATATCGTATATCCGCCG | 30 | -26.99 |
| | Complement | CGGCGGATATACGATATATATATATAATAA | 30 | |
| HairpinOpt (Uncleaved) | | GGATATACGATATATATATATAATAATTAATATATATATATATCGTATATCC | 52 | -13.70 |
| HairpinOpt (Cleaved) | Target | TTAATATATATATATATCGTATATCC | 26 | -18.46 |
| | Complement | GGATATACGATATATATATATAATAA | 26 | |
| Bulge6 (Uncleaved) | Target | AGGGGCGTGCATATATATATATCGTATATCCGTTGGTGACG | 41 | -42.91 |
| | Complement | CGTCACCAACGGATATACTTGTGTGATATATATATATGCACGCCCCT | 47 | |
| Bulge6 (Cleaved) | Target | AGGGGCGTGCATATATATATATCGTATATCCGTTGGTGACG | 41 | -48.63 |
| | Complement 1 | CGTCACCAACGGATATACTTG | 21 | |
| | Complement 2 | TGTGATATATATATATGCACGCCCCT | 26 | |
| Bulge11 (Unleaved) | Target | AGGGGCGTGCATATATATATATCGTATATCCGTTGGTGACG | 41 | -42.31 |
| | Complement | CGTCACCAACGGATATACATCTATGTGTTGATATATATATATGCACGCCCCT | 52 | |
| Bulge11 (Cleaved) | Target | AGGGGCGTGCATATATATATATCGTATATCCGTTGGTGACG | 41 | -48.81 |
| | Complement 1 | CGTCACCAACGGATATACATCTA | 23 | |
| | Complement 2 | TGTGTTGATATATATATATGCACGCCCCT | 29 | |

*Minimum free energy structure, simulated in NUPACK (material = DNA, Temperature = 37°C, model = dna04.2, stacking = all stacking, Na = 0.05 M, Mg = 0.01 M, max complex size = 4).[1,2]
